# MultiPopPred: A Trans-Ethnic Disease Risk Prediction Method, and its Application to the South Asian Population

**DOI:** 10.1101/2024.11.26.625410

**Authors:** Ritwiz Kamal, Manikandan Narayanan

## Abstract

Genome-wide association studies (GWAS) have guided significant contributions towards identifying disease associated Single Nucleotide Polymorphisms (SNPs) in Caucasian populations, albeit with limited focus on other understudied low-resource non-Caucasian populations. There have been active efforts over the years to understand and exploit the population specific versus shared aspects of the genotype-phenotype relation across different populations or ethnicities to bridge this gap. However, the efficacy of transfer learning models that are simpler than existing approaches and utilize individual-level data remains an open question. We propose MultiPopPred, a novel and simple trans-ethnic polygenic risk score (PRS) estimation method that taps into the shared genetic risk across populations and transfers information learned from multiple well-studied auxiliary populations to a less-studied target population. The default version of MultiPopPred (MPP-PRS+) harnesses individual-level data using a specially designed Nesterov-smoothed penalized shrinkage model and an L-BFGS optimization routine. Extensive comparative analyses performed on simulated genotype-phenotype data, assuming an infinitesimal/omnigenic model, reveal that MPP-PRS+ improves PRS prediction in the South Asian population by 38% on average across all simulation settings when compared to state-of-the-art trans-ethnic PRS estimation methods including SBayesRC-Multi and PROSPER. This improvement is enhanced in settings with low target sample sizes and in semi-simulated settings. These predictive advantages are further echoed in real-world evaluations against 16 UK Biobank quantitative and binary traits. For example, compared to SBayesRC-Multi, MPP-PRS+ achieves comparable performance (within 5% of SOTA performance) on 3 traits, superior performance on 7 other traits, and lags behind on 4 (lipid-related) traits. Neither method could meaningfully predict the remaining 2 traits. MPP-PRS+ maintains a similarly competitive, albeit distinct, performance trend compared to PROSPER. These performance trends are promising and encourage application of MultiPopPred for reliable PRS estimation in low-resource populations with individual-level data for complex omnigenic traits.

## 1 Introduction

Humans across the world are known to have differences in their genetic makeup alongside various environmental factors; these differences in genetic and environmental factors can translate to different populations or ethnicities having varying degrees of predisposition to complex diseases [1–3]. Complex (polygenic) diseases usually manifest as a consequence of a large number of common genetic factors (single-nucleotide polymorphisms or SNPs) contributing in minuscule to moderate amounts simultaneously to the disease trait [4, 5]. Typical examples of such diseases include type 2 diabetes, cardiovascular diseases, and neurological disorders. For these diseases, studies have reported that people of South Asian ancestry exhibit different risk profiles compared to Caucasian individuals — for instance, South Asians exhibit a higher risk for cardiovascular diseases [6] and a higher prevalence of diabetes [7] as compared to Caucasians.

Traditionally, genome-wide association studies (GWAS), employing linear regression models, have served as a popular and established means to study associations between genetic factors or SNPs and traits of interest. Studies with sufficiently large sample sizes may be used to construct polygenic risk scores (PRS), which are single-valued estimates aggregating the effects of SNPs across the human genome to provide insights into an individual’s predisposition to develop a given trait [8]. While large GWAS (with ≥ 10^6^ individuals in the study) conducted on Caucasian (mainly European) populations have been significantly insightful [9, 10], there is a clear lack of representation of non-Caucasian populations (such as the South Asian population) in the field. GWAS conducted on South Asian populations usually have around a few hundred to a few thousand individuals only in the study [11–14], thereby rendering these studies statistically underpowered to produce estimates reliable enough to compute PRS suitable for downstream utility. Consequently, PRS estimation studies [15, 16] conducted on South Asian populations also suffer from a similar deficit of samples. Moreover, owing to differences in linkage disequilibrium (LD) patterns, allele frequencies, and varying trait heritability across populations, estimates from European GWAS cannot simply be plugged into a South Asian population [17, 18], without risking exacerbation of health disparities [19]. There are active efforts underway to address this under-representation of low-resource non-Caucasian populations, such as the South Asian population.

A seemingly straightforward solution to the above problem would be to collect more data on South Asian individuals before conducting GWAS, albeit at an unnecessarily higher cost of time, resources, and effort. Recent efforts (such as SBayesRC-Multi [20], PROSPER [21], Poly-Pred+ [22], PRS-CSx [23], and TL-Multi [24]) instead aim to understand and harness the differences and commonalities in SNP effect sizes (or risk towards a disease) between populations or ethnicities. Subsequently, they aim to utilize this information on population-specific versus shared genetic disease risk to perform transfer learning from well-studied, informative auxiliary populations (such as European populations) to a less-studied target population (such as the South Asian population). Kachuri et al. [25] present a comprehensive review of other similar published works in this domain. While each pre-existing method (including SBayesRC-Multi [20], PROSPER [21], PRS-CSx [23], TL-Multi [24], and Lassosum [26, 27]) adds an important new perspective to the problem at hand, they leave ample scope to integrate information from multiple auxiliary populations in a manner that allows for accurate estimation of SNP effect sizes, and consequently, disease risk in the target population. For example, while several existing methods estimate PRS using GWAS summary statistics and external LD (from a closely matched cohort) due to ease of accessibility, the potential hosted in using individual-level data to capture true LD — especially for low resource populations — remains an important open question. This positioning of existing methods to use summary-rather than individual-level data often leads to complex methodological frameworks, leaving room for a simpler approach without compromising performance. As also observed before [21, 25], there continues to be a strong need for complementary trans-ethnic methods to estimate disease risk, because no single existing method unanimously outperforms all other methods in estimating the risk of diseases across diverse settings. Consequently, there is also a pressing need for best practices and recommendations on which methods to use among the many available in the literature, depending on differences in genetic architectures of traits, population structures, and sample sizes.

We propose MultiPopPred, a novel and simple trans-ethnic PRS estimation method that simultaneously integrates information on SNP effect sizes learned from multiple auxiliary populations (such as Caucasian populations) and transfers the same to a target population. MultiPopPred employs a specially designed Nesterov-smoothed penalized shrinkage model and an L-BFGS optimization routine. We present MPP-PRS+ as the default version of MultiPopPred due to its utilization of individual-level data, and also present alongside four other versions that can adapt to different degrees of availability of individual-level versus summary-level data and work with weights assigned to auxiliary populations. Extensive comparative analyses performed on simulated genotype-phenotype data, assuming an infinitesimal model for both quantitative and binary phenotypes, reveal that MPP-PRS+ improves PRS prediction on average in the South Asian population by 38% overall across all simulation settings and 91% across simulation settings with low target sample sizes when compared with state-of-the-art (SOTA) methods. Specifically, MPP-PRS+ reported an average performance gain of 32% (27%, 17%, and 32% against SBayesRC-Multi, PROSPER, and PRS-CSx, respectively) across all simulation settings with quantitative phenotypes; noticeably higher gains of 127% (52%, 92%, and 159%, respectively) were observed across simulation settings with low target sample sizes. Similar average performance gains of 43% and 148% were observed across all simulated settings with binary phenotypes and semi-simulated settings with quantitative phenotypes, respectively.

When evaluating our method on 16 real-world quantitative and binary traits from UK Biobank, MPP-PRS+ demonstrated robust predictive advantages against SOTA methods. Specifically, compared to SBayesRC-Multi, MPP-PRS+ exhibited comparable performance (within 5% of SBayesRC-Multi’s performance) on 3 traits and a superior performance on 7 other traits. Conversely, SBayesRC-Multi proved more suitable for 4 quantitative lipid traits characterized by highly sparse architectures (High and Low Density Lipoprotein, Total Cholesterol and Triglycerides). Neither method could meaningfully predict the remaining 2 traits (Systolic and Diastolic Blood Pressure). Similarly with respect to PROSPER, MPP-PRS+ demonstrated comparable performance (within 5% of PROSPER’s performance) on 6 traits, superior performance on 6 traits, lagging behind on 2 traits while the 2 remaining traits remained unpredictable.

Ablation studies and sensitivity analyses on a subset of the benchmarks show that both our input choice to use individual-level data (and associated true LD information) and our methodological choice to use a simpler (L-BFGS optimized, Nesterov-smoothed, *L*_1_-penalized, linear regression model) play a role in our competitive performance. These performance trends are promising and encourage application of MultiPopPred towards reliable PRS estimation for complex omnigenic traits under resource-constrained real-world settings when individual-level data is available. For other cases, we provide a triage of recommendations on which methods to use depending on the type of data available and the underlying genetic architecture of the trait in consideration.

## 2 Results

### 2.1 Overview of MultiPopPred

MultiPopPred builds upon the advancements made by previous trans-ethnic PRS estimation methods, while overcoming their shortcomings, to provide improved effect-size estimates of SNPs for low-resource populations. MultiPopPred (Figure 1) employs an *L*_1_-penalized lasso regression approach to close the gap between a target population’s effect sizes and a set of aggregated effect sizes over multiple (*K* ≥ 1) auxiliary populations. By doing so, our method capitalizes on the shared genetic makeup across populations and aims to perform transfer learning rather than learning weights from scratch when there is a dearth of trainable data for the target population. Depending upon the availability of the quantity and type of auxiliary and target data (i.e., individual-level versus summary-level data) and the manner of quantification of similarity between a given pair of populations, we propose five versions of MultiPopPred. MultiPopPred-PRS+ (MPP-PRS+) serves as the default version of our method and assumes the availability of complete information for training, requiring individual-level data for the target as well as auxiliary populations to perform transfer learning from joint all-SNP auxiliary PRS models to a joint all-SNP target PRS model. We propose another formal PRS version, MultiPopPred-PRS (MPP-PRS), that requires only the target population’s individual-level data while using external LD reference panels for the auxiliary populations’ PRS computation. Next, we propose three heuristic PRS versions that perform transfer learning directly from per-SNP models (GWAS summary statistics) to a joint all-SNP PRS model. These include (i) MultiPopPred-GWAS (MPP-GWAS), (ii) MultiPopPred-GWAS-Admixture (MPP-GWAS-Admix), and (iii) MultiPopPred-GWAS-TarSS (MPP-GWAS-TarSS). MPP-GWAS and MPP-GWAS-Admix require individual-level target data for training along with summary-level auxiliary data. MPP-GWAS-TarSS mimics the more commonly occurring real-world scenario and operates only with the target and auxiliary summary statistics, along with an external LD reference panel for the target population. Except for MPP-GWAS-Admix, all four versions of MPP integrate information from multiple auxiliary populations by equally weighting each source of information. MPP-GWAS-Admix weighs each auxiliary population in proportion to its admixture component in target individuals. Further details can be found in Methods and Figure 1. All five versions of MultiPopPred employ a Nesterov-smoothed penalized shrinkage model and an L-BFGS optimizer for accurate and effective convergence to produce precise *β* estimates of SNPs. A comprehensive overview of all simulation and real-world benchmarks that we have used to evaluate our method against and performed comparative analysis has been detailed in Section 2.2 and Supplementary Data 1-2.

**Fig. 1.**
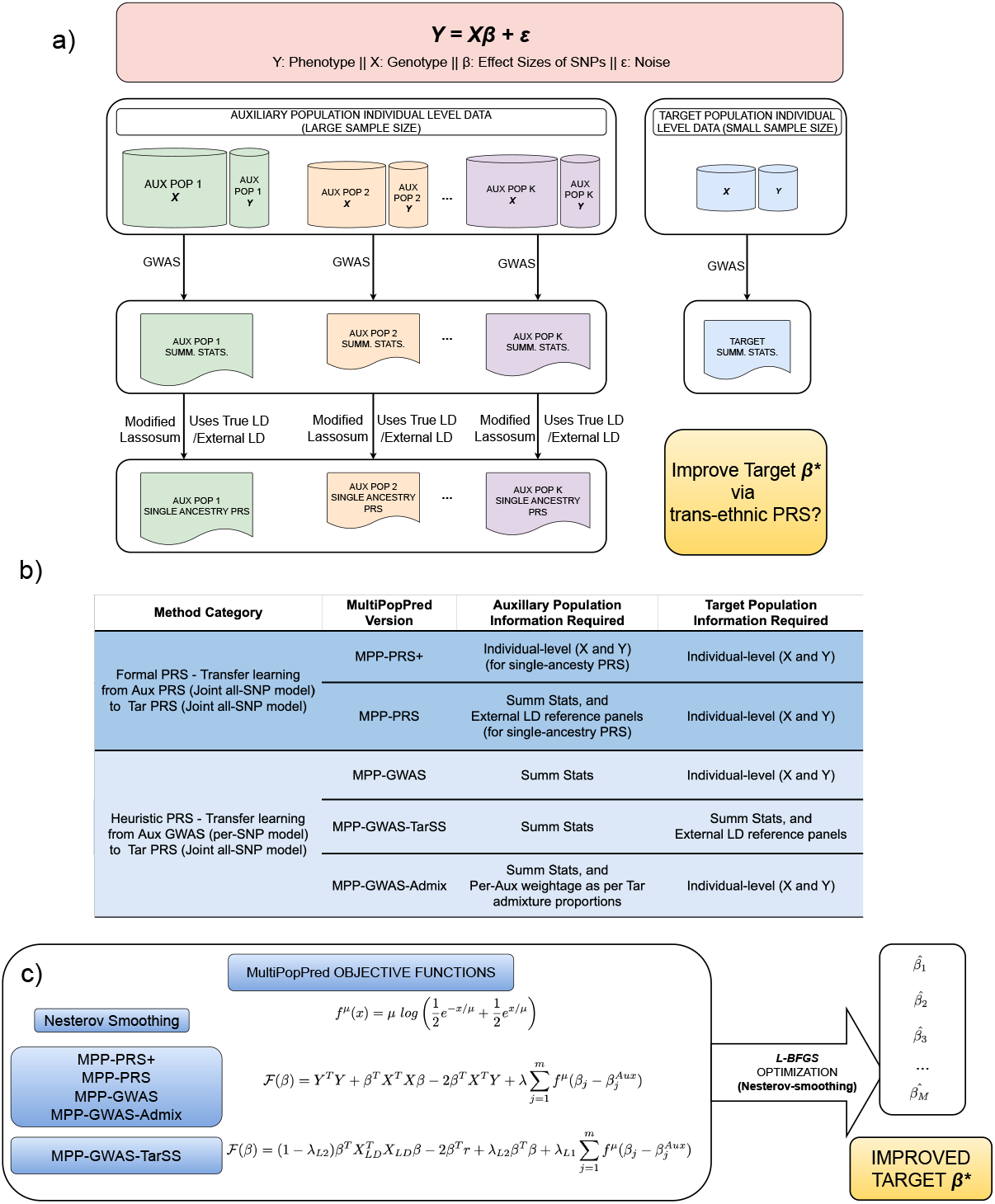
Overview of MultiPopPred Methodology: (a) GWAS conducted on auxiliary and target populations of varying sample sizes are shown. Summary statistics of a population with larger sample sizes are statistically more reliable. MultiPopPred aims to transfer effect sizes learned from auxiliary populations to the target population to improve prediction accuracy. (b) Five versions of MultiPopPred based on the availability of individual-level versus summary-level data and the weights of each auxiliary population. MPP-PRS+, our default version, harnesses individual-level data as shown. (c) MultiPopPred objective functions employed by the different versions for quantitative traits are shown here (see Methods for similar details for binary traits analyzed using a logistic module). Here *X, Y*, and *β* refer to the genotype, phenotype, and effect sizes of SNPs, respectively. For more details on the formulae, see Methods. Abbreviations: Aux: Auxiliary Population, Tar: Target Population, Summ. Stats.: Summary Statitics.

### 2.2 Overview of comparative benchmarks

We have performed a comprehensive comparative analysis of MultiPopPred with several established SOTA PRS estimation methods, which can be broadly categorized into three types depending on their input requirements. First, we have the single-population-based methods such as Baseline GWAS [28], SBayesRC [20] and Lassosum2 [21, 26]. Next, we have (one auxiliary, one target) population-based methods such as TL-Multi [24]. Finally, we have (multiple auxiliary, one target) population-based methods such as PRS-CSx [23], PROSPER [21], and SBayesRC-Multi [20] besides MultiPopPred. Brief descriptions of these methods and the hyperparameters used in each case can be found in Methods. We assess the accuracy of our method’s PRS estimates under several simulation, semi-simulation, and real-world settings, involving a range of sample sizes and heritability values (Sections 2.3, 2.4, and 2.5). Wherever applicable, like in any traditional machine learning setup, the available data has been split into training (to learn a predictive model), validation (to tune hyperparameters), and testing (to report performance on) sets. Hyperparameter tuning results for the *L*_1_ and *L*_2_ penalizations in the case of MultiPopPred can be found in Supplementary Table 1. We tuned MultiPopPred’s hyperparameters for a subset of simulated settings and froze the optimal hyperparameter values for the rest of the simulated settings (Supplementary Data 3). In the real-world setting, we tuned MultiPopPred’s hyper-parameters for each trait separately (Supplementary Data 3). We used 1000 Genomes Project (1KG) Phase 3 [29] defined super-populations (EUR, EAS, AMR, AFR, SAS) as reference panels wherever applicable (Methods).

### 2.3 Benchmarking using simulated data

We evaluated the performance of MultiPopPred and other SOTA methods on fully simulated genotype and phenotype data, with a major focus on quantitative traits and additional analyses on binary traits. Individual-level genotypes for each population were simulated using HAPGEN2 [30] and the corresponding per-population 1KG samples as reference (Methods). The ground truth effect sizes of SNPs were generated using a *K*-dimensional multivariate Gaussian distribution (*K* corresponding to the number of populations) with a fixed covariance matrix (Σ) (Methods). Subsequently, the phenotypes were generated in accordance with a linear additive model: *Y*_(*n×*1)_ = *X*_(*n×m*)_*β*_(*m×*1)_ + *ϵ*_(*n×*1)_ (Methods). We assumed each SNP to have some contribution towards the phenotype, no matter how minuscule, so as to resemble the real-world setting.

We designed three major analyses to understand the importance and optimality, with respect to real-world resemblance, of three major parameters that controlled the different simulation settings: (i) heritability (*h*^2^), (ii) number of target population samples, and (iii) the number of auxiliary population samples. Here, heritability (more precisely, narrow-sense heritability) *h*^2^ refers to the proportion of the variation in *Y* explained by the genetic component alone, i.e., 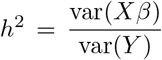. For each parameter configuration, the simulation was repeated 5 times, and for each simulated setting, MultiPopPred was run under 15 different configurations of auxiliary-target population tuples (Supplementary Data 4-6). Figure 2 depicts the comparative performances of the different PRS estimation methods for a subset of these configurations. Performances were evaluated on test datasets using the metric correlation ratio (see Methods).

**Fig. 2.**
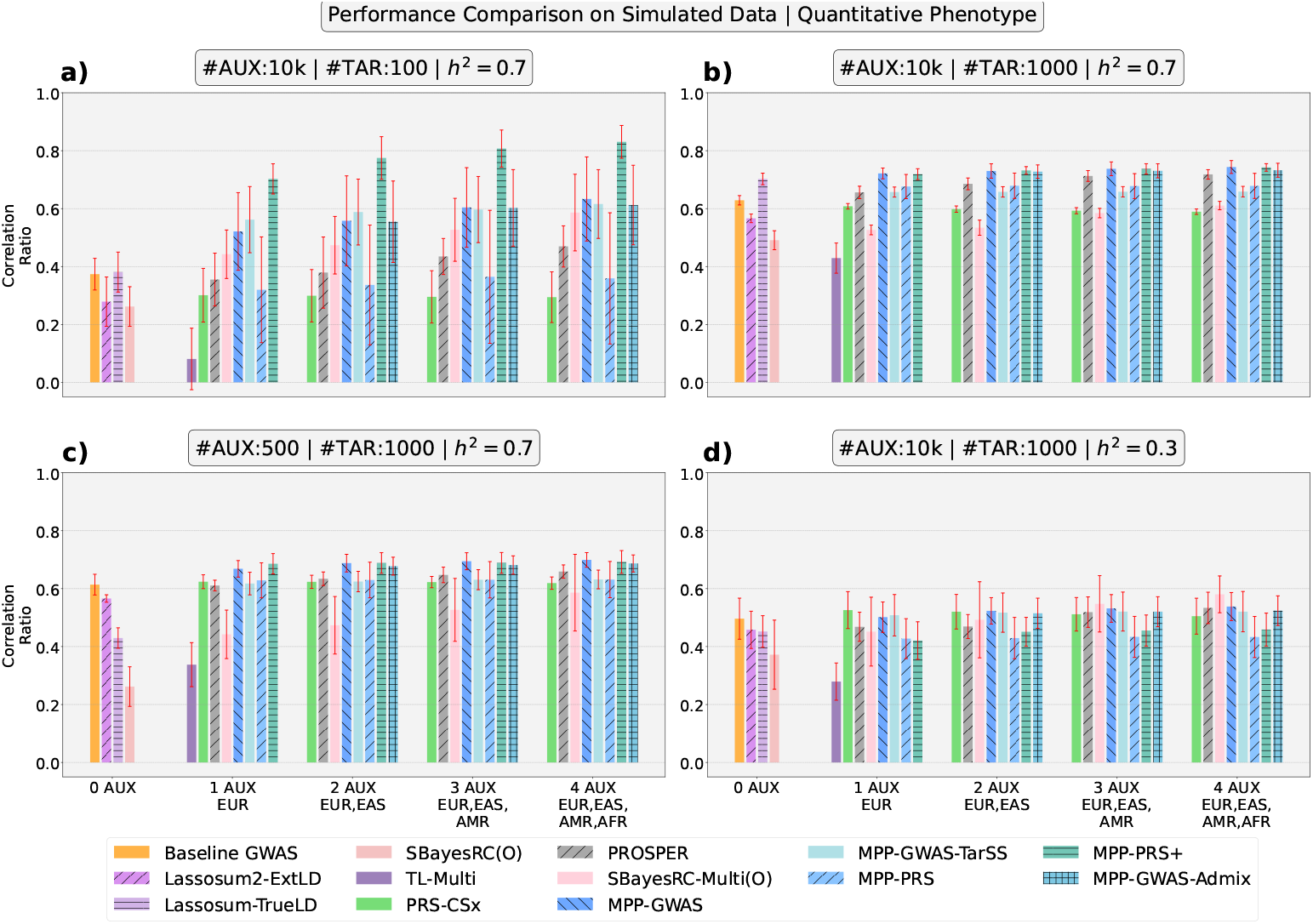
Performance Comparison on Simulated Data (Quantitative Phenotype Prediction): We depict the comparative performance of MultiPopPred with SOTA methods under four different simulation settings and four auxiliary-target population tuples. For all simulation settings, inter-population correlation was set to 0.8. (A) The number of samples per auxiliary population was set to 10,000, while the number of target population samples was kept very low, i.e., 100, to represent an extreme condition. *h*^2^ was set to 70%, akin to heritability values exhibited by most complex traits. (B-D) Subsequently, each of these simulation parameters was changed one at a time to values indicated atop the panels to observe trends in the comparative performance. Note that MPP-GWAS-Admix is not applicable under the single auxiliary population setting since calculating admixture proportions requires ≥ 2 auxiliary populations. On the other hand, TL-Multi is not applicable for *>* 1 auxiliary population settings. SBayesRC(O) and SBayesRC-Multi(O) refer to the “oracle” version of these two methods. Abbreviations: Aux: Auxiliary Population, Tar: Target Population, *h*^2^: Heritability, Correlation ratio: Correlation between the actual and predicted phenotypes, divided by the maximum achievable value of that correlation (higher value indicates better performance).

#### 2.3.1 Increase in the number of auxiliary populations leads to better target estimates

Before elaborating on the specific observations for each of the three analyses mentioned above, we would like to point out a general trend observed across all simulation settings and experiments: an increase in performance as the number of auxiliary populations used for transfer learning was increased from 1 to 4 (see Figure 2). The increase was subtle but consistent. The subtlety of the increase can be attributed to the way simulations were designed, such that the correlation between the ground truth effect sizes of SNPs for each pair of populations used in the study was fixed at a relatively high value of 0.8. This observation also validates our hypothesis that information from multiple auxiliary populations, if aggregated appropriately, lends to efficient transfer learning that in turn translates to improved PRS estimates for a target population.

#### 2.3.2 MultiPopPred performs on-par or better than SOTA methods at varying heritabilities

In our first major analysis, we fixed the number of auxiliary samples per population at 10,000 (mimicking the high resources available for well-studied populations), the number of target samples at 1000 (mimicking the typically available low resources for an understudied population) and varied the heritability (*h*^2^) from 10% to 100% in steps of 20 (*h*^2^ = 10% representing a phenotype having a much higher contribution from the noise component and *h*^2^ = 100% representing a phenotype having no contribution from the noise component; see Supplementary Data 4). We observed higher variability in performance at very low *h*^2^ values (Figure 2d), expectedly so, due to the true phenotype having very little contribution from the genetic component. As the *h*^2^ value and consequently the contribution of the genetic component to the phenotype increased, each method was observed to report an improvement in performance both in terms of the observed correlation ratio and the stability of the same. Moreover, at low *h*^2^ values (10%-30%), almost all methods were noted to have more or less comparable performances (Figure 2d) except for TL-Multi. At moderate *h*^2^ (50%-70%) and higher *h*^2^ (90%-100%) values, MultiPopPred (specifically the MPP-PRS+, MPP-GWAS, and MPP-GWAS-Admix versions) was observed to have a clear edge over other methods (Figures 2a, 2b, 2c).

Figure 3 depicts the overall improvements, across all simulation settings, in performance with respect to correlation ratio when the two best performing versions of MultiPopPred (MPP-PRS+ and MPP-GWAS) were compared against the three best performing and leading SOTA methods (SBayesRC-Multi, PROSPER and PRS-CSx). Averaging across the simulated scenarios (with quantitative phenotypes), MPP-PRS+ demonstrated 27%, 17% and 32% improvements over SBayesRC-Multi, PROSPER and PRS-CSx, respectively, while MPP-GWAS was observed to obtain a 48%, 13% and 26% improvement over SBayesRC-Multi, PROSPER and PRS-CSx, respectively. Further comparisons of the MultiPopPred versions with SOTA methods can be found in Supplementary Figure 1.

**Fig. 3.**
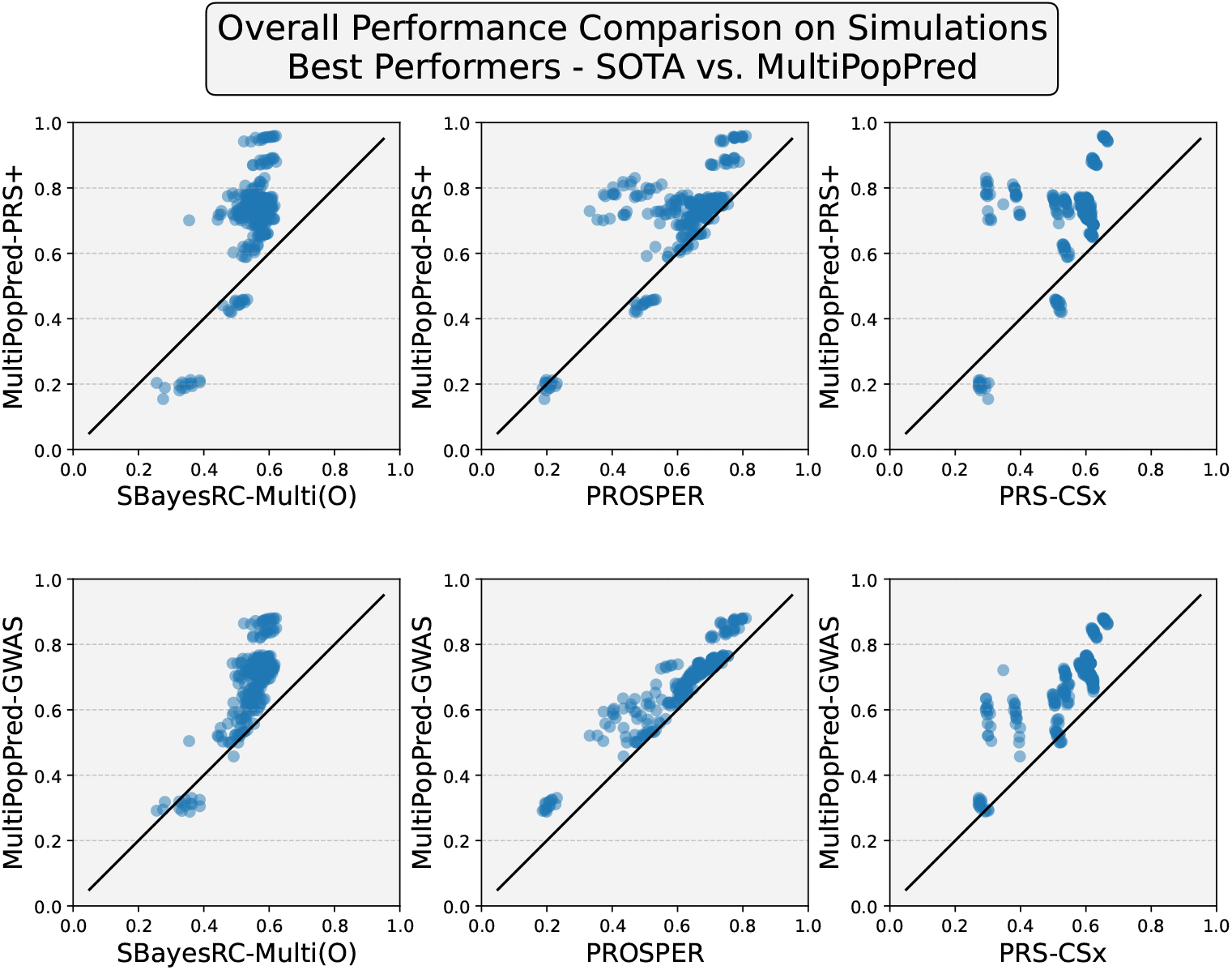
Overall Performance Comparison On All Simulation Scenarios Tested (with respect to Quantitative Phenotype Prediction using correlation ratio): We depict the comparative performance of the two best performing versions of MultiPopPred (MultiPopPred-PRS+ and MultiPopPred-GWAS) against the three best performing and leading SOTA methods (SBayesRC-Multi(O), PROSPER and PRS-CSx). This figure has been plotted using a superset of the results shown in Figure 2 and a subset of results available in Supplementary Data 4-6. Here, each point in each scatter plot corresponds to the average performance measure over 5 replicate datasets per simulation configuration. There are 3 simulation parameters, 6 different values per simulation parameter, 15 different auxiliary-target tuples per simulation configuration, making a total of 270 points per scatter plot.

#### 2.3.3 MultiPopPred outperforms SOTA methods, especially at low target sample sizes

In the second analysis, *h*^2^ was fixed at 70% (mimicking the typically estimated heritability for known complex traits such as height, schizophrenia, etc. [31, 32]) along with 1000 target samples while the number of auxiliary samples per population was varied from 500 to 20,000. As expected, with an increase in the number of auxiliary samples, the corresponding GWAS estimates became more statistically powerful, thereby resulting in an improved transfer learning to the target population (see Supplementary Data 5). It was also observed that in our simulations, around 5000-10,000 auxiliary samples per population should suffice to do an efficient transfer learning operation, following which the performance saturates.

In the third analysis, *h*^2^ was fixed at 70% along with 10,000 auxiliary samples per population while the number of target samples was varied from 100 to 2000 (see Supplementary Data 6). All five versions of MultiPopPred were observed to clearly outperform other methods at low target samples, depicting its robustness (Figure 2a). In the low target sample setting (Figure 2a), MPP-PRS+ demonstrated an 52%, 92%, and 159% average improvement in correlation ratio over SBayesRC-Multi, PROSPER, and PRS-CSx, respectively. For comparison, note that these gains are respectively 28%, 7% and 23% in a high target sample size setting (Figure 2b). While in general, the higher the number of target samples, the better it is for prediction, through this experiment, we observed that MultiPopPred can be used reliably with a number of target samples even 10x to 100x times less than the auxiliary populations (given that the auxiliary populations have sufficiently large sample sizes ≥ 10, 000).

#### 2.3.4 MultiPopPred performs on-par or better than SOTA methods for binary traits

We further investigated the performance of our approach relative to SOTA methods when subjected to simulated binary phenotypes under five different simulation settings, a subset of which is depicted in Figure 4. These five simulated settings were designed to correspond to (i) low target sample size (100), (ii) high heritability (70%), (iii) low heritability (30%), (iv) low auxiliary sample size (500), and (v) low inter-population correlation (0.3), respectively (see Supplementary Data 7). Binary phenotypes were simulated following a liability-thresholding model (see Methods). Consistent with the trends observed in simulated quantitative traits, MPP-PRS+ remained the best-performing version of our approach. MPP-PRS+ reported a 31%, 52%, and 72% gain in performance over SBayesRC-Multi, PROSPER, and PRS-CSx, respectively, in low target sample settings. These performance gains were 33%, 30%, and 57%, respectively, across all simulation settings.

**Fig. 4.**
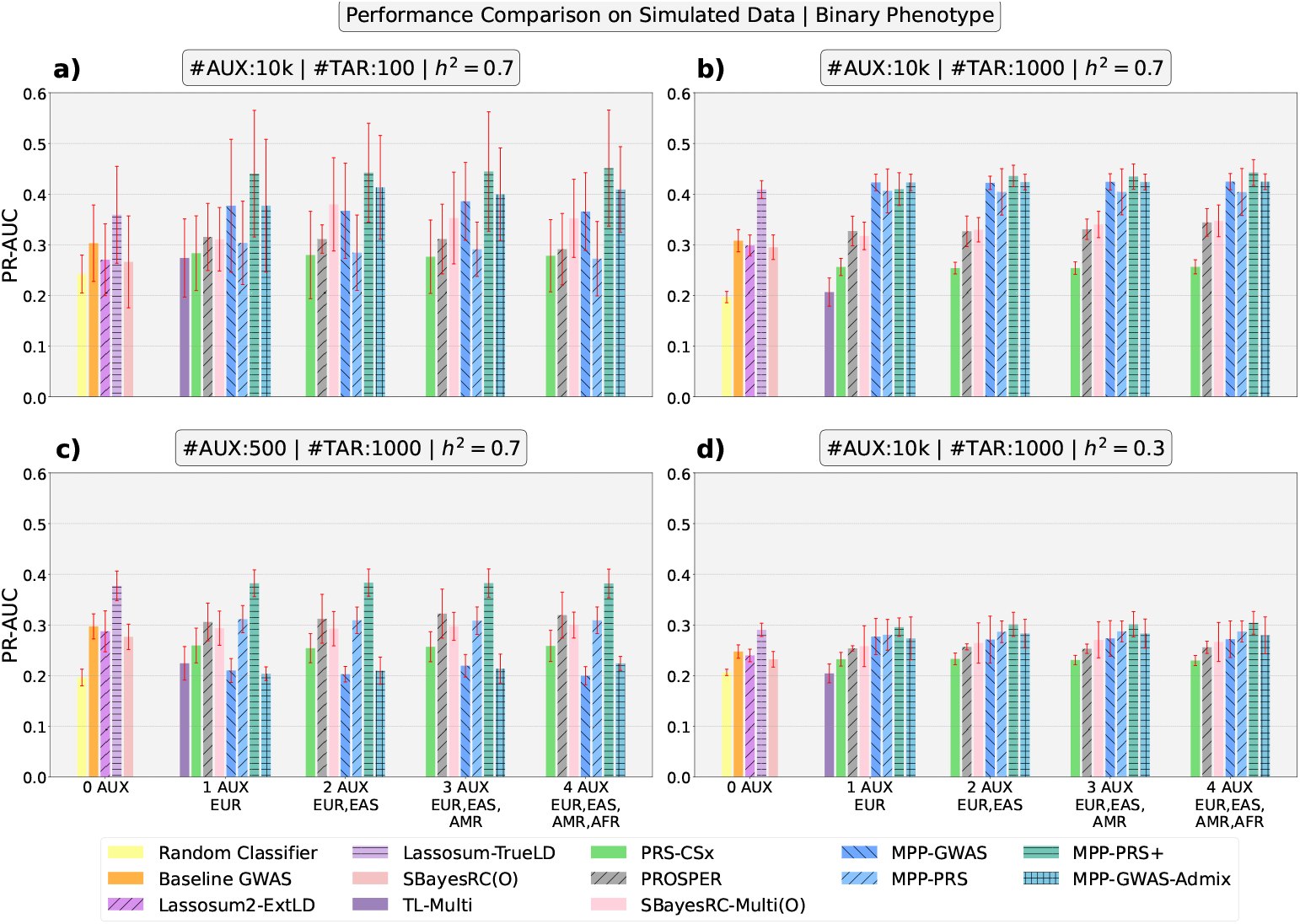
Performance Comparison on Simulated Data (Binary Phenotype Prediction): We depict the comparative performance of MultiPopPred with SOTA methods under four different simulation settings and four auxiliary-target population tuples. For all four simulation settings, inter-population correlation was set to 0.8, and a prevalence of 20% was used for cases. Simulation configurations for panels (A-D), abbreviations used, and non-applicability of certain methods remain the same as in Figure 2. Additionally, MPP-GWAS-TarSS was excluded from these analyses due to the absence of a TarSS framework compatible with the logistic module. Performances were evaluated using Area Under Precision-Recall Curve (PR-AUC).

#### 2.3.5 Evaluating MultiPopPred on PROSPER’s benchmarks

The results presented thus far focused on our simulation strategies for genotypes and phenotypes, where the ground-truth effect sizes of SNPs for each population (target as well as auxiliary) were designed to be spread around 0 as per a normal distribution. This design choice was made to maintain close resemblance to the infinitesimal model or omnigenic model [33–35] of phenotypes such as height, that are known to have a high degree of polygenicity [36]. A different (non-infinitesimal) model of SNP effect size distribution popular in the literature and adopted in PROSPER’s benchmarks is the sparse model, wherein a small number of causal SNPs drive the entirety of the genetic component towards the phenotype, and the other non-causal SNPs have zero effect sizes. This sparse effects model is closer to the oligogenic model of certain real-world traits [37, 38] such as gene expression [39], or lipid-related [40–42] traits, whose genetic architecture lies somewhere between a monogenic and polygenic model. Either models of SNP effect sizes may be pertinent depending on the trait considered. Methods that learn tighter estimates of *β*s (concentrated around 0 with a high peak at the mean) are expected to perform better for traits following the sparse model, whereas those that learn *β* estimates spread around 0 are expected to perform better for omnigenic traits.

To test the applicability of MultiPopPred under a sparse genetic model, it was run on three different simulation benchmarks [43] proposed by PROSPER [21] and its predecessor CT-SLEB [43] (see Supplementary Table 2). SBayesRC-Multi and PROSPER were observed to be the top two performers on all three benchmarks. While MPP-PRS+ was observed to perform worse than PROSPER on all three benchmarks, the gap in MPP and PROSPER’s performance remained relatively smaller for the first benchmark (0.01 fraction of causal SNPs). This gap in performance was seen to increase significantly as the fraction of causal SNPs was reduced progressively to 0.001 and 0.0005 in benchmarks two and three, respectively. This can be attributed to the fact that PROSPER produces *β* estimates more concentrated around 0 than MPP, thereby aligning more with their simulation framework (Supplementary Figure 2). Conversely, the better performance of MPP over PROSPER in our simulations can be attributed to the fact that MPP learns more spread out *β*s, which is in line with our simulation framework (Supplementary Figure 3). The insights from this analysis indicate that, depending on the underlying sparse versus infinitesimal distribution of SNP effect sizes for a given trait, SBayesRC-Multi/PROSPER, or MPP could be used appropriately to compute PRS (see Discussion).

#### 2.3.6 Secondary simulations: Robustness of MultiPopPred under varying input scenarios

We performed a set of secondary simulation analyses to further validate the results and observations mentioned above and to refine our understanding of the working of MultiPopPred under various conditions.

First, we benchmarked the performance of MultiPopPred against HAPNEST [44], a publicly available simulated genotype dataset, in low and high target sample size settings similar to Figure 2a and 2b. This simulation was performed with two different sets of SNPs, with and without LD structures. In accordance with previously observed results, MultiPopPred was seen to clearly outperform SOTA methods (Supplementary Figure 4), with PROSPER being the second-best performer.

Next, we evaluated MultiPopPred under six simulation settings, comprising all combinations of three inter-population correlation levels (low: 0.3, moderate: 0.6, and high: 0.8) and two target sample size levels (low: 100 and high: 1,000). Supplementary Figure 5 depicts that MultiPopPred maintains its performance trends over SOTA in the case of low and moderate inter-population correlations (see Supplementary Results 1.2.1).

Finally, we performed a series of ablation studies and sensitivity analyses to quantify how individual components of MultiPopPred drive performance. Our results demonstrate that the combination of the L-BFGS optimizer, Nesterov smoothing, and *L*_1_ penalization works in coherence to achieve the observed gains (detailed in Methods, Supplementary Results 1.1, and Supplementary Figures 6–11). Furthermore, the consistently better performance of MPP-PRS+ over other MPP versions and SOTA methods (all of which rely on summary-level data of auxiliary populations) advocates the use of individual-level data and associated true LD of the target and auxiliary populations.

To summarize, MPP-PRS+ was observed to report an overall 38% improvement in PRS prediction on average across all simulation settings and 91% across simulation settings with low target sample sizes when compared with SOTA methods. Individually, each MultiPopPred version reported an overall improvement of 38% (MPP-PRS+), 11% (MPP-PRS), 43% (MPP-GWAS), 73% (MPP-GWAS-TarSS), and 44% (MPP-GWAS-Admix), over all SOTA methods and all simulation settings (both quantitative and binary phenotypes, wherever applicable). Supplementary Data 8 presents a detailed account of percentage improvements obtained by the different MultiPopPred versions against all SOTA methods under various simulation settings.

### 2.4 Benchmarking using semi-simulated data

Semi-simulated data analysis was performed using real-world genotype data on EUR, EAS, AFR, and SAS populations from UK Biobank [45] while phenotypes were simulated as per the procedure described in Section 2.3 (also, see Methods). We performed a comparative analysis of MultiPopPred and other SOTA methods with respect to phenotype predictions under two different semi-simulated settings corresponding to low and high target sample sizes. Similar to the fully simulated analysis, each semi-simulation was repeated 5 times, and for each setting, MultiPopPred and other methods were run under 3 different configurations of auxiliary-target population tuples (see Figure 5, Supplementary Data 9). On average, MPP-PRS+ was observed to outperform SBayesRC-Multi, PROSPER, and PRS-CSx by a significant margin of 382%, 84%, and 123%, respectively and by 148% overall (see Supplementary Data 8). It must still be maintained that the drop in PROSPER’s performance, especially in setting with low target sample sizes, can be attributed to the non-suitability of the extreme setting for the method.

**Fig. 5.**
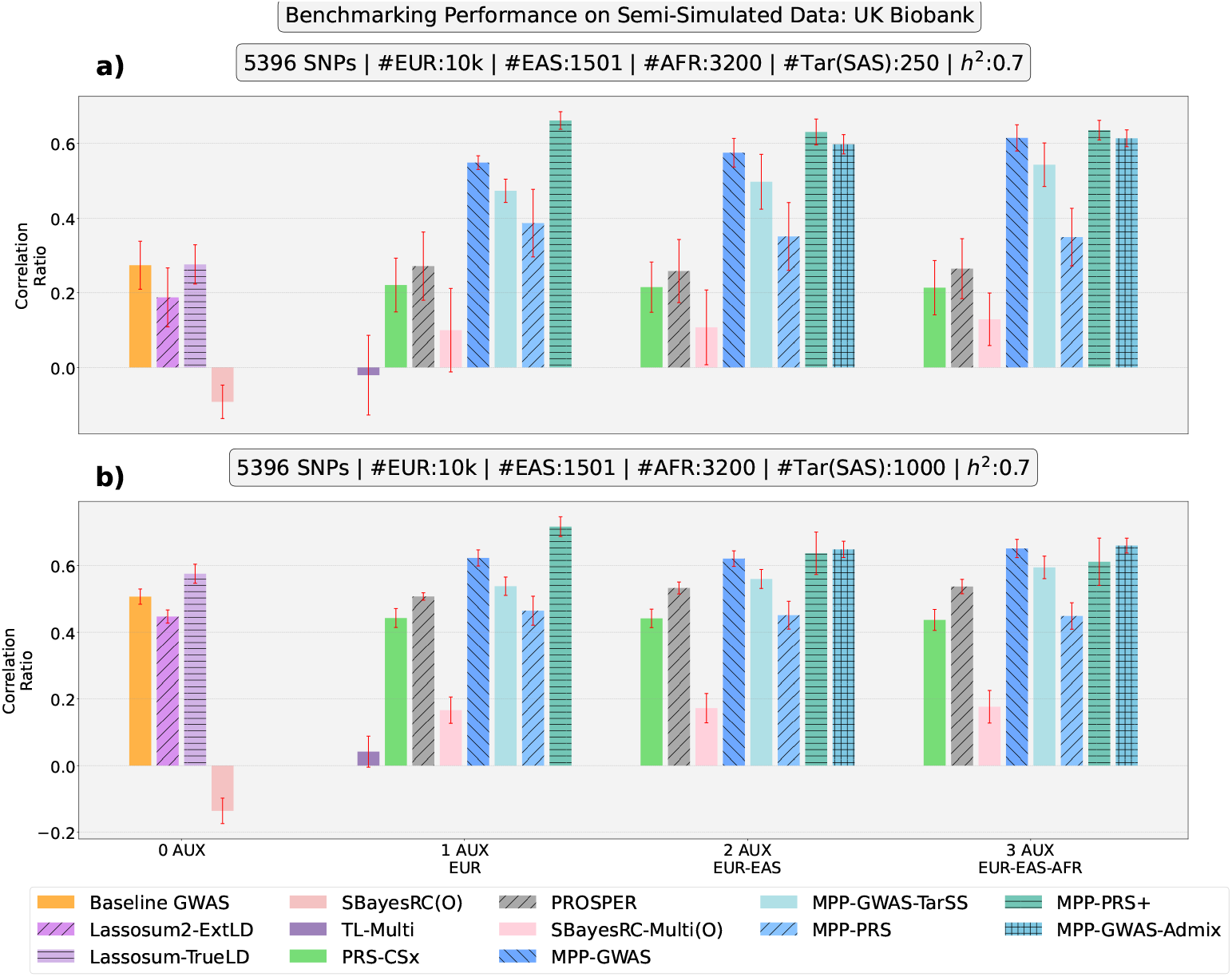
Performance Comparison on Semi-Simulated Data (Phenotype Prediction): We depict the comparative performance of MultiPopPred with SOTA methods under two different semi-simulation settings and three auxiliary-target population tuples. First, the number of samples per auxiliary population was set to 10,000 for EUR, 1501 for EAS, and 3200 for AFR, and the number of target population samples was set to 250, with *h*^2^ being 70%. Next, the number of auxiliary samples and *h*^2^ remained the same as before, but the number of target samples was increased to 1000. The analysis was restricted to a set of 5396 SNPs (from chromosome 22) that were common across UK Biobank and 1000 Genomes. SBayesRC(O) and SBayesRC-Multi(O) refer to the “oracle” version of these two methods. Abbreviations: Aux: Auxiliary Population, Tar: Target Population, *h*^2^: Heritability.

### 2.5 Benchmarking using real-world data

Real-world analysis was performed on 8 quantitative traits (namely Height, Body Mass Index - BMI, Systolic Blood Pressure - SBP, Diastolic Blood Pressure - DBP, High-Density Lipoprotein - HDL, Low-Density Lipoprotein - LDL, Total Cholesterol - TC, and Triglycerides - TG), and 8 binary traits from UK Biobank (namely Risk Taking - RT, Educational Qualification - EDQ, Allergy - ALG, Any Cardiovascular Disorder - Any CVD, Asthma - AST, Dyslipidemia - DLP, Morning Person - MP, and Type 2 Diabetes - T2D) (see Methods; and Supplementary Data 10 and 11). These traits were chosen to cover a range of heritability (*h*^2^) values among the quantitative traits (four having relatively higher heritability values and four having relatively lower heritability values) and a range of prevalences among the binary traits. The *h*^2^ values for these traits reported in earlier studies [20, 46, 47] are consolidated in Supplementary Table 3. SAS was set as the target population while EUR, EAS, and AFR were used as auxiliary populations wherever applicable.

For each trait, MultiPopPred and SOTA methods were evaluated across 3 random training-validation-testing splits of the dataset, and the average performance was reported, along with an error bar indicating twice the standard deviation. Performance was measured using *R*^2^ for quantitative traits (instead of correlation ratio due to the absence of ground-truth effect sizes of SNPs) and PR-AUC for binary traits. Please refer Supplementary Methods 2.1 and 2.2 for our detailed prediction and evaluation strategy respectively. For each trait, Supplementary Data 12-14 contain a detailed account of the training, validation, and testing performances of all methods; and Supplementary Data 15 and 16 report the percentage differences as well as delta differences (Δ*R*^2^ for quantitative traits and ΔPR-AUC for binary traits) between the different methods.

#### 2.5.1 Heuristic PRS versions show promise of GWAS as a new baseline

Traditionally, whenever a PRS estimate is talked about, one refers to a joint all-SNP model for training and subsequent predictions, as opposed to a per-SNP model. We observe in our real-world analysis that the Baseline GWAS approach, despite being a per-SNP model, performs decently enough across all traits (see Figures 6, 7, and Supplementary Data 15, 16). This observation warrants further exploration of heuristic-based PRS approaches, which rely on auxiliary *β*s estimated from (per-SNP) GWAS. To further investigate this trend observed in heuristic PRS versions, we performed a comparative analysis of Baseline GWAS alone with Clumping and Thresholding (C&T) [43] — a traditional and very simplistic PRS estimation method that has served as the baseline for most previous SOTA methods existing in the literature. Supplementary Figure 12 shows that at stringent p-value cutoffs (5 *×* 10^−8^, 5 *×* 10^−7^, 5 *×* 10^−6^), C&T performs poorly compared to Baseline GWAS (see Supplementary Data 17). As the p-value threshold increases, the performance of C&T improves, reaching an on-par or in some cases a better level than Baseline GWAS. An exception is observed in case of HDL wherein C&T’s performance peaks at a 5 *×* 10^−8^ p-value cutoff and worsens progressively, indicating that the trait is likely driven by a few very powerful SNP effects rather than thousands of tiny ones. Similarly for TG, C&T deviates showing a U-shaped performance (close to Baseline GWAS around 5 *×* 10^−8^, dipping in the middle around 5 *×* 10^−5^ and rising up again close to 1). Overall, these performance trends indicate that while filtering the set of available SNPs prior to PRS computation is essential, it must not come at the cost of truly causal or truly informative SNPs. Retaining additional SNPs during PRS computation generally does not seem to harm the predictive performance by a huge margin. A logical solution to this situation could be to use the complete available set of SNPs for PRS estimation while employing sufficiently regularized models (like MultiPopPred or PROSPER) to ensure that the contribution of non-informative SNPs towards prediction is controlled. Furthermore, the use of simple GWAS as the baseline should be promoted, as opposed to C&T, while proposing novel PRS estimation methods.

**Fig. 6.**
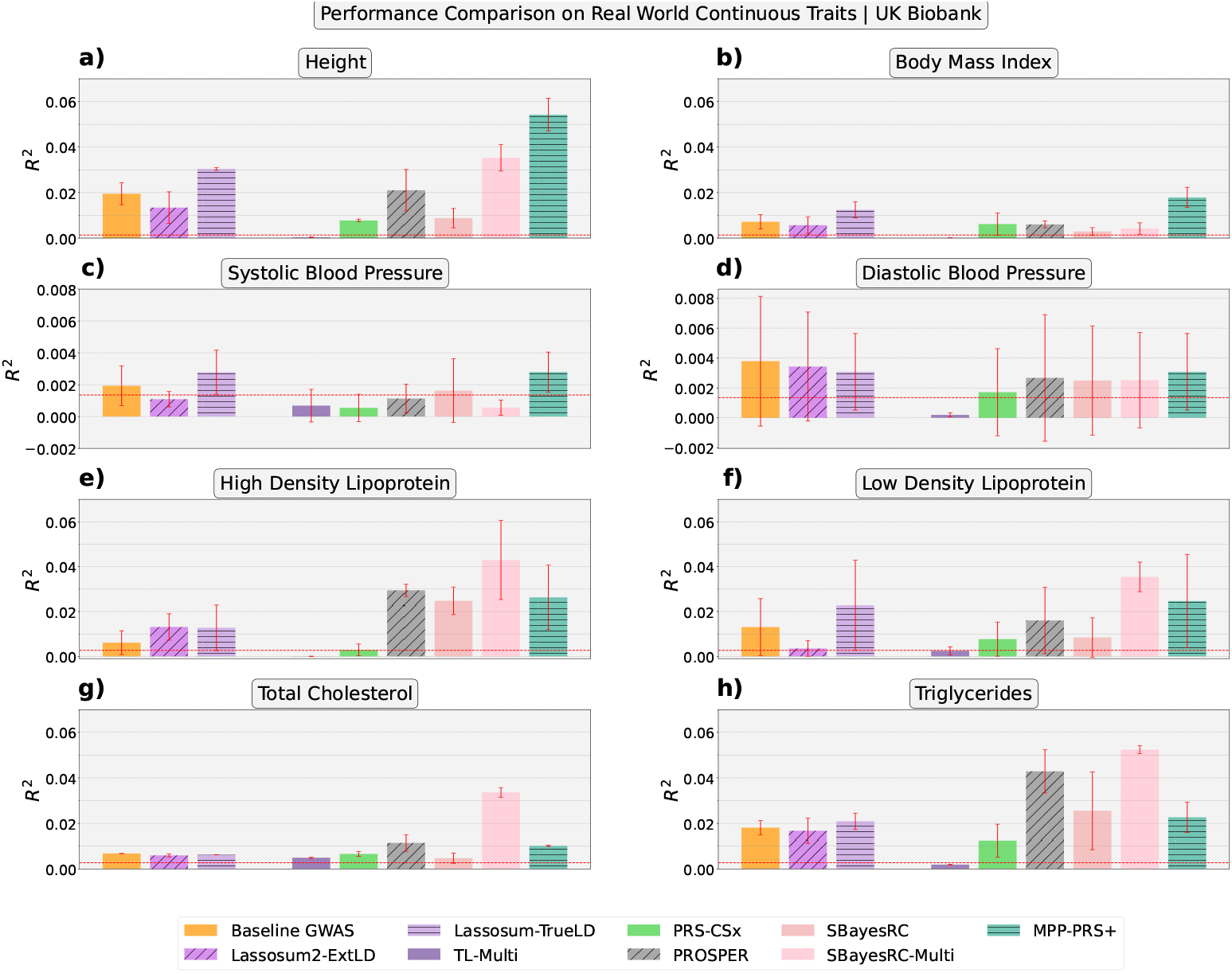
Performance Comparison on Real-World Data (Phenotype Prediction): We depict the comparative performance of MultiPopPred with SOTA methods when applied to 8 different quantitative traits from UK Biobank.The horizontal red line in each panel indicates the critical *R*^2^ required to achieve a statistically significant (Pearson’s) correlation at *α*-level 0.05 between the PRS and ground-truth phenotype. Supplementary Figure 13 depicts a zoomed-in version of each individual panel in this figure for better clarity of relative trends between different methods, including other versions of MPP.

**Fig. 7.**
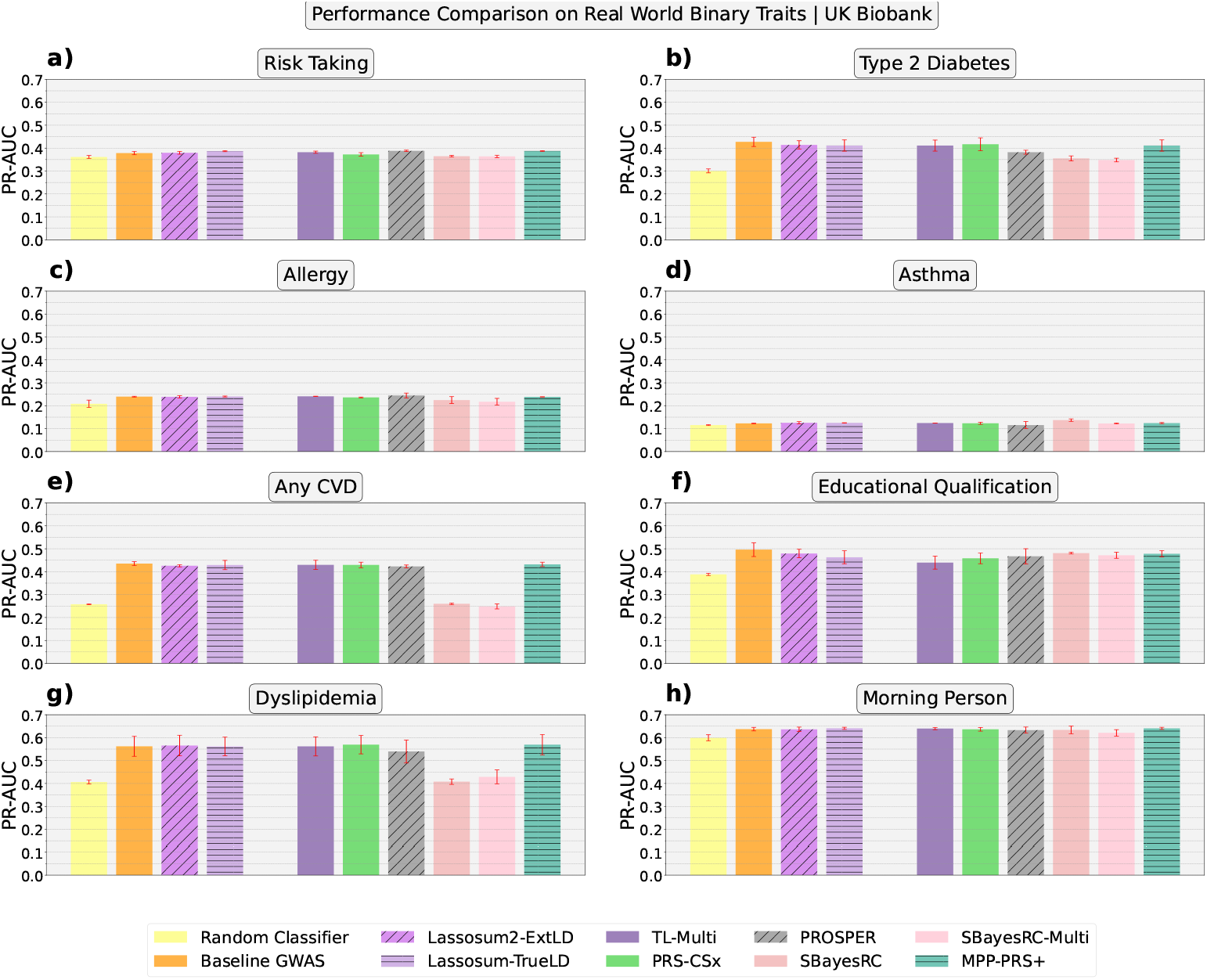
Performance Comparison on Real-World Data (Phenotype Prediction): We depict the comparative performance of MultiPopPred with SOTA methods when applied to 8 different binary traits from UK Biobank. Supplementary Figure 17 depicts a zoomed-in version of each individual panel in this figure for better clarity of relative trends between different methods, including other versions of MPP.

#### 2.5.2 MultiPopPred-PRS+ demonstrates superior performance across real-world omnigenic traits

MPP-PRS+ was observed to be the best performer among the five MultiPopPred versions, which is consistent with the trends observed in our simulation and semi-simulation analyses. This improved performance is also in line with the full information, i.e., individual-level target and auxiliary information available to MPP-PRS+. PROSPER and SBayesRC-Multi use individual-level target information only during hyperparameter tuning in the validation dataset, while PRS-CSx uses only summary-level information from the target and auxiliary populations (see Methods). Figure 6 shows that MPP-PRS+ outperforms all SOTA methods for Height and BMI on average. MPP-PRS+ further demonstrates a slight edge over other methods for SBP and DBP albeit with overlapping error bars for several SOTA methods, indicating that these two traits are inherently difficult to predict irrespective of the method. We categorize these four traits as omnigenic and the remaining four lipid traits (HDL, LDL, TC, and TG) as oligogenic, in line with existing studies [35, 36, 40–42]. For the lipid-related traits, MPP-PRS+ was seen to be lagging in performance compared to leading SOTA like SBayesRC-Multi and PROSPER, in line with similar trends observed for simulated sparse effect traits in Section 2.3.5. Overall, these results suggest that for low-resource populations, individual-level information should be used for training as much as possible, so that the LD structure used by the method matches the true LD structure of the population being studied and thereby maximizes predictive power.

To verify if the gains obtained by MPP-PRS+ with true LD were not putting other SOTA methods (which do not use true LD information) at an unfair disadvantage, we provided each method with the exact same set of inputs pertaining to all the quantitative UK Biobank traits. As detailed in Supplementary Results 1.2.2, for each quantitative trait, MPP-PRS+, PROSPER, and SBayesRC-Multi (and their respective base layers) were each run under two different settings — full access to the target as well as auxiliary populations’ (i) True LD, and separately (ii) External LD reference panels. The performance trends were observed to remain largely the same under this fair comparison, as shown in Figure 6 for all three methods, even with true LD. This implies a better effectiveness of MPP-PRS+ in capturing the information available through true LD, specifically for the non-lipid traits. MPP-PRS+ was also observed to exhibit a higher sensitivity to LD mismatch than PROSPER and SBayesRC-Multi, manifesting as a noticeable drop in performance when supplied external LD. We further investigated these performance gains by decomposing them into gains from each method’s base layer (only auxiliary population-based predictions) versus the meta layer (post transfer learning predictions). We observed that the performance improvement of MPP-PRS+ over PROSPER (under true LD) and SBayesRC-Multi (under true LD) came from gains in both base and meta layers over the two methods (see Supplementary Results 1.2.2 for details). These analyses are further elaborated in Supplementary Results 1.2.2, Supplementary Data 18-21, and Supplementary Figures 14-15.

Supplementary Figure 16 depicts the distribution of percentage improvement reported by MPP-PRS+ over two leading SOTA methods, SBayesRC-Multi and PROSPER, with respect to all 8 quantitative and all 8 binary real-world traits (thereby expanding on the results in Figure 6 and 7 respectively). MPP-PRS+ was observed to outperform SBayesRC-Multi by average margins of 55% (Δ*R*^2^ 0.019), 469% (Δ*R*^2^ 0.0137), 1498% (Δ*R*^2^ 0.0022), and 101% (Δ*R*^2^ 0.0006) for Height, BMI, SBP, and DBP, respectively, while lagging behind by 24% (Δ*R*^2^ −0.0167), 36% (Δ*R*^2^ −0.0108), 69% (Δ*R*^2^ −0.0235), and 54% (Δ*R*^2^ −0.0287) for HDL, LDL, TC, and TG, respectively (see Supplementary Data 15). Similarly, against PROSPER, MPP-PRS+ reported average performance gains of 178% (Δ*R*^2^ 0.0332), 219% (Δ*R*^2^ 0.0119), 414% (Δ*R*^2^ 0.0017), 2195% (Δ*R*^2^ 0.0004), and 170% (Δ*R*^2^ 0.0087) for Height, BMI, SBP, DBP, and LDL respectively, and average performance drops of 8% (Δ*R*^2^ −0.0031), 3% (Δ*R*^2^ −0.0013), and 44% (Δ*R*^2^ −0.0192) for HDL, TC, and TG, respectively. It must be noted, however, that for SBP and DBP, all methods report an *R*^2^ marginally above the critical *R*^2^, indicating that these traits are inherently more difficult to predict and/or suffer from a higher noise-to-signal ratio.

In case of binary traits, Any CVD, DLP, T2D, ALG, and RT, MPP-PRS+ was observed to outperform SBayesRC-Multi by margins of 74% (ΔPR-AUC 0.1834) (Figure 7e), 32% (ΔPR-AUC 0.1405) (Figure 7g), 18% (ΔPR-AUC 0.0632) (Figure 7b), 10% (ΔPR-AUC 0.0213) (Figure 7c), and 6% (ΔPR-AUC 0.0235) (Figure 7a) on average respectively. For all other binary traits, MPP-PRS+ reported comparable performance (within 5% of SBayesRC-Multi’s performance), indicating no unanimous winner (see Supplementary Data 16). Similarly, with respect to PROSPER, MPP-PRS+ reported superior performance on binary traits AST (9%, ΔPR-AUC 0.0088), T2D (8%, ΔPR-AUC 0.0291), and DLP (6%, ΔPR-AUC 0.0297) while remaining comparable (within 5% of PROSPER’s performance) for the remaining 5 binary traits.

Figure 8 summarizes the improvement attained by MPP-PRS+ over three leading SOTA methods SBayesRC-Multi, PROSPER, and PRS-CSx. As a final robustness check, we carried out decile calibration analysis [48–50] based on MPP-PRS+ predictions for all 8 quantitative (Supplementary Figure 20) and 8 binary (Supplementary Figure 21) real-world traits to assess the practical impact of the predictions. We observed that our model was well calibrated for 4 out of 8 quantitative traits (Height, BMI, HDL, and TG) and 4 out of 8 binary traits as well (Any CVD, DLP, T2D, and EDQ). For the other traits, while our model appears slightly miscalibrated, it does report an acceptable Bias (≈ − 0.55) for quantitative traits and E/O ratio (≈ 1.006) for binary traits (see Supplementary Figure 20 and 21 captions for formal description of these measures). This indicates that for these specific traits, the model correctly predicts the mean deciles (i.e., achieves global calibration) but suffers from certain local miscalibrations.

**Fig. 8.**
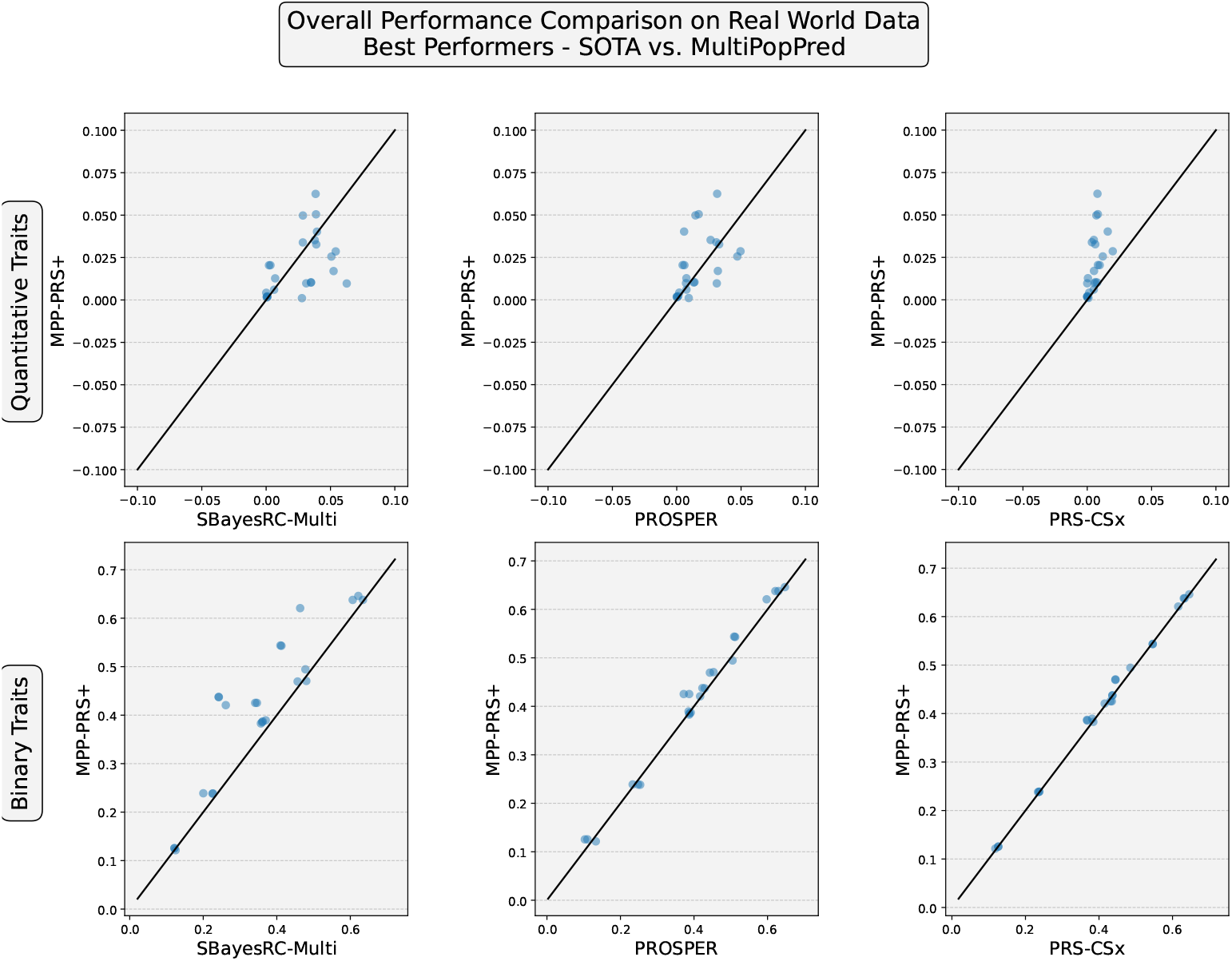
Overall Performance Comparison on all real-world traits from UK Biobank (with respect to Phenotype Prediction using *R*^2^ for quantitative traits and PR-AUC for binary traits): We depict the comparative performance of MultiPopPred-PRS+ against the three best performing and leading SOTA methods (SBayesRC-Multi, PROSPER and PRS-CSx). This figure has been plotted using a subset of the data shown in Figures 6 and 7. Here, each point in each scatter plot corresponds to the performance measure for an individual replicate dataset for a trait. There are 8 traits and 3 replicate datasets per trait, making a total of 24 points per scatter plot. Supplementary Figures 18 and 19 depict similar scatter plots comparing the overall performance of all MultiPopPred versions with all SOTA methods.

## 3 Discussion

In this work, we have proposed MultiPopPred, a novel trans-ethnic PRS estimation method that aims to harness the transferability of learned information from multiple auxiliary populations to a target population. Through extensive simulation and semi-simulation analyses, assuming an infinitesimal model and covering a range of trait heritabilities, auxiliary and target population sample sizes, we have observed that MultiPopPred robustly outperforms other methods in extreme conditions where there is a dearth of target samples. MPP-PRS+ was seen to emerge as the default and the best performing version of MultiPopPred, owing to its use of full information (true LD) from target and auxiliary individual-level data. We believe this is an important conclusion from our comprehensive benchmarking and emphasizes the use of true LD information for reliable PRS prediction, especially in the context of low-resource populations. Subsequent evaluations on application to quantitative and binary real-world traits from UK Biobank report superior performance of MultiPopPred on a majority of traits (which are omnigenic in nature), with the exception of 4 lipid traits (HDL, LDL, TC, and TG). A key highlight is the superior performance of MultiPopPred on the binary traits Any CVD, Type 2 Diabetes, and Dyslipidemia, all of which are reported in the literature and/or UK Biobank to have an increased prevalence in the South Asian population (see Introduction).

MultiPopPred’s effective performance in a majority of simulated and real-world scenarios could potentially be attributed to the use of (i) true LD from the target population and (ii) Nesterov-smoothing along with the L-BFGS optimizer which seems to be more efficient at locating the optima than coordinate descent which was employed by most of the other methods. Both these potential reasons were verified via ablation studies and sensitivity analyses. As in any transfer learning approach, MultiPopPred is expected to perform better when the auxiliary populations, from which transfer of knowledge occurs, are sufficiently closely related to the target population in terms of LD structure and hence the underlying distribution of SNP effect sizes. In the absence of such genetically similar auxiliary populations, MultiPopPred’s predictive performance is expected to be guided largely by the number of target and auxiliary samples available. A surprising takeaway from our simulation, semi-simulation, and real-world analyses was the decent performance of the heuristic PRS versions of MultiPopPred that use auxiliary *β*s estimated from (per-SNP) GWAS. This observation demonstrates the potential and scope for further exploration of per-SNP GWAS as the standard baseline to benchmark methods against as opposed to the traditionally used C&T.

We now arrive at our second important conclusion — MultiPopPred is expected to report larger improvements in performance for a given trait when the underlying distribution of SNP effect sizes follows a non-sparse, i.e., infinitesimal/omnigenic model. When evaluated on PROSPER’s simulated benchmarks, which include traits with a sparse underlying distribution of SNP effect sizes, MultiPopPred reported marginally worse to comparable performance on one benchmark (1% causal SNPs) while PROSPER was observed to perform better on the other two benchmarks (0.1% and 0.05% causal SNPs). SBayesRC-Multi was observed to have an edge over both PROSPER and MultiPopPred on all three benchmarks. MultiPopPred was also observed to report a less competitive performance on 4 real-world lipid traits from UK Biobank. We hypothesize that these trends can be explained through the underlying genetic architecture of the specific traits. The 4 lipid traits (HDL, LDL, TC, and TG) exhibit characteristics of an oligogenic architecture (a small number of SNPs contributing towards a major chunk of the explainable variation in the phenotype as opposed to a large number of SNPs contributing with small to moderate effect sizes towards the phenotypic variance). This is evidenced by highly significant peaks ( − log_10_ *p* ∈ [140, 250]) observed on chromosomes 16 (HDL), 19 (LDL, TC), and 11 (TG) in our European GWAS Manhattan plots (Supplementary Data Plots 25.5-25.8). These observations also find support in existing GWAS conducted on lipid traits across large and small consortia [40–42]. MultiPopPred’s performance likely degrades in such highly sparse settings due to excessive shrinkage, where large-effect SNPs may be overly penalized or treated as noise. In contrast, traits like Height or BMI, where MultiPopPred performs well, exhibit a more polygenic signal [35, 36] distributed across the chromosomes.

Taking into account these observed performance trends on simulated infinitesimal traits as well as real-world lipid and non-lipid traits, we put forward a triage recommending the use of MultiPopPred and competing SOTA methods (SBayesRC-Multi and PROSPER) depending on the genetic architecture (infinitesimal versus sparse) and nature (quantitative versus binary) of the trait in consideration. We posit that in the case of quantitative traits exhibiting characteristics of an infinitesimal/omnigenic model, MultiPopPred (specifically MPP-PRS+) would be the best choice. Conversely, in the case of quantitative traits exhibiting characteristics of a sparse/non-infinitesimal/oligogenic model, either of SBayesRC-Multi or PROSPER would be a reasonable choice. In the case of binary traits, we recommend the use of any one of MPP-PRS+, SBayesRC-Multi, or PROSPER, irrespective of the underlying genetic architecture, albeit with a relatively higher inclination towards MPP-PRS+ given its superior performance on prevalent disease traits in the South Asian population.

We also note that there is divergence in MultiPopPred’s performance between quantitative lipid traits (where MPP underperforms) and binary DLP (where MPP outperforms). This divergence is likely driven by the way the phenotypes are measured or defined. As per UK Biobank documentation, the quantitative lipid traits were based on NMR metabolomics profiling of the study participants. On the other hand, the binary case/control assignments in DLP were clinically determined on the basis of standard ICD-10 (E78) codes, rather than being derived directly from the quantitative lipid traits (Supplementary Data 10). Further, we observed the Point Biserial correlation (*R*_*pb*_) of DLP with each quantitative lipid trait to be low in magnitude albeit showing statistical significance (*R*_*pb*_ = −0.13, −0.08, −0.12 and 0.14 for DLP and HDL, LDL, TC, and TG respectively across 272, 595 individuals with both measurements). Together, these observations indicate that DLP is not strongly correlated with the quantitative lipid traits, consistent with the diverging performance.

A major bottleneck observed while estimating PRS in low-resource populations, such as the South Asian population, was the limited availability of samples for training, validation, and testing. While methods like MultiPopPred provide a reasonable alternative in such extreme situations, we must set realistic expectations on the maximum achievable performance in such cases. To set a benchmark on the variation of predictive performance as the sample size increases, we conducted a simple analysis on the EUR samples from UK Biobank for the trait Height. While all our previous analyses have treated SAS as the target population, this time, EUR was chosen as the target owing to the flexibility offered by EUR in switching from very low sample sizes to very high sample sizes through simple subsetting of samples at random. As the training sample size increased, we noted the performance of Baseline GWAS and Lassosum2, which denote prediction approaches that apply the base layer of MultiPopPred heuristic and PRS versions (GWAS and Lassosum2, respectively) on the target population. Supplementary Data 22 shows that the best performance (test *R*^2^ = 0.047 for Baseline GWAS and 0.081 for Lassosum2) was observed with training samples as high as 432,526. As the sample size was decreased, the performance was also observed to dip proportionately (test *R*^2^ = 0.019 for Baseline GWAS and 0.006 for Lassosum2 with 10, 000 training samples). This shows that although at the outset, the observed absolute *R*^2^ values in our real-world analyses might seem pretty low from a standard machine learning point of view (and may even indicate overfitting when training and test performances are compared), on a relative scale to the best achievable performance, they demonstrate good predictive power.

While the observed results are promising, MultiPopPred has certain limitations, such as its less competitive performance on non-infinitesimal/oligogenic traits as already mentioned above. In addition, a limitation of MPP-GWAS-Admix is it’s exclusive reliance on genotypic information to estimate admixture proportions and thereby the weights of populations, rather than tailoring it to the specific phenotype being considered. Such phenotypic dependencies in admixture proportions are something that needs further investigation in the future. Some more challenging aspects in the field of trans-ethnic PRS estimation that could benefit from future investigation are the inclusion of rare or population-specific SNPs, the inclusion of non-linear effects (epistasis) of SNPs towards the phenotype, and the inclusion of non-genetic, environmental (often unobserved confounders) while determining a target individual’s disease risk. Careful consideration and redressal of such challenges in future work can open up the advantage of transfer learning to more (non-linear or epistatic) scenarios. Nevertheless, through comprehensive simulation and real-world benchmarking, we have demonstrated that the existing version of MultiPopPred still performs comparably or better than existing SOTA methods when individual-level data is available as per the triage recommendations above, and can pave the way for its application to understand the genetic architecture of multiple phenotypes in understudied populations.

## 4 Methods

### 4.1 MultiPopPred

#### 4.1.1 Background and preliminary notations

Traditional GWAS rely solely on a simple (per-SNP) univariate linear regression approach to arrive at per-SNP effect sizes or *βs*. Traditional PRS employs a joint (all-SNP) multivariate linear regression model (as depicted in Equations 1a and 1b below).

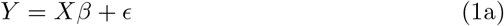

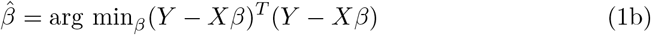

Here, *X* represents the *n × m* genotype matrix (where *n* is the number of individuals in the study and *m* is the number of SNPs), *Y* is the *n ×* 1 phenotype (or disease trait) vector, *β* is a *m ×* 1 vector containing per SNP effect sizes, and *ϵ* is the *n ×* 1 noise vector. Traditionally, the genotype matrix *X* is normalized so that each column of the matrix has a zero mean and unit variance. Similarly, the phenotype vector *Y* is also normalized so that it has a zero mean and unit variance. In addition to the standard assumptions associated with a simple linear regression model [51], this approach assumes that (i) the SNPs have a linear and additive relationship with the phenotype, (ii) there are no interactions among the SNPs themselves, (iii) the noise term includes contributions from other non-additive and environmental factors. Traditional GWAS and/or PRS approaches work well when there are large enough samples, but fail to produce reliable estimates of *β* in case of smaller sample sizes.

#### 4.1.2 MultiPopPred objective function

Let us define **auxiliary populations** as those that are high-resource, or in other words, have a sufficiently large sample size (*n*) to estimate statistically reliable SNP effect sizes (*β*s). Let us define a **target population** as one that contrastingly has considerably fewer number of samples available (small *n*, usually in the range of a few hundreds to a couple of thousands). Let us say, we have *K* auxiliary populations with *β* estimates 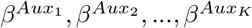, with each 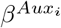 having dimension (*m ×* 1). Additionally, we have the target population’s genotype 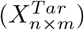 and phenotype 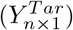 information. We propose to solve Equation 1a above through an *L*_1_ penalized regression approach (Equation 2) that strives to make the target *β* estimates as close as possible to the auxiliary population *β*s, while minimizing the standard least-squares based loss. Here, ℱ denotes the objective function to be optimized, 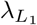 denotes the *L*_1_ shrinkage parameter, and *β*^*Aux*^ refers to an aggregate of 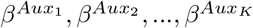. The process of aggregation of these *β*s has been elaborated in Section 4.1.4, where we discuss the different versions of MultiPopPred.

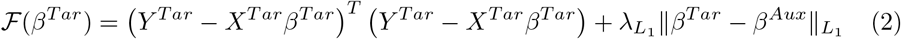

#### 4.1.3 MultiPopPred optimization

Going forward we will drop the superscript *Tar* from *X*^*Tar*^, *Y* ^*Tar*^ and *β*^*Tar*^ for the sake of brevity. For optimization of our objective function (Equation 2), we employ the Limited Memory Broyden-Fletcher-Goldfarb-Shanno (L-BFGS) method [52]. Although L-BFGS is suited for optimizing smooth, differentiable functions, studies have highlighted its applicability and efficiency even in the case of some non-smooth functions and some smooth approximations of non-smooth functions [53]. In our case, we employ Nesterov smoothing (smoothing parameter *µ* fixed at 0.1) [54] with an entropy prox function *f* ^*µ*^(*x*) (Equation 3) [55] to produce a smoothed approximation of the objective function as depicted in Equation 4 below. We supply the first order derivative of Equation 4 to the L-BFGS method for faster convergence, whenever smoothing is used. A detailed step-by-step derivation has been presented in Supplementary Methods 2.3. Furthermore, we conducted several ablation studies and sensitivity analyses to justify the choice of the different components that make up MultiPopPred, such as the optimizer, penalization strategy, smoothing function, and the weighting scheme for auxiliary populations (see Supplementary Results 1.1 and Supplementary Figures 6-11). A consolidated pseudocode detailing the input-output requirements, data pre-processing, and optimization modules of MultiPopPred has also been presented in Supplementary Methods 2.4. A running time and memory usage analysis outlining the computational requirements of MultiPopPred have been made available through Supplementary Results 1.3 and Supplementary Data 23.

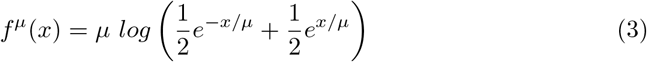

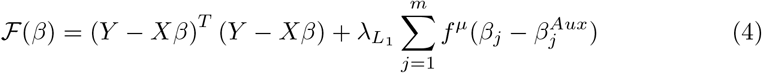

#### 4.1.4 MultiPopPred versions

MultiPopPred-PRS+ (described below) serves as the default version of our approach. In addition to that, we have proposed four other methodological variants of MultiPopPred-PRS+, allowing flexibility in terms of the availability of the quantity and type of auxiliary and target data (i.e., individual-level versus summary-level data, true LD versus external LD) and the manner of quantification of similarity between a given pair of populations. First, we propose two formal PRS versions that perform a joint model PRS to PRS transfer learning. These include (i) MultiPopPred-PRS+, and (ii) MultiPopPred-PRS. Next, we propose three heuristic PRS versions that aim to perform transfer learning directly from per-SNP model GWAS summary statistics to a joint PRS model. These include (i) MultiPopPred-GWAS, (ii) MultiPopPred-GWAS-TarSS, and (iii) MultiPopPred-GWAS-Admixture. Each of these versions is described in detail in the following subsections. It must be noted that we did not develop admixture and summary-level-data-only based versions of MPP-PRS because the advantages offered by MPP-GWAS-TarSS and MPP-GWAS-Admix over MPP-GWAS were observed to be quite subtle in our simulation studies.

##### MultiPopPred — Default version (Full-Data)

###### MultiPopPred-PRS+

This version of MultiPopPred (MPP-PRS+) assumes the availability of complete information on target as well as auxiliary populations, for the purpose of training. In other words, MPP-PRS+ requires the target as well as the auxiliary populations’ genotype (*X*) and phenotype (*Y*), along with their respective GWAS summary statistics as inputs. Auxiliary populations’ single-ancestry PRS is computed using a modified version of Lassosum (Lassosum-TrueLD) that uses the true LD for each auxiliary population computed from its respective genotype. In this version of MultiPopPred, we give equal weights to each auxiliary population. In other words, if there are *K* auxiliary populations each having their own set of *β* estimates (each of size *m ×* 1) from *K* independent joint models, as described in Sections 4.1.1 to 4.1.2, then the aggregated *β*^*Aux*^ is defined as per Equation 5 below.

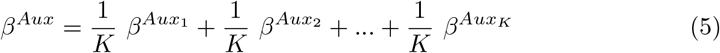

If Nesterov smoothing is used along with this MPP version, the gradient vector (again of size *m ×* 1) supplied to the L-BFGS optimizer can be derived to get Equation 6 below (see Supplementary Methods 2.3.1 for a detailed gradient derivation).

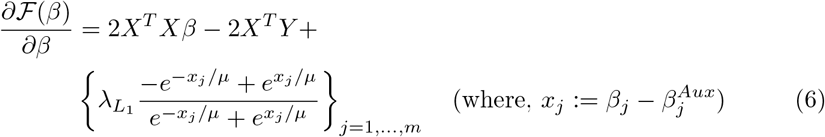

##### MultiPopPred — Other formal PRS versions (Reduced-Data)

###### MultiPopPred-PRS

This version of MultiPopPred (MPP-PRS) relaxes one of the constraints on the input requirements imposed by MPP-PRS+. MPP-PRS assumes that we have the target population’s individual-level genotype and phenotype, along with its GWAS summary statistics, as well as the auxiliary populations’ GWAS summary statistics available to us. However, the auxiliary populations’ genotype and phenotype are not available. For each auxiliary population, MPP-PRS uses its GWAS summary statistics and its external LD reference panel as inputs to Lassosum2 (Lassosum-ExtLD) to compute its single-ancestry PRS. Hereon, MPP-PRS resembles MPP-PRS+ in terms of the weights given to each auxiliary population and the objective function to be optimized.

##### MultiPopPred — Other heuristic PRS versions (Reduced-Data)

The heuristic PRS versions of MPP derive their auxiliary *β* estimates from *K* independent per-SNP models (GWAS) corresponding to the *K* auxiliary populations as opposed to the *K* joint-SNP models employed in the formal PRS counterparts.

###### MultiPopPred-GWAS

This version of MultiPopPred (MPP-GWAS) requires the target population’s genotype (*X*) and phenotype (*Y*), along with its GWAS summary statistics, as well as each auxiliary population’s GWAS summary statistics as inputs. Like the preceding MPP versions, MPP-GWAS also assigns equal weights to each auxiliary population (Equation 5) and uses Equation 6 for optimization. The difference between MPP-GWAS and the preceding two versions, as the name suggests, lies in the basis for transfer learning. In case of MPP-GWAS, 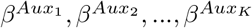 (each of size *m ×* 1) correspond to each auxiliary population’s GWAS summary statistics, and not the single-ancestry PRS. MPP-GWAS thus aggregates the effect sizes of SNPs from each auxiliary population’s summary statistics before proceeding with the optimization as per Equations 6.

###### MultiPopPred-GWAS-Admixture

Like MPP-GWAS, this version of MultiPopPred also requires the target *X* and *Y* along with auxiliary GWAS summary statistics as inputs. In this version, however, the auxiliary populations are given weights in accordance with their admixture proportions in the target population individuals. Suppose there are *K* auxiliary populations and *n* individuals in the target population. Then, for each individual, we compute the admixture proportions of the *K* auxiliary populations using an appropriate software such as ADMIXTURE [56]. (We employ the ADMIXTURE tool in a supervised manner with quasi-Newton convergence acceleration [57].)

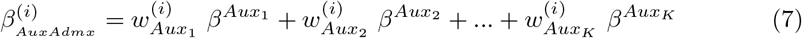

In Equation 7, 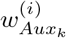 corresponds to the admixture proportion of auxiliary population *k* in the *i*^*th*^ target individual such that 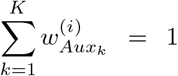. Furthermore, 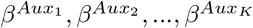 (each *m ×* 1) refer to each of the *K* auxiliary populations’ *β*s from their respective GWAS summary statistics. 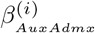 (*m ×* 1) refers to the admixed *β* for *i*^*th*^ target individual. The aggregated *β*^*Aux*^ is computed by taking the median across all *n* individuals in the target population as depicted in Equations 8 and 9 below.

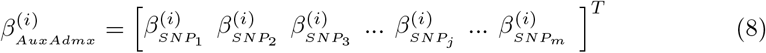

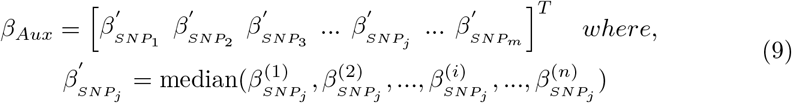

###### MultiPopPred-GWAS-TarSS

This version of MultiPopPred (MPP-GWAS-TarSS) is designed to take only the target and auxiliary populations’ summary statistics as input, along with an external target LD reference panel that may be derived from 1000 Genomes [29] (or any other appropriate resources). The intuition behind this version is to cater to the commonly encountered situation of non-availability of individual-level genotype and phenotype data for the target population due to scarcity, privacy, or ethical constraints. In such scenarios, MPP-GWAS-TarSS offers an efficient and fast (owing to Nesterov smoothing) alternative without a considerable loss in performance. Akin to the MPP-GWAS version, the aggregated *β*^*Aux*^ is computed as per Equation 5, giving equal weights to each auxiliary population.

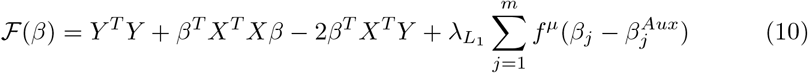

While this version also aims to optimize Equation 4 rewritten as Equation 10, an issue encountered is the source of *X* in *X*^*T*^ *X* and *X*^*T*^ *Y* being different (as pointed out by Mak et al. [26]). Since we do not have access to the actual target *X* and *Y, X*^*T*^ *X* is substituted by 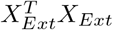 computed using an external LD panel (1000 Genomes) while *X*^*T*^ *Y* is substituted by the SNP-wise correlation (*r*) computed from the target population’s summary statistics using the formula *r* = (*n*_*Tar*_*β*^*Tar*^)_*m×*1_ (*n*_*Tar*_ referring to the number of samples used for target GWAS). We address this issue in a fashion similar to Mak et al. [26], by replacing 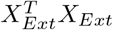 with 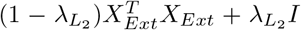 where *I* is an identity matrix and 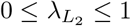. Ignoring the constant terms, Equation 10 can thus be rewritten as Equation 11, transforming our lasso optimization problem into an elastic-net one. The gradient vector supplied to L-BFGS also gets transformed as per Equation 12.

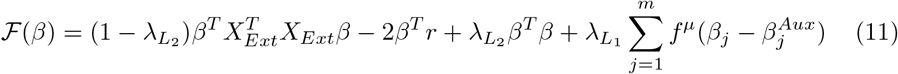

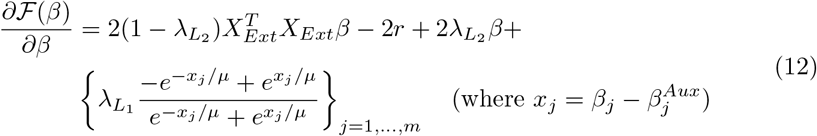

#### 4.1.5 MultiPopPred logistic module for binary phenotypes

MultiPopPred-PRS+ includes a separate module that implements a logistic model for binary phenotypes, as opposed to the default linear model used by the method. Assuming that the notations and dimensions of variables introduced earlier in Section 4.1.1 remain the same, the cost function ℱ (*β*) to be optimized is now defined as per Equation 13 below. While the penalization term remains the same as the linear module, here we make use of the cross-entropy loss as opposed to the least squares loss. Since the cross-entropy loss mandatorily requires the ground-truth *Y*, an equivalent external-LD version of MultiPopPred (similar to MPP-GWAS-TarSS) is not possible in this case. Other versions of MultiPopPred, however, continue to function exactly the same, with the new cost function depicted below.

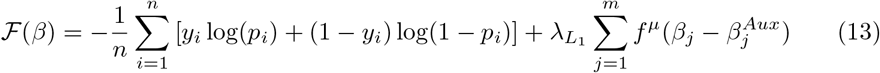

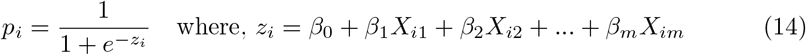

Here, *y*_*i*_ refers to the ground-truth phenotype value for the *i*^*th*^ target individual, and *X*_*ij*_ refers to the genotype value of the *i*^*th*^ individual at the *j*^*th*^ SNP. Additionally, the predicted probability *p*_*i*_ of a sample *i* belonging to class 1 (case) and the corresponding logit (or linear predictor) *z*_*i*_ are defined as per Equation 14. Let us collect these probabilities into a vector *P* = {*p*_*i*_} _*i*=1,…,*n*_. The first order gradient for ℱ (*β*) can then be written as per Equation 15 below (see Supplementary Methods 2.3.2 for a detailed gradient derivation).

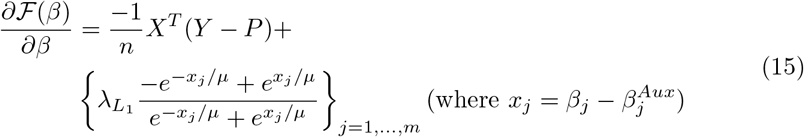

#### 4.1.6 MultiPopPred additional implementation details

MultiPopPred adopts a dynamic approach for the estimation of effect sizes of SNPs depending on the number of SNPs available as inputs. When the number of SNPs is small enough (*<* 30, 000), MultiPopPred processes the whole set of SNPs together as a single block. On the other hand, when the number of SNPs is large enough (*>* 30, 000), MultiPopPred processes the set of SNPs in blocks of a pre-set size. In our real-world analyses, the SNPs are processed in per-chromosome blocks (i.e., all SNPs in a chromosome constitute a single block). In our comparative analyses on PROSPER’s benchmark (Harvard Dataverse [43]), SNPs are processed in blocks of size 100 (i.e., each block containing 100 contiguous SNPs) per chromosome. These implementation decisions on block-wise processing have been made considering computational efficiency in mind, in line with the standard implementation of pipelines employed by PROSPER [21] and PRS-CSx [23]. Please note that in the standard implementation of all these methods, one regression model is fitted per block, and a final prediction is obtained by combining predictions from each of these models following a strategy as described in Supplementary Methods 2.1.

Algorithm 3 presented in Supplementary Methods 2.4 outlines the general termination criteria for the L-BFGS optimizer employed by MultiPopPred: reaching a maximum of 10^4^ iterations (`max_iter`), achieving a tolerance below 10^−4^ (`tol`), or exceeding 10^5^ maximum function evaluations (`max_fun_eval`), whichever is satisfied first. Throughout our simulation, semi-simulation, and real-world binary trait analyses we strictly applied the default `max_fun_eval` = 10^5^. However, for real-world quantitative trait analyses alone, we restricted `max_fun_eval` to 10. This adjustment served as a necessary computational safeguard against the substantial runtime overhead of computing the LD matrices (*X*^*T*^ *X*) required for the continuous trait objective function. This restriction, however, did not bias our reported results, as empirically evidenced from an additional sensitivity analysis across all 16 real-world traits using `max_fun_eval` thresholds of 10, 100, and 10^5^. Supplementary Data 24 depicts that the performance trends remained largely unchanged across all the different `max_fun_eval` settings.

### 4.2 Datasets and evaluation metrics used for benchmarking

#### 4.2.1 Generating simulated data

##### Genotypes

We used HAPGEN2 [30] to simulate individual-level genotypes for five different populations: South Asian (SAS), European (EUR), East Asian (EAS), American (AMR), and African (AFR). HAPGEN2 requires at minimum the following resources for simulations: (i) known haplotypes encoded as 0/1 (with one row per SNP and one column per haplotype) for a given reference population for resampling from, (ii) a legend file containing per SNP rsIDs, base pair positions, alternate and reference alleles, and (iii) a map file detailing fine-scale recombination rates across the given region. The known haplotypes for each of the five super-populations were obtained from 1000 Genomes [29, 58, 59] with SAS containing GIH samples, EUR containing CEU, FIN, GBR, IBS, TSI samples, EAS containing CDX, CHB, CHS, JPT, KHV samples, AMR containing CLM, MXL, PEL, PUR samples, and AFR containing ACB, ASW, LWK, YRI samples grouped together respectively. These SNPs were selected based on the availability of their respective genetic maps and recombination rates (Data availability). For a given super-population, the genetic maps and recombination rates were linearly interpolated at each position (Code availability), followed by averaging across the sub-populations and then proceeding with the final simulations. On consultation with the authors of PRS-CSx [23], who followed a similar workflow for data simulation, we noted that simulation of sub-population level genotype data followed by merging them into a super-population cohort would be an equally valid approach.

Although we simulated genotype data for multiple chromosomes (1-22 for SAS, and 19-22 for EUR, EAS, AMR, AFR), we have restricted all simulation analyses presented in Results section to only chromosome 22, in favor of computational efficiency. In total, we have 40,000 simulated samples available for SAS, and 50,000 available for each of EUR, EAS, AMR, and AFR. The set of SNPs used for simulation was restricted to the 1,297,432 HapMap3 variants [60], which were further subjected to a series of filters. The first two filters are standard and involve: (i) removal of triallelic and strand ambiguous variants, and (ii) retention of SNPs whose MAF *>* 1%. The third and final filter is a custom filter applied (again in favor of computational efficiency) to select SNPs in gene loci that are associated with diseases such as Type 2 Diabetes or Coronary Artery Disease, as reported in earlier GWAS of these diseases [9, 10, 13, 14]. This resulted in a final SNPs tally of 8215 in SAS, 9887 in EUR, 9353 in EAS, 9873 in AMR, and 9731 in AFR, all focused on chromosome 22. Please note that focusing only on these subsets of SNPs in chromosome 22, rather than all SNPs in the genome, is not expected to compromise the rigor/outcome of our simulation study. Because, our phenotype simulations rely on an infinitesimal model described below, wherein all selected SNPs (regardless of which chromosome they are in or which disease associations they exhibit) are treated similarly in terms of their contribution towards the simulated phenotype.

##### Continuous phenotypes

For simulating continuous phenotypes, we followed the linear model depicted in Equation 16 (an extension of Equation 1a, which also includes the heritability *h*^2^). We sampled effect sizes independently across SNPs. For each SNP, its (*K* +1) population-specific effect sizes (*β*) were jointly drawn from a (*K* +1)-variate multivariate Gaussian distribution having mean vector zero and permitting dependence across populations via the covariance matrix Σ. Here, we assume we have *K* auxiliary populations and one target population (*K* + 1 = 5 in our case corresponding to the 5 super-populations). Σ was designed to capture the correlation (*ρ*) of the effect size of a SNP between every pair of populations. For all major experiments, *ρ* (which we also call the inter-population correlation) was set to 0.8, unless otherwise specified.

In other words, Σ_*ij*_ = 0.8 if *i* ≠ *j* and Σ_*ij*_ = 1 if *i* = *j*, where *i, j* refers to auxiliary populations *i* and *j* respectively. The noise component of the phenotype was sampled from a zero-mean and unit-variance standard univariate Gaussian distribution. Finally, the contribution of the genetic components (SNPs) and the noise component (*ϵ*, including non-additive and environmental factors) towards the phenotype was controlled using a heritability parameter *h*^2^ ∈ [0, 1], as depicted in Equation 16. Here *X* corresponds to a (*n × m*) normalized genotype matrix 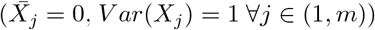, *β* is a (*m ×* 1) effect size vector, *ϵ* ~ *N* (0, 1) is the (*n ×* 1) noise vector. A higher value of heritability indicates a greater contribution of the genetic components towards the phenotype and vice versa. Standard/Baseline (per-SNP) GWAS was performed on the simulated data using `big_univLinReg`(`bigstatsr`) in R [61]. All scripts to perform data simulation as per the above description are available on our GitHub (Code availability).

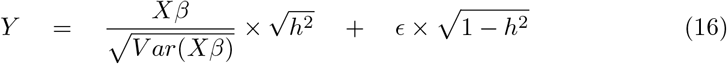

##### Binary phenotypes

For simulating binary phenotypes, we followed the standard liability threshold model [62, 63]. This model assumes that underlying a binary disease status (case versus control), there is a continuous, unobserved variable called liability. These liability phenotypes were simulated for each auxiliary and the target population using the exact same procedure for continuous phenotypes as described above. Next, we estimated a threshold *T* = Φ^−1^(1 Prevalence) for classifying the liability values into cases (1 − if liability *> T*) and controls (0 if liability ≤ *T*). Here Φ is the cumulative distribution function for a standard normal distribution. The prevalence of the binary traits was fixed to 20% for all simulations. Standard/Baseline (per-SNP) GWAS was performed on the simulated data using `plink --logistic --ci 0.95` flags. All scripts to perform data simulation as per the above description are available on our GitHub (Code availability).

#### 4.2.2 Generating semi-simulated data

##### Semi-simulated data

For semi-simulated analysis, we obtained real-world genotypes from UK Biobank. The 502,140 samples in UK Biobank were first grouped into subgroups based on self-identified ancestry (Datafield 21000). We further filtered out samples with any potential ambiguity in self-identified ancestry to retain 455,497 EUR (British and Irish) samples, 1571 EAS (Chinese) samples, 3390 AFR (African) samples, and 8015 SAS (Indian, Pakistani, and Bangladeshi) samples. The set of SNPs was restricted to the set of common SNPs across the four different ancestry groups (EUR, EAS, AFR, and SAS) and 1000 Genomes after application of appropriate filters such as MAF *>* 1% and removal of non-biallelic and strand-ambiguous variants. This resulted in a set of 5396 SNPs in chromosome 22 that was used for further analysis. Subsequently, continuous phenotypes were generated using the multivariate Gaussian approach as described above.

#### 4.2.3 Pre-processing real-world data

##### Real-world data

For real-world data analysis, we obtained real-world assayed genotypes from UK Biobank. All distinct samples were used in this study. Genotype encoding (0/1/2) followed the default encoding scheme as present in UK Biobank without any modifications. Eight quantitative phenotypes were obtained from UK Biobank: Standing Height, (ii) Body Mass Index, (iii) Systolic Blood Pressure, (iv) Diastolic Blood Pressure, (v) HDL Cholesterol, (vi) LDL Cholesterol, (vii) Total Cholesterol, and (viii) Total Triglycerides. Furthermore, eight binary phenotypes were obtained from UK Biobank - (i) Risk Taking, (ii) Educational Qualification, (iii) Allergy, (iv) Any CVD, (v) Asthma, (vi) Dyslipidemia, (vii) Morning Person, and (viii) Type 2 Diabetes. The corresponding data-field IDs, per population sample-sizes, and case-control distributions with prevalence for each of these traits have been detailed in Supplementary Data 10. All samples with missing values in the phenotypes were discarded. The missing values in the genotypes were imputed with simplistic approaches (mean across individuals, while ensuring that the standard deviation for each SNP remained *>* 0) owing to the fraction of missing values being pretty low (≤ 1%). The set of SNPs was restricted to those remaining post application of standard filters such as intersection with 1000 Genomes SNPs, MAF *>* 1%, and removal of non-biallelic and strand ambiguous variants. This resulted in a total number of samples and SNPs as reported in Supplementary Data 10-11. All genotype and (continuous) phenotype data were standardized (0 mean and unit variance), and the (continuous) phenotypes were adjusted for covariates such as Genetic Principal Components (PCs) 1-10 (Datafields 22009-a1 to 22009-a10), Age at recruitment (Datafield 21022), and Sex (Datafield 31). For binary phenotypes, the covariates were included as features in the joint logistic model. Individuals from UK Biobank were categorized into EUR, EAS, AFR, and SAS ethnic groups based on self-reported ethnicity (Datafield 21000). For all comparative analyses, the number of EUR samples was fixed at 50,000 due to computational resources’ limitations. For all continuous and binary traits, the target population, 1000 randomly selected samples were reserved for validation and another 1000 randomly selected samples for testing, while all other available samples were used for training. For 4 continuous traits, namely, Height, BMI, SBP, and DBP alone, another set of analyses was conducted with 2000 randomly selected validation and testing samples each to assess whether the variability in performance reduces with this increased test sample size. It was ensured that the set of training, validation, and testing samples did not overlap. For each trait, 3 sets of randomly shuffled datasets were used, and the performance was reported as the average over these datasets. (Per-SNP) GWAS, including GWAS done on each auxiliary population and Baseline GWAS done on the target population, was conducted on each real-world dataset using standard PLINK [64] procedures (`plink --assoc` for continuous traits and `plink --logistic –ci 0.95` for binary traits, both of which use Wald test results for reporting two-sided p-values). Supplementary Data Plots 25.1-25.20 depict the Manhattan and QQ-plots corresponding to the GWAS conducted on the 16 real-world continuous as well as binary traits. The QQ-plots report genomic inflation factor (scaled for 1000 samples) *λ*_1000_ values between 0.9 and 1.05 for all cases (except for TG GWAS conducted on AFR), indicating acceptable genomic inflation levels. The Manhattan plots, with thresholds set at p-value 5 *×* 10^−8^, further indicate that our GWAS pass standard quality checks.

#### 4.2.4 Evaluation metrics used in benchmarking

##### Evaluation metric (simulation and semi-simulation)

For comparing performances with respect to phenotype prediction in simulations as well as semi-simulations, we define correlation ratio as 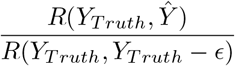, the ratio of the correlation between the true (ground-truth) and predicted phenotypes with the correlation between the true phenotype and the predicted phenotype if we somehow had access to the ground truth effect sizes (*β*_*Truth*_). Note that, *Xβ*_*Truth*_ = *Y*_*Truth*_ − *ϵ*. The correlation ratio gives us the ratio of how well a particular method is doing versus the best that could be done with respect to phenotype prediction. Ideally, we would want the correlation ratio to be as close to 1 as possible. A detailed explanation of correlation ratio as well as its relation to the standard metric *R*^2^ and heritability *h*^2^ is provided in Supplementary Methods 2.2.2.

##### Evaluation metric (continuous real-world traits)

For comparing performances with respect to phenotype prediction in continuous real-world traits, we have used *R*^2^, which is the square of the Pearson’s correlation between the true (ground-truth) and the predicted phenotypes. Ideally, we would want the *R*^2^ to be as close to the heritability of the trait as possible. Note that we could not use the correlation ratio since *β*_*Truth*_ is not known in real-world data. A detailed account of the evaluation strategy and the calculation of percentage improvement obtained (in terms of *R*^2^) by MultiPopPred over other SOTA methods has been provided in Supplementary Methods 2.2.1.

##### Evaluation metric (binary real-world traits)

For comparing performances with respect to phenotype prediction in binary real-world traits, we have used area under the precision-recall curve (`PR-AUC`) or the average precision. Additionally, we also report performances in terms of area under the ROC curve (ROC-AUC), while acknowledging its unsuitability in evaluating traits with highly imbalanced case-control distributions. A detailed account of the evaluation strategy and the calculation of percentage improvement obtained (in terms of both `PR-AUC` and Logit Variance) by MultiPopPred over other SOTA methods has been provided in Supplementary Methods 2.2.1.

### 4.3 State-of-the-art methods used for comparative analyses and their parameter settings

The various SOTA methods that were used for performing comparative analysis with MultiPopPred can be categorised broadly into three sets based on whether and how many auxiliary populations they make use of to estimate target effect sizes: (i) multiple auxiliary population-based methods, namely, SBayesRC-Multi, PROSPER, PRS-CSx, (ii) single auxiliary population-based methods, namely, TL-Multi, and (iii) solely target population-based methods namely, SBayesRC, Lassosum2, Baseline GWAS. All the above-mentioned methods were used either at their default hyperparameter values or were allowed to learn their own hyperparameter values from the input data, if applicable as described below.

#### SBayesRC

SBayesRC [20] is a Bayesian approach for single-ancestry PRS estimation that enhances PRS estimation by integrating GWAS summary statistics with functional genomic annotations through a resource-efficient low-rank model. SBayesRC employs a mixture of prior distributions corresponding to different magnitudes of causal SNP effect sizes and an MCMC framework to learn the posterior SNP effect sizes for a given population. The workflow of SBayesRC can be broken down into 3 major steps: (i) tidying GWAS summary statistics, (ii) imputing missing SNPs to match the reference LD panel, and (iii) the actual Bayesian mixture model estimation incorporating functional annotations. In this work, we use SBayesRC with SAS as the target population. The LD supplied to the method was computed from 1000 Genomes SAS individuals and has been made available through our GitHub (Data availability). SBayesRC was used at its default hyperparameter values namely, eigen variance cutoff thresholds (0.995, 0.99, 0.95, 0.9), starting *h*^2^ 0.5, starting *π* (0.99, 0.005, 0.003, 0.001, 0.001), *γ* (0, 0.001, 0.01, 0.1, 1), 3000 MCMC iterations (with 1000 burnin steps), ‘allMixVe’ method to resample residual, and 1.1 threshold to resample residual. Further, we used 96 functional annotations from the baseline model 2.2 [20] and the 1.2M SNP density panel [20] for all runs of SBayesRC in this work. It must be noted that this choice of 1.2M panel deviates from the authors’ recommendation of 7M SNP density panel which is derived from and suited for use with the UK Biobank imputed genotypes. The primary reason for our choice can be attributed to our use of UK Biobank assayed genotypes as opposed to imputed ones in favour of computational resource restrictions. We therefore encourage readers to interpret all comparative analyses presented in this work with this context in consideration.

#### SBayesRC-Multi

SBayesRC-Multi [20] is an extension of SBayesRC for cross-ancestry PRS prediction. In this framework, estimates of SNP effect sizes for each population are computed separately. Subsequently, a validation dataset from the target population is used to optimally combine individual PRS predictions into a final cross-ancestry prediction.

#### SBayesRC(O) and SBayesRC-Multi(O)

For certain simulation benchmarks (based on the infinitesimal model of phenotypes), the default starting *π* values of SBayesRC and SBayesRC-Multi could not achieve convergence. In such cases, an “oracle” version of these methods, SBayesRC(O) and SBayesRC-Multi(O) (that reveals some extra information about the Gaussian distribution of effect sizes in our simulations), was employed with a custom starting *π* (0.001, 0.679, 0.27, 0.04, 0.01). This starting *π* was decided by trying out several different sets of values that assumed differently distributed mixture of priors — (0.99, 0.005, 0.003, 0.001, 0.001), (0.7, 0.15, 0.08, 0.05, 0.02), (0.001, 0.249, 0.25, 0.25, 0.25) and (0.001, 0.679, 0.27, 0.04, 0.01). All other hyperparameters were used at their default settings as described above.

#### PROSPER

Polygenic Risk scOres based on enSemble of PEnalized Regression models was proposed by Zhang et al. in 2024 [21] as a PRS computation method that makes use of multiple populations to improve predictions on an understudied target population. The workflow can be broken down into three major steps. First, independent per population single-ancestry PRS computation is performed via Lassosum2 [21, 26] using training and validation datasets and the optimal parameters 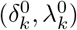 for each population *k* are stored.. The next step involves modeling multiple joint cross-population penalized regression tasks (corresponding to multiple populations) that make use of the single-ancestry PRS and optimal parameters 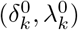 computed using Lassosum2 in the previous step. Finally, a super learning model is trained to compute an ensemble of PRS computed in the second step. In addition to 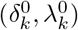, PROSPER uses 2 hyperparameters for tuning - *λ* (for controlling sparsity), and *c* (for controlling cross-ancestry effect similarity). The hyperparameter space spans a default 5 *×* 5 grid where *λ* takes 5 values spanning on a logarithmic scale from 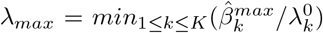 to *λ*_*min*_ = (0.001 ∗ *λ*_*max*_), with 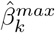 being the maximum absolute value from the GWAS summary statistics for population *k. c* is hard-coded to take values from (2, 8.01, 22.41, 50.74, 100).

#### PRS-CSx

Polygenic Risk Scores - Continuous Shrinkage was proposed by Ruan et al. in 2022 [23] as an extension to their previously published PRS-CS [65] single-ancestry PRS computation method. PRS-CSx makes use of global and local shrinkage parameters with only GWAS summary statistics from multiple populations and external LD reference panels per population to estimate per-population effect sizes via a Markov Chain Monte Carlo (MCMC) approximation strategy. PRS-CSx can be said to be following an approach, in theory, similar to an *L*_2_ penalized regression model. It has served as an established benchmark for comparing and contrasting newly developed PRS estimation methods over the years [25]. We ran PRS-CSx in the `auto` mode with its default hyperparameter values *a* = 1 and *b* = 0.5 in the gamma-gamma prior, with the global shrinkage parameter *ϕ* being learnt via a fully Bayesian approach from the input data. Other parameters, such as the number of burn-in and total MCMC iterations, were also used at the default values of 500\**K* and 1000\**K*, respectively, *K* being the number of discovery populations. The MCMC tinning factor was also used at a default value of 5.

#### TL-Multi

Transfer Learning - Multi was proposed by Tian et al. in 2022 [24] as a PRS estimation method that makes use of *L*_1_ penalization to minimize the difference between a single auxiliary population and a target population’s effect sizes. Similar to PROSPER, TL-Multi also makes use of Lassosum (Mak et al. [26]) as a subroutine to compute per-population GWAS followed by another instance of Lassosum to transfer learned effect sizes from the auxiliary to the target population. TL-Multi has also served as the basis for developing our MultiPopPred method. The publicly available code for TL-Multi was observed to have consistency issues (difference between their implemented and the paper’s objective function), and hence a corrected version of the same was used for all analyses in this work. The range of hyperparameter values (*L*_1_ regularization) employed by TL-Multi for grid search was used at the default levels of (0.2, 0.5, 0.9, 1).

#### Lassosum

Lassosum was proposed originally by Mak et al. in 2017 [26] as a single-ancestry PRS estimation method that utilizes *L*_1_ penalization, as suggested by the name. In this work, we use our own implementation of Lassosum (which employs the L-BFGS optimizer). We ran Lassosum under two configurations: Lassosum-TrueLD, which uses the true LD information (computed from individual-level *X* and *Y* data provided as inputs, to estimate effect sizes of the joint-SNP PRS model); and Lassosum-ExtLD, which uses the external LD reference panels (to convert the effect sizes of a per-SNP GWAS model provided as input to that of the joint-SNP PRS model). Lassosum-TrueLD and Lassosum-ExtLD use the same hyperparameters and hyperparameter space as MPP-PRS+ and MPP-GWAS-TarSS, respectively (see Supplementary Data 3).

#### Lassosum2

Lassosum2, which was implemented by Zhang et al. alongside PROSPER, was used in this work to facilitate our comparative analyses (especially to PROSPER). Both Lassosum2 and the original Lassosum [26] are the same in theory, and the only difference between them is the implementation. Lassosum2 uses coordinate descent for optimization and 2 hyperparameters - *λ* (*L*_1_ regularization, for controlling sparsity) and *δ* (*L*_2_ regularization, for controlling LD shrinkage). For each population, the hyperparameter space spans a default 5 *×* 5 grid, where *λ* takes 5 values spanning on a logarithmic scale from 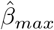 to *min*(0.001, 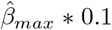), with 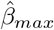 being the maximum absolute value from the GWAS summary statistics. And *δ* is hard-coded to take values from (0.5, 5.41, 20.07, 49.82, 100). Lassosum2 by default refers to Lassosum2-ExtLD, which uses external LD reference panels. In some specific analyses (such as Supplementary Results 1.2.2), we ran Lassosum2 using the true LD matrix instead of the external LD matrix, and therefore call it Lassosum2-TrueLD.

##### Baseline GWAS

This method refers to a simple GWAS performed on the target population training data without any additional information from other auxiliary populations. Like in any traditional GWAS, this method employs a simple linear regression-based approach without any penalization. We include this method in our comparative analysis to observe the relative improvements in performance that come with the inclusion of informative auxiliary populations. Baseline GWAS follows a per-SNP model as opposed to a joint all-SNP model employed for PRS estimation.

#### Clumping and Thresholding

This method refers to a clumping and thresholding applied to a simple GWAS (performed on the target population training data without any additional information from other auxiliary populations) to select subsets of SNPs for PRS computation. A total of 540 different configurations were used for this method corresponding to the following parameters - 9 p-value thresholds (5 *×* 10^−8^, 5 *×* 10^−7^, 5 *×* 10^−6^, 5 *×* 10^−5^, 5 *×* 10^−4^, 5 *×* 10^−3^, 5 *×* 10^−2^, 5 *×* 10^−1^, 1), 6 *r*^2^ thresholds (0.001, 0.01, 0.05, 0.1, 0.2, 0.5), 5 LD window thresholds (50kb, 100kb, 250kb, 500kb, 1000kb) and 2 reference genotypes (UK Biobank and 1000 Genomes). The best configuration was chosen based on a held-out validation dataset.

## Supporting information

Supplementary Information (Supplementary Results/Methods/Tables/Figures/DataFiles)

## 4.4 Data availability

Data generated in this study have been made available through https://bit.ly/4cHWaI3. MultiPopPred’s Supplementary Data files are available for access here: http://bit.ly/46SC61y. Data used in this study have been referred to appropriately: Access to the UK Biobank data was obtained under the project number 221732 from https://www.ukbiobank.ac.uk/; 1000 Genomes Phase 3 reference panel and genotypes: https://mathgen.stats.ox.ac.uk/impute/1000GP_Phase3.html, https://cncr.nl/research/magma/; Genetic maps for each sub-population as per 1000 Genomes: https://ftp.1000genomes.ebi.ac.uk/vol1/ftp/technical/working/20130507_omni_recombination_rates/; HAPNEST: https://www.ebi.ac.uk/biostudies/studies/S-BSST936; PROSPER’s Harvard Dataverse Simulations: https://dataverse.harvard.edu/dataset.xhtml?persistentId=doi:10.7910/DVN/COXHAP, Eigen-decomposed SAS LD (1000 Genomes as well as UK Biobank, separately) will be made available via: https://github.com/BIRDSgroup/MultiPopPred.

## 4.5 Code availability

The code developed in this study has been made available via the following GitHub link: https://github.com/BIRDSgroup/MultiPopPred. Codes of other relevant tools developed in other studies are accessible as follows: HAPGEN2: https://mathgen.stats.ox.ac.uk/genetics_software/hapgen/hapgen2.html; Interpolation of genetic maps: https://github.com/joepickrell/1000-genomes-genetic-maps; PRS-CSx: https://github.com/getian107/PRScsx; PROSPER and Lassosum2: https://github.com/Jingning-Zhang/PROSPER; SBayesRC and SBayesRC-Multi: https://github.com/zhilizheng/SBayesRC/tree/main; TL-Multi: https://github.com/mxxptian/TLMulti; ADMIXTURE: https://dalexander.github.io/admixture/; PLINK: https://zzz.bwh.harvard.edu/plink/plink2.shtml.

## 5 Acknowledgments

We thank Tom Michoel, and members of our BIRDS research group for their valuable inputs during the course of this work. We also thank Brintha VP and Saish Jaiswal for their inputs on this manuscript. This work was supported by the Prime Minister’s Research Fellowship (PMRF) Grant SB21221945CSPMRF008892 awarded to RK and Wellcome Trust/DBT India Alliance Intermediate Fellowship Grant IA/I/17/2/503323 awarded to MN. We would also like to acknowledge the use of LLMs for copy-editing certain sentences of the manuscript, and for obtaining code for certain minor helper functions, which were subsequently manually verified.

