## Supplementary Information (Supplementary Results/Methods/Tables/Figures/DataFiles) for "MultiPopPred: A Trans-Ethnic Disease Risk Prediction Method, and its Application to the South Asian Population"

2.5 Methodological details pertaining to analyses using true LD versus external LD . 17

**3 Supplementary Tables** **19**

**4 Supplementary Figures** **22**

**5 Supplementary Data Files** **44**

### 1 Supplementary Results

#### 1.1 Ablation studies and sensitivity analyses of MultiPopPred components

We conducted several ablation studies and sensitivity/perturbation analyses to systematically evaluate the choice we have made for the different components that make up MultiPopPred by quantifying their impact on performance. The following subsections elaborate on these analyses pertaining to the (i) optimizer, (ii) penalization strategy, (iii) smoothing function, and (iv) weighing scheme for the different auxiliary populations.

##### 1.1.1 On the choice of the optimizer

MultiPopPred employs the Limited-memory Broyden Fletcher Goldfarb Shanno (L-BFGS) [1] method as its optimization routine. L-BFGS is a quasi-Newton method that uses an estimate of the inverse Hessian matrix (instead of the full inverse Hessian) to explore the search space and locate an optimum, thereby making it an efficient choice for optimization problems involving a large number of parameters. Although L-BFGS is suited for optimization of smooth functions, its applicability to smoothed-versions of non-smooth functions has been well documented in the literature [2]. We elaborate more on that in the subsequent subsections. We specifically use the version of L-BFGS implemented in Python's SciPy library [3] via the `scipy.optimize.minimize()` functionality.

To compare the performance and utility of L-BFGS against traditionally employed optimization routines in disease risk estimation methods, namely Coordinate Descent (CD) [4], Proximal Gradient Descent (PGD) [5, 6], and Fast Iterative Shrinkage-Thresholding Algorithm (FISTA) [5, 6], we tested all these 4 optimization routines against simulated genotype-phenotype data. These simulations were performed under the default setting used throughout this work namely (i) the infinitesimal model assumption, with a $h^2$ of 70%, (ii) 4 auxiliary populations EUR, EAS, AFR, AMR with 10,000 samples each and a target population SAS with 1000 independent samples each for training, validation and testing, (iii) a value of 0.8 for the inter-population correlation (i.e., correlation between ground truth effect size $\beta$ vectors of a given pair of populations), and (iv) 8128 SNPs from chromosome 22 alone (assuming that these 8128 SNPs drive the entire genetic component of the simulated phenotype's heritability). Supplementary Figure 6 depicts that L-BFGS outperforms CD and FISTA while being better than or on-par with PGD in all five versions of MultiPopPred. To quantify the contribution of L-BFGS, replacing our L-BFGS optimizer with PGD leads to only an average 6% drop in performance, but replacing it with FISTA and CD optimizers leads to larger drops in average performance, namely 9% and 14%, respectively. Specifically, replacing L-BFGS with CD, PGD and FISTA respectively leads to a drop of 7%, -0.05%, 5% respectively on average in MPP-PRS+, 7%, 0.6%, 7% in MPP-PRS, 22%, 14%, 11% in MPP-GWAS, 12%, 0.02%, 9% in MPP-GWAS-TarSS, and 23%, 15%, 11% in MPP-GWAS-Admix.

Additionally, we examined several other optimization routines offered through Python's SciPy library against L-BFGS to convince ourselves about the non-availability of a better alternative. Supplementary Figure 7 depicts that L-BFGS has a better correlation ratio than the Nelder-Mead [3] as well as the Powell [3] optimization routines. Other solvers available via SciPy were not included due to either their failure to reach convergence or incompatibility with the optimization problem.

##### 1.1.2 On the penalization strategy

MultiPopPred aims to optimize a modified version of the traditional penalized least squares regression loss (or the penalized cross entropy loss in case of a logistic regression model for binary traits) wherein instead of penalizing just the target effect size estimates $\|\beta^{Tar}\|$, we penalize the difference/delta $\|\beta^{Tar} - \beta^{Aux}\|$ thereby forcing the target effect size estimates to be as close as possible to an aggregation of the auxiliary effect size estimates in proportion to the level of penalization. The choice of the $L_1$ penalization strategy in our model is motivated by the inherent property of $L_1$ penalization that drives features that are less important for prediction to have zero (or almost zero) delta effect size estimates, thereby also lending to the interpretability of the model. $L_2$ penalization, theoretically, is expected to be less stringent in terms of this feature selection. To empirically verify this hypothesis, we compared the performance of all five MultiPopPred versions with $L_1$ and $L_2$ penalization strategies, respectively, on simulated genotype-phenotype data. The simulation was done under the same assumptions as described in Supplementary Section 1.1.1. Supplementary Figure 8 depicts that employing the $L_1$ penalization strategy leads to a better performance in case of MPP-PRS, MPP-GWAS, and MPP-GWAS-Admix, while being on-par with $L_2$ penalization in case of MPP-PRS+, and MPP-GWAS-TarSS.

##### 1.1.3 On the smoothing function

The incorporation of $L_1$ penalization on the term $\|\beta^{Tar} - \beta^{Aux}\|_{L_1}$ in the cost function used by MultiPopPred makes it a non-smooth function, hence non-differentiable and difficult to optimize. We address this issue by employing Nesterov smoothing [7, 8] to obtain a smoothed approximation of the original cost function. To examine the impact of Nesterov smoothing on the predictive performance of MultiPopPred, we compared the performance of all five MultiPopPred versions with and without Nesterov smoothing on simulated genotype-phenotype data. The simulation was done under the same assumptions as described in Supplementary Section 1.1, except this time we had two configurations for target sample sizes: 100 and 1000. In the presence of Nesterov smoothing, the gradient was supplied to the L-BFGS optimizer, while in the absence of smoothing, the L-BFGS solver was allowed to estimate a numerical approximation of the true gradient through its internal mechanisms. Supplementary Figure 9 depicts that removing Nesterov smoothing alone from MultiPopPred leads to a 14% drop in performance averaged across all five versions and all simulated configurations. MPP-PRS+ reports the largest drop of 40% in average performance, while MPP-GWAS-TarSS reports a marginal average gain of 2% in performance. MPP-PRS, MPP-GWAS, and MPP-GWAS-Admix report drops of 7%, 12%, and 14%, respectively, in average performance without Nesterov smoothing.

##### 1.1.4 On the weighing scheme for auxiliary populations

All versions of MultiPopPred, except MPP-GWAS-Admix, employ an equal weighting scheme for the auxiliary populations. In other words, MPP-PRS+, MPP-PRS, MPP-GWAS, and MPP-GWAS-TarSS assign equal weights to the SNP effect size estimates coming from each auxiliary population. MPP-GWAS-Admix, on the other hand, assigns each auxiliary population a weight in accordance with its admixture proportion in the target population samples (see Methods). While the admixture weighting scheme might seem more intuitive theoretically, empirical observations in our analyses suggest that it offers no major advantage in performance over the equal weighing scheme in terms of phenotype prediction. Supplementary Figure 10 shows the admixture proportions of EUR, EAS, and AFR assigned to the target SAS samples across all 16 real-world (continuous as well as binary) traits studied in this work. EUR was seen to obtain the highest admixture proportion, followed by EAS and AFR. This is also in line with earlier published studies [9, 10], which report EUR, EAS, and AFR to be genetically closer to SAS, in that order (EUR>EAS>AFR), in terms of the $F_{st}$ metric. Consequently, MPP-GWAS-Admix unanimously awarded a higher weight to EUR, followed by a proportionate weight to EAS and AFR, respectively. However, in terms of performance on real-world traits, MPP-GWAS-Admix was observed to be on par, offering no significant improvement over its equal weighting employing counterparts MPP-GWAS and MPP-PRS+ (see Supplementary Figures 13 and 17).

To further verify the validity and soundness of our observations and MultiPopPred's implementation, we conducted simulation analyses to evaluate the five MPP versions under two additional simulation settings: (i) varying auxiliary sample sizes, and (ii) varying auxiliary-target genetic similarity. Both simulation settings were designed such that EUR got the highest weight, while other auxiliary populations got lower weights. For the first simulation setting, EUR, EAS, AMR, and AFR auxiliary populations were assigned sample sizes of 10000, 1000, 500, and 100, respectively, while the target SAS sample size was fixed at 100 and 1000. All other simulation parameters were fixed to be the same as described in Supplementary Section 1.1.1. For the second simulation setting, the inter-population correlation between SAS and EUR was fixed at 0.8, while that of EAS-SAS, AMR-SAS, and AFR-SAS was fixed at 0.2, 0.1, and 0.2, respectively. Similar to the previous setting, the target sample sizes were fixed at 100 and 1000. All other simulation parameters were fixed to be the same as described in Supplementary Section 1.1.1. For each of these simulation settings, three types of weighting schemes were compared: (i) equal weighting, (ii) admixture-based weighting, and (iii) sample size-based or genetic similarity-based weighting, as applicable. Supplementary Figure 11 depicts the comparative performance of all five MultiPopPred versions under these 3 weighting schemes. Under low target sample (100) settings, sample size-based weighting (or genetic similarity-based weighting, depending on the simulation setting) outperformed admixture-based weighting, which outperformed equal weighting. On the other hand, in the high target sample (1000) settings, all three weighting schemes were observed to be more or less comparable to each other across all MPP versions (except MPP-PRS+, for which equal weighting underperformed the other weighting schemes).

These observations indicate that in controlled and near-ideal settings (like our simulations), equal weighting may not be the best choice. Nevertheless, for real-world traits, the optimal weighting to be assigned to each auxiliary population is rarely driven solely by the sample size, admixture proportion, or genetic closeness to the target population. In our tests, equal weighting acted as a robust regularizer, preventing the model from becoming overly dependent on a single large or "close" auxiliary population (thereby mitigating erroneous transfer of potential biases in the effect sizes of the auxiliary population to the effect sizes of the target population). In such real-world scenarios, each auxiliary population's optimal weight may be determined by a complex interplay of the above and several other unobserved factors, which seemingly renders little to no difference when using equal weighting versus another weighting scheme (at least based on our real-world performance comparison of MPP-GWAS versus MPP-GWAS-Admix).

#### 1.2 Robustness of MultiPopPred under varying input data scenarios

##### 1.2.1 Varying degrees of similar versus dissimilar auxiliary and target populations

MultiPopPred was observed to perform better than or on par with state-of-the-art (SOTA) methods in simulations under a default setting where the auxiliary and target populations were known to be genetically similar or close to each other. This genetic similarity or closeness was quantified in simulations through the covariance matrix $\Sigma$ that defines how strongly or weakly correlated the ground-truth SNP effect size values are between the auxiliary and target population pairs. In all our major simulations, this correlation value was fixed to be 0.8 for all auxiliary-target pairs (see Methods).

To test the robustness of MultiPopPred in scenarios where the auxiliary populations are all weakly or moderately correlated with the target population, all five versions of MultiPopPred were subjected to evaluation under six different simulation settings. Genotype-phenotype data were simulated under three different inter-population correlation settings (low 0.3, moderate 0.6, and high 0.8) and two different target sample size settings (low 100 and high 1000). All other simulation parameters were fixed to be the same as described in Supplementary Section 1.1.1. Supplementary Figure 5 depicts that MultiPopPred maintains its performance trends in the case of low and moderate inter-population correlations. Specifically, all five versions of MultiPopPred were seen to outperform the SOTA methods in the low target sample settings across low, moderate, and high inter-population correlation levels. In case of high target sample settings, PROSPER was observed to catch up, performing on-par or marginally better than MPP-PRS, MPP-GWAS, MPP-GWAS-TarSS, and MPP-GWAS-Admixture, while MPP-PRS+ still remained the best performer.

##### 1.2.2 On the use of true LD versus external LD

MultiPopPred advocates the use of individual-level genotype and, hence, the true LD to estimate reliable trans-ethnic PRS for a low-resource population. To understand the impact of using true LD versus external LD for estimating PRS, we conducted the following analyses on 8 real-world continuous traits from UK Biobank. For each trait, MultiPopPred and the two leading SOTA methods, namely, SBayesRC-Multi and PROSPER, were run under two different settings: (i) true LD setting, and separately (ii) external LD setting. In each setting, it was ensured that every method to be compared was supplied with the exact same input files for a fair comparison. Subsequently, each method was allowed to process these input files as per its own internal frameworks, and then each method's final outputs were compared. We note that despite ensuring an input-matched scenario for all three methods in both true LD and external LD settings, finer differences in terms of each method's implementations and design choices remain, which are elaborated in Supplementary Methods 2.5.

In the first setting (true LD), each method was allowed full access to individual-level target as well as auxiliary genotype and phenotype. This renders MPP-PRS+ (which uses Lassosum-TrueLD as its base layer and is MultiPopPred's default mode of operation), PROSPER under true LD (which uses Lassosum2-TrueLD as its base layer), and SBayesRC-Multi under true LD (which uses SBayesRC outputs as its base layer). True LD was computed from UK Biobank data for PROSPER and SBayesRC-Multi in the format expected by the respective methods using their own LD computation codes, available through their respective GitHub repositories [11, 12]. In the second case (external LD), each method was allowed access to only target and auxiliary GWAS summary statistics and an external reference LD panel (1000 Genomes) for the target and auxiliary populations denoted as **Ext**LD. This renders PROSPER under external LD (which uses Lassosum2-**Ext**LD as its base layer and is PROSPER's default mode of operation), SBayesRC-Multi under external LD (which uses SBayesRC outputs as its base layer and is SBayesRC-Multi's default mode of operation), and MPP-PRS-TarSS (which uses Lassosum-**Ext**LD as its base layer). Supplementary Data 18 and 19 provide the training, validation, and testing performances for all methods under the true LD and external LD settings respectively.

In the true LD setting (Supplementary Figures 14, 15), the performance trends were observed to remain largely in line with those in Figure 6, wherein all methods were run at their default input requirement scenarios. With true LD, we observed that MultiPopPred (i.e., MPP-PRS+) continued to report better performance than PROSPER for Height, BMI, and SBP while reporting comparable performances for DBP, LDL, and HDL, and lagging in the case of TC and TG. With respect to SBayesRC-Multi, MPP-PRS+ reported better performance for Height, BMI, and SBP; comparable performances for DBP and LDL, while being surpassed in the case of HDL, TC, and TG. In the external LD setting, however, MultiPopPred was seen to clearly lag behind SBayesRC-Multi and PROSPER for all traits except BMI, SBP, and DBP, for which performances were comparable. Supplementary Data 21 depicts the total number of SNPs provided as input to each method, and the total number of SNPs retained in the output by each method for both true LD and external LD settings. All three methods were observed to retain almost the same number of SNPs in the true LD setting, indicating that any performance trends observed are not confounded by the difference in the number of SNPs processed by a given method. In the external LD setting, however, a noticeable difference was observed in the number of SNPs processed by each method. Overall, all these observations indicate a higher effectiveness of MultiPopPred in utilizing the information hosted in true LD as compared to SBayesRC-Multi and PROSPER.

To understand which component of each method was contributing to the overall performance gains, we decomposed each method's performance into two parts: the base layer and the meta layer. Supplementary Figures 14 and 15 depict that the performance improvement of MultiPopPred (under true LD, i.e., its version MPP-PRS+) over PROSPER (under true LD) and SBayesRC-Multi (under true LD) came from gains in both base and meta layers over the two methods, implying contributions from both the input choice (true LD over external LD) and methodological choices (Nesterov smoothing, L-BFGS optimizer, etc.) of MPP-PRS+.

#### 1.3 A note on the computational efficiency of MultiPopPred

To evaluate the computational efficiency of MultiPopPred against SOTA methods, we compared MPP-PRS+, SBayesRC-Multi, PROSPER, and PRS-CSx using the Height phenotype from UK Biobank for a typical genome-wide run (i.e., considering SNPs across all chromosomes 1 to 22). We selected MPP-PRS+ as the representative version of our approach, as running time and memory usage are expected to be consistent across the MultiPopPred versions. All analyses were performed on an Intel Xeon Platinum 8180 CPU (2.50GHz) with 1TB RAM. To ensure a fair comparison, each method was restricted to five (out of 112) logical CPU cores. Running times were reported as the elapsed wall-clock time. Memory usage was reported as the peak RAM (Random Access Memory) usage. All measurements were made using the GNU Linux command: `/usr/bin/time -v taskset -c 0-4 bash your_script.sh`.

For each method, we report the running time required by their respective base layer as well as meta layer in Supplementary Data 23. MPP-PRS+ was observed to take $\sim 2.1$ hours overall for a complete genome-wide run ($\sim 30 - 33$ minutes for each of the 3 auxiliary populations using Lassosum in the base layer and $\sim 33$ minutes for the meta MPP layer). Similar running times were observed for SBayesRC-Multi ($\sim 2.52$ hours overall with $\sim 36 - 40$ minutes for each auxiliary population in the SBayesRC base layer and $\sim 1$ minute for the meta Multi layer) and PROSPER ($\sim 2.25$ hours overall with $\sim 19 - 20$ minutes for each auxiliary population in the Lassosum2 base layer and $\sim 57$ minute for the meta PROSPER layer). PRS-CSx reported the largest running time of $\sim 5.28$ hours overall.

Memory usage was reported as a whole for each method. MPP-PRS+ was observed to use $\sim 14$ GB of memory, which is expected given its use of individual-level data. SBayesRC-Multi, PROSPER, and PRS-CSx were observed to use significantly lower memory: $\sim 6.2$ GB, 3.4 GB, and 0.94 GB, respectively. These observations on running time and space usage indicate that MultiPopPred can be comfortably run on a workstation, and its computational resource requirements are largely comparable to SOTA methods.

To further analyze the performance of MultiPopPred alone as a function of the sample size and number of SNPs used, we report the observed running time for the meta-layer of MPP-

PRS+ under two different training sample sizes (2000 and 5000) in Supplementary Data 23. The number of SNPs used was increased progressively, one chromosome at a time, starting with 37,957 (chromosome 1) to 4,85,309 (chromosomes 1 to 22). The running time was observed to increase steadily from $\sim$ 17 minutes (chromosome 1) to $\sim$ 33 minutes (all chromosomes) in the case of 2000 samples and $\sim$ 25 to 48 minutes for 5000 samples.

### 2 Supplementary Methods

#### 2.1 A note on the prediction strategy used for MultiPopPred

MultiPopPred processes data in a per-chromosome fashion, i.e., it fits one model per chromosome to output per-chromosome SNP effect size estimates. These estimates can eventually be used to calculate per-chromosome phenotype predictions. These per-chromosome phenotype predictions can then be summed up to arrive at the final phenotype prediction. Summing the per-chromosome predictions to obtain the final phenotype prediction is also in line with several published works, such as [11, 12] before.

Let us assume we have our per-chromosome partitioned target population dataset $(X^{(k)}, Y)$, $k \in \{1, 2, ..., 22\}$. $X^{(k)}$ is a $(N \times M_k)$ sized matrix of genotypes, $N$ being the total number of samples available, $M_k$ being the total number of SNPs in chromosome $k$. $Y$ is a $(N \times 1)$ vector of ground-truth phenotypes. First we partition our dataset into training, validation and testing sets $(X^k_{Train}, Y_{Train}), (X^k_{Val}, Y_{Val}), (X^k_{Test}, Y_{Test})$, $k \in \{1, 2, ..., 22\}$. Here $X^{(k)}_{Train}, X^{(k)}_{Val}$, and $X^{(k)}_{Test}$ are $(N_{Train} \times M_k), (N_{Val} \times M_k)$, and $(N_{Test} \times M_k)$ sized genotype matrices respectively, with $N_{Train}, N_{Val}$, and $N_{Test}$ being the number of samples allotted to the training, validation, and testing sets respectively. Similarly, $Y_{Train}, Y_{Val}$, and $Y_{Test}$ are $(N_{Train} \times 1), (N_{Val} \times 1)$, and $(N_{Test} \times 1)$ sized ground-truth phenotypes corresponding to the training, validation and testing sets respectively. At this point, we keep the testing set aside and do not touch it until we want to report final performance evaluations for the method. We use the per-chromosome training sets for training the models for each chromosome and the validation sets for tuning the model hyper-parameters.

Post-training, let us suppose we have per-chromosome SNP effect size estimates $\hat{\beta}^{(k)}$, where $k \in \{1, 2, ..., 22\}$ and $\hat{\beta}^{(k)}$ is a vector of size $(M_k \times 1)$. We use these estimates to calculate per-chromosome phenotype predictions $\hat{Y}^{(k)}$. This can be done using `plink2 --bfile genotype --score score_file`. However, since `plink2` [13] outputs the averaged predicted phenotype across alleles, we need to multiply each averaged predicted phenotype $\hat{Y}^{(k)}_{Avg}$ with $2 * M_k$ to get $\hat{Y}^{(k)}$. These predictions can then be summed up across the 22 chromosomes to get the final predicted $\hat{Y}$ as per Equation 1 below.

$$\hat{Y} = \sum_{k=1}^{22} \hat{Y}^{(k)}_{Avg} * 2 * M_k \quad (1)$$

Once we have the final predictions, we can calculate the performance metrics as described in the following subsections.

#### 2.2 A note on the evaluation strategy used for MultiPopPred

##### 2.2.1 Evaluation in real-world datasets

In case of continuous or quantitative phenotypes, we report the performance in terms of adjusted $R^2$. Here, we assume that the phenotype has already been adjusted for covariates.

Specifically, in this work, we adjusted all quantitative phenotypes for Age, Sex, and Principal Components (PCs) 1-10 of the genotype matrix. $R^2$ can be computed as either the square of the Pearson correlation between the ground-truth phenotype and the predicted phenotype ($\texttt{cor}(Y_{True}, \hat{Y})^2$ in R statistical environment) or via fitting a linear model and inspecting its summary ($\texttt{summary(lm(}Y_{True} \sim \hat{Y}\texttt{))}$ in R).

In case of binary phenotypes, we report the performance in terms of (i) area under the ROC curve (ROC-AUC), and (ii) area under the Precision-Recall (PR) curve or the average precision (PR-AUC). For highly imbalanced datasets, PR-AUC is a better performance metric than ROC-AUC. In the presence of covariates, fitting a joint logistic regression model including all the covariates and $\hat{Y}$ is advised instead of pre-adjusting the phenotype for covariates (since pre-adjusting essentially turns the binary phenotype into a continuous one with a potential loss in information). A covariate-adjusted predicted phenotype $\hat{Y}_{Adj}$ can then be obtained from the logistic model and used for computation of PR-AUC or ROC-AUC. We call the R functions in Equations 2 and 3 below to implement this approach.

$$\begin{aligned} \texttt{model = glm(}&Y_{True} \sim Age + Sex + PC1 + ... + PC10 + \hat{Y}, \\ &\texttt{data=validation\_data, family='binomial')} \end{aligned} \tag{2}$$

$$\hat{Y}_{Adj} = \texttt{predict(model, test\_data, type='link')} \tag{3}$$

To calculate the percentage improvement obtained by MultiPopPred over a SOTA method, we use Equations 4, 5, and 6. Here, $\phi$ is the cumulative distribution function of the standard normal distribution.

$$\left( \frac{R^2_{MPP} - R^2_{SOTA}}{R^2_{SOTA}} \right) * 100 \qquad \texttt{[For quantitative traits]} \tag{4}$$

$$\left( \frac{\texttt{PR\_AUC}_{MPP} - \texttt{PR\_AUC}_{SOTA}}{\texttt{PR\_AUC}_{SOTA}} \right) * 100 \qquad \texttt{[For binary traits]} \tag{5}$$

$$\left( \frac{\phi^{-1}(\texttt{MPP ROC\_AUC}) - \phi^{-1}(\texttt{SOTA ROC\_AUC})}{\phi^{-1}(\texttt{SOTA ROC\_AUC})} \right) * 100 \qquad \texttt{[For binary traits]} \tag{6}$$

##### 2.2.2 Evaluation in simulated and semi-simulated datasets

Simulated and Semi-simulated datasets offer us the unique advantage of having ground truth SNP effect sizes $\beta^{(k)}_{True}$, $k \in \{1, 2, ..., 22\}$ available. We use these ground truth SNP effect sizes to estimate an upper bound on how well any method could do with respect to the prediction of phenotypes. We define correlation ratio (CR), a custom metric, to evaluate the performance of methods on simulated and semi-simulated datasets (Equation 7). CR can be defined as the ratio of two correlations $C_1$ and $C_2$. Here, $C_1$ refers to the Pearson correlation between the ground-truth phenotype $Y$ and the method's predicted phenotype $\hat{Y}$. On the other hand, $C_2$ refers to the Pearson correlation between the ground-truth phenotype $Y$ and the predicted phenotype $Y_G = \sum_{k=1}^{22} X^{(k)} \beta^{(k)}_{True} = (Y - \epsilon)$ if the method had access to the ground truth SNP effect sizes through an oracle (here, $\epsilon$ refers to the ground-truth noise component of the ground-truth phenotype).

$$CR = \frac{C_1}{C_2} = \frac{\texttt{cor}(Y, \hat{Y})}{\texttt{cor}(Y, Y_G)} = \sqrt{\frac{R^2}{h^2_{SNP}}} \tag{7}$$

The relation between $CR$ and $R^2$ can be explained as follows. Let the ground-truth phenotype $Y$ be the sum of two components (i) the genetic component $Y_G$, and (ii) the environmental or noise component $Y_E = \epsilon$. The numerator ($C_1$) of CR can be calculated as the square root of a traditional $R^2$ metric for a phenotype prediction: $C_1 = \sqrt{R^2} = \sqrt{\texttt{cor}(Y, \hat{Y})^2}$. The denominator ($C_2$) of CR can be derived as per Equations 8 and 9 as follows.

$$C_2 = \texttt{cor}(Y, Y_G) = \frac{\texttt{Cov}(Y, Y_G)}{\sqrt{\texttt{Var}(Y)}\sqrt{\texttt{Var}(Y_G)}} \tag{8}$$

Now, $\texttt{Cov}(Y, Y_G) = \texttt{Cov}(Y_G + \epsilon,\ Y_G) = \texttt{Cov}(Y_G, Y_G) + \texttt{Cov}(\epsilon, Y_G) = \texttt{Var}(Y_G) + 0 = \texttt{Var}(Y_G)$, where we've used $\texttt{Cov}(\epsilon, Y_G) = 0$ due to $Y_G$ and $\epsilon$ being independent random variables. Therefore,

$$C_2 = \frac{\texttt{Var}(Y_G)}{\sqrt{\texttt{Var}(Y)}\sqrt{\texttt{Var}(Y_G)}} = \sqrt{\frac{\texttt{Var}(Y_G)}{\texttt{Var}(Y)}} = \sqrt{h^2_{SNP}} \tag{9}$$

#### 2.3 Gradient derivation for linear and logistic modules of MultiPop-Pred

##### 2.3.1 Gradient derivation for linear module

The cost function $\mathcal{F}(\beta)$ to be optimized by MultiPopPred (MPP-PRS+, MPP-PRS, MPP-GWAS and MPP-GWAS-Admix) is defined as per Equation 10 below for the linear module. Although the notations and conventions remain the same as Methods 4.1.1 to 4.1.2, we repeat them here for easy reference. Let us assume we have $n$ samples and $m$ SNPs. Here, $Y$ refers to the ground-truth phenotype vector of dimension $(n \times 1)$. $X$ refers to the ground-truth individual level genotype matrix having dimension $(n \times m)$, $\beta$ is a vector of SNP effect sizes having dimension $(m \times 1)$, $\beta^{Aux}$ is vector of aggregated SNP effect sizes from the auxiliary populations again having dimension $(m \times 1)$, $\lambda_{L_1}$ is the $L_1$ penalty/regularization parameter, and $f^{\mu}(x)$ is the Nesterov smoothing function defined as per Equation 11 while $\mu$ is the smoothing parameter.

$$\mathcal{F}(\beta) = (Y - X\beta)^T (Y - X\beta) + \lambda_{L_1} \sum_{j=1}^{m} f^{\mu}(\beta_j - \beta_j^{Aux}) \tag{10}$$

$$f^{\mu}(x) = \mu \log\left(\frac{1}{2}e^{-x/\mu} + \frac{1}{2}e^{x/\mu}\right) \tag{11}$$

To calculate the first-order gradient $\frac{\partial \mathcal{F}(\beta)}{\partial \beta}$ let us break down $\mathcal{F}(\beta)$ into two parts: (i) the least squares part $h_1(\beta) = (Y - X\beta)^T (Y - X\beta)$, and (ii) the Nesterov smoothed penalization part $h_2(\beta) = \lambda_{L_1} \sum_{j=1}^{m} f^{\mu}(\beta_j - \beta_j^{Aux})$ where $\beta_j$ and $\beta_j^{Aux}$ correspond to the $j^{th}$ SNP. We derive the gradient of both these parts one by one as follows.

$$
\begin{aligned}
\frac{\partial h_1(\beta)}{\partial \beta} &= \frac{\partial}{\partial \beta}\left(Y - X\beta\right)^T\left(Y - X\beta\right) \\
&= \frac{\partial}{\partial \beta}\left(Y^T - \beta^T X^T\right)\left(Y - X\beta\right) \\
&= \frac{\partial}{\partial \beta}\left(Y^T Y - Y^T X\beta - \beta^T X^T Y + \beta^T X^T X\beta\right) \\
&= \frac{\partial}{\partial \beta}\left(Y^T Y - 2\beta^T X^T Y + \beta^T X^T X\beta\right) \\
&= \frac{\partial (Y^T Y)}{\partial \beta} - 2\frac{\partial(\beta^T X^T Y)}{\partial \beta} + \frac{\partial(\beta^T X^T X\beta)}{\partial \beta} \\
&= 0 - 2X^T Y + 2X^T X\beta \qquad \text{(Since, } Y^T Y \text{ is a constant)} \\
&= 2X^T X\beta - 2X^T Y
\end{aligned} \tag{12}
$$

$$
\begin{aligned}
\frac{\partial h_2(\beta)}{\partial \beta_j} &= \frac{\partial}{\partial \beta_j}\left(\lambda_{L_1}\sum_{\ell=1}^{m} f^{\mu}(\beta_\ell - \beta_\ell^{Aux})\right) \\
&= \lambda_{L_1}\frac{\partial f^{\mu}(x)}{\partial x}.\frac{\partial x}{\partial \beta_j} \qquad \text{(here, } x := \beta_j - \beta_j^{Aux}\text{)} \\
&= \lambda_{L_1}\frac{\partial}{\partial x}\left(\mu \log\left(\frac{1}{2}e^{-x/\mu} + \frac{1}{2}e^{x/\mu}\right)\right).\frac{\partial}{\partial \beta_j}(\beta_j - \beta_j^{Aux}) \\
&= \lambda_{L_1}.\mu\frac{\partial}{\partial x}\left(\log\left(\frac{1}{2}e^{-x/\mu} + \frac{1}{2}e^{x/\mu}\right)\right).1 \\
&= \lambda_{L_1}.\mu\frac{1}{\left(\frac{1}{2}e^{-x/\mu} + \frac{1}{2}e^{x/\mu}\right)}.\left(\frac{-1}{2\mu}e^{-x/\mu} + \frac{1}{2\mu}e^{x/\mu}\right) \\
&= \frac{\lambda_{L_1}.\mu}{\mu}.\frac{-e^{-x/\mu} + e^{x/\mu}}{e^{-x/\mu} + e^{x/\mu}} \\
&= \lambda_{L_1}\frac{-e^{-x/\mu} + e^{x/\mu}}{e^{-x/\mu} + e^{x/\mu}} \qquad \text{(recall, } x := \beta_j - \beta_j^{Aux}\text{)}
\end{aligned} \tag{13}
$$

We can collect these partial derivatives across all $j$ in Equation 13 into a vector to obtain the gradient of $h_2$, denoted as $\frac{\partial h_2(\beta)}{\partial \beta}$. Specifically, $\left(\frac{\partial h_2(\beta)}{\partial \beta}\right)_j = \frac{\partial h_2(\beta)}{\partial \beta_j}$. Adding this gradient $\frac{\partial h_2(\beta)}{\partial \beta}$ to the gradient $\frac{\partial h_1(\beta)}{\partial \beta}$ from Equation 12, we get:

$$
\frac{\partial \mathcal{F}(\beta)}{\partial \beta} = 2X^T X\beta - 2X^T Y + \left\{\lambda_{L_1}\frac{-e^{-x_j/\mu} + e^{x_j/\mu}}{e^{-x_j/\mu} + e^{x_j/\mu}}\right\}_{j=1,\ldots,m} \qquad \text{(where, } x_j := \beta_j - \beta_j^{Aux}\text{)} \tag{14}
$$

Similarly, the cost function for MPP-GWAS-TarSS is defined as per Equation 15 below. Here, $X_{Ext}^T X_{Ext}$ refers to the external LD reference having dimension $(m \times m)$, $r$ refers to the SNP-wise correlation $X^T Y$ computed from the target GWAS summary statistics as $r = (n_{Tar}\beta^{Tar})_{m\times 1}$

($n_{Tar}$ referring to the number of samples used for target GWAS), and $\lambda_{L_1}, \lambda_{L_2}$ referring to the $L_1$ and $L_2$ penalty parameters respectively while all other variables remain the same as before. The first-order gradient of Equation 15 can then be computed as per Equation 16 below.

$$\mathcal{F}(\beta) = (1 - \lambda_{L_2})\beta^T X_{Ext}^T X_{Ext}\beta - 2\beta^T r + \lambda_{L_2}\beta^T\beta + \lambda_{L_1}\sum_{j=1}^{m} f^{\mu}(\beta_j - \beta_j^{Aux}) \tag{15}$$

$$\begin{aligned}\frac{\partial\mathcal{F}(\beta)}{\partial\beta} &= 2(1-\lambda_{L_2})X_{Ext}^T X_{Ext}\beta - 2r + 2\lambda_{L_2}\beta + \\ &\left\{\lambda_{L_1}\frac{-e^{-x_j/\mu} + e^{x_j/\mu}}{e^{-x_j/\mu} + e^{x_j/\mu}}\right\}_{j=1,\dots,m} \qquad (\text{where } x_j = \beta_j - \beta_j^{Aux})\end{aligned} \tag{16}$$

##### 2.3.2 Gradient derivation for logistic module

The cost function $\mathcal{F}(\beta)$ to be optimized by MultiPopPred for the logistic module is defined as per Equation 17 below. We use the same notations and conventions as detailed in the previous Section 2.3.1. In addition, we have the predicted probability $p_i$ of a sample $i$ belonging to class 1 (case) and the corresponding logit (or linear predictor) $z_i$, which are defined as per Equation 18 below.

$$\mathcal{F}(\beta) = -\frac{1}{n}\sum_{i=1}^{n}[y_i\log(p_i) + (1-y_i)\log(1-p_i)] + \lambda_{L_1}\sum_{j=1}^{m} f^{\mu}(\beta_j - \beta_j^{Aux}) \tag{17}$$

$$p_i = \frac{1}{1+e^{-z_i}} \quad , \text{where } z_i = \beta_0 + \beta_1 X_{i1} + \beta_2 X_{i2} + \dots + \beta_m X_{im} \tag{18}$$

Similar to the linear module, to calculate the first-order gradient $\frac{\partial\mathcal{F}(\beta)}{\partial\beta}$ we can break down $\mathcal{F}(\beta)$ into two parts: (i) the cross entropy part $h_1(\beta) = -\frac{1}{n}\sum_{i=1}^{n}[y_i\log(p_i) + (1-y_i)\log(1-p_i)]$, and (ii) the Nesterov smoothed penalization part $h_2(\beta) = \lambda_{L_1}\sum_{j=1}^{m} f^{\mu}(\beta_j - \beta_j^{Aux})$ where $\beta_j$ and $\beta_j^{Aux}$ correspond to the $j^{th}$ SNP. We can derive $\frac{\partial h_2(\beta)}{\partial\beta}$ in the same way as described for the linear module previously in Supplementary Methods 2.3.1. The first-order gradient derivation for $h_1(\beta)$ is described as follows. Let us first derive the first-order gradient $\frac{\partial h_1(\beta)_i}{\partial\beta_j}$ for the $i^{th}$ sample with respect to the $j^{th}$ SNP.

$$\frac{\partial h_1(\beta)_i}{\partial\beta_j} = \frac{\partial h_1(\beta)_i}{\partial p_i}.\frac{\partial p_i}{\partial z_i}.\frac{\partial z_i}{\partial\beta_j} \tag{19}$$

$$
\begin{aligned}
\frac{\partial h_1(\beta)_i}{\partial p_i} &= \frac{\partial}{\partial p_i}\left[y_i \log(p_i) + (1-y_i)\log(1-p_i)\right] \\
&= \frac{y_i}{p_i} + \frac{(1-y_i)}{(1-p_i)}.\frac{\partial}{\partial p_i}(1-p_i) \\
&= \frac{y_i}{p_i} - \frac{(1-y_i)}{(1-p_i)} \\
&= \frac{y_i - p_i}{p_i(1-p_i)} \qquad (20)
\end{aligned}
$$

$$
\begin{aligned}
\frac{\partial p_i}{\partial z_i} &= \frac{\partial}{\partial z_i}\left(\frac{1}{1+e^{-z_i}}\right) \\
&= -1.\frac{1}{(1+e^{-z_i})^2}.\frac{\partial}{\partial z_i}(1+e^{-z_i}) \\
&= \frac{-1}{(1+e^{-z_i})^2}.(-e^{-z_i}) \\
&= \frac{1}{(1+e^{-z_i})}\frac{e^{-z_i}}{(1+e^{-z_i})} \\
&= \frac{1}{1+e^{-z_i}}\left(1 - \frac{1}{1+e^{-z_i}}\right) \\
&= p_i(1-p_i) \qquad (21)
\end{aligned}
$$

$$
\begin{aligned}
\frac{\partial z_i}{\partial \beta_j} &= \frac{\partial}{\partial \beta_j}(\beta_0 + \beta_1 X_{i1} + \beta_2 X_{i2} + ... + \beta_j X_{ij} + ... + \beta_m X_{im}) \\
&= X_{ij} \qquad (22)
\end{aligned}
$$

Combining $\frac{\partial h_1(\beta)_i}{\partial p_i}$, $\frac{\partial p_i}{\partial z_i}$, and $\frac{\partial z_i}{\partial \beta_j}$ from Equations 20, 21, and 22 respectively into Equation 19 we get,

$$
\begin{aligned}
\frac{\partial h_1(\beta)_i}{\partial \beta_j} &= \frac{\partial h_1(\beta)_i}{\partial p_i}.\frac{\partial p_i}{\partial z_i}.\frac{\partial z_i}{\partial \beta_j} \\
&= \frac{y_i - p_i}{p_i(1-p_i)}.p_i(1-p_i).X_{ij} \\
&= (y_i - p_i)X_{ij} \qquad (23)
\end{aligned}
$$

The first-order gradient for all samples with respect to the $j^{th}$ SNP can then be written as per Equation 24 below.

$$
\begin{aligned}
\frac{\partial \mathcal{F}(\beta)}{\partial \beta_j} &= \frac{\partial h_1(\beta)}{\partial \beta_j} + \frac{\partial h_2(\beta)}{\partial \beta_j} \\
&= \frac{-1}{n}\sum_{i=1}^{n}\frac{\partial h_1(\beta)_i}{\partial \beta_j} + \frac{\partial h_2(\beta)}{\partial \beta_j} \\
&= \frac{-1}{n}\sum_{i=1}^{n}(y_i - p_i)X_{ij} + \lambda_{L_1}\frac{-e^{-x_j/\mu} + e^{x_j/\mu}}{e^{-x_j/\mu} + e^{x_j/\mu}} \quad (\text{where } x_j = \beta_j - \beta_j^{Aux}) \\
&= \frac{-1}{n}(X_{\{:,j\}})^T(Y - P) + \lambda_{L_1}\frac{-e^{-x_j/\mu} + e^{x_j/\mu}}{e^{-x_j/\mu} + e^{x_j/\mu}} \quad (\text{where } x_j = \beta_j - \beta_j^{Aux}) \qquad (24)
\end{aligned}
$$

In vector notation, we can write Equation 24 as per Equation 25,

$$
\frac{\partial \mathcal{F}(\beta)}{\partial \beta} = \frac{-1}{n}X^T(Y - P) + \left\{\lambda_{L_1}\frac{-e^{-x_j/\mu} + e^{x_j/\mu}}{e^{-x_j/\mu} + e^{x_j/\mu}}\right\}_{j=1,\ldots,m} \quad (\text{where } x_j = \beta_j - \beta_j^{Aux}) \qquad (25)
$$

Unlike in the linear module, we do not have an external LD version (TarSS) for our logistic module since the cross-entropy loss used here mandatorily requires the ground-truth $Y$. Furthermore, the objective function of the logistic module is not linear in $\beta$, therefore making a TarSS version problematic in this case.

#### 2.4 MultiPopPred pseudocode

This section describes the working of MultiPopPred through three pseudocode modules pertaining to data loading, data pre-processing, and optimization tasks. While all notations and variables used in the pseudocode modules have been introduced wherever applicable within the pseudocode itself, it must be mentioned that a detailed elaboration on the exact input requirements and methodological details can be found in the main text, Figure 1, and Methods.

**Algorithm 1** MultiPopPred Data Loading Module

**procedure** LoadGenomicDataAndBetaAux($V$) ▷ Module 1: Data Loading

  **if** $V \in$ {MPP-PRS+, MPP-PRS, MPP-GWAS, MPP-GWAS-Admix} **then**

    Load target genotype $X_{(n \times m)}$, phenotype $Y_{(n \times 1)}$, and covariates $C_{(n \times c)}$

    **if** $V$ = MPP-PRS+ **then**
      Load auxiliary SNP effect-size estimates $(\beta^{Aux_1}, \beta^{Aux_2}, ..., \beta^{Aux_K})$ each $(m \times 1)$ sized, from single-ancestry Lassosum-TrueLD PRS
    **else if** $V$ = MPP-PRS **then**
      Load auxiliary SNP effect-size estimates $(\beta^{Aux_1}, \beta^{Aux_2}, ..., \beta^{Aux_K})$ each $(m \times 1)$ sized, from single-ancestry Lassosum2-ExtLD PRS
    **else if** $V \in$ {MPP-GWAS, MPP-GWAS-Admix} **then**
      Load auxiliary SNP effect-size estimates $(\beta^{Aux_1}, \beta^{Aux_2}, ..., \beta^{Aux_K})$ each $(m \times 1)$ sized, from GWAS summary statistics
    **end if**

    **return** $\mathcal{D} \leftarrow (X, Y, C, \beta^{Aux_1}, \beta^{Aux_2}, ..., \beta^{Aux_K})$

  **else if** $V \in$ {MPP-PRS-TarSS, MPP-GWAS-TarSS} **then**

    Load external target reference panel $X_{Ext}$ sized $(n' \times m)$, $n \neq n'$

    **if** $V$ = MPP-PRS-TarSS **then**
      // MPP-PRS-TarSS is the sixth version of MPP introduced to match inputs for
      // the analyses on true LD versus external LD (Supplementary Results 1.2.2)
      Load auxiliary SNP effect-size estimates $(\beta^{Aux_1}, \beta^{Aux_2}, ..., \beta^{Aux_K})$ each $(m \times 1)$ sized, from single-ancestry Lassosum-ExtLD PRS
    **else if** $V$ = MPP-GWAS-TarSS **then**
      Load auxiliary SNP effect-size estimates $(\beta^{Aux_1}, \beta^{Aux_2}, ..., \beta^{Aux_K})$ each $(m \times 1)$ sized, from GWAS summary statistics
    **end if**

    **return** $\mathcal{D} \leftarrow (X_{Ext}, \beta^{Aux_1}, \beta^{Aux_2}, ..., \beta^{Aux_K})$

  **end if**

**end procedure**

**Algorithm 2** MultiPopPred Data Pre-Processing Module

```
procedure PREPROCESSDATA(𝒟, V)                          ▷ Module 2: Pre-Processing

    Construct 𝒟_processed using:
    Identify set of common SNPs 𝒮 across all input populations
    Filter 𝒟 to include only SNPs ∈ 𝒮
    Standardize genotype X        ▷ Each column of X (corresponding to a SNP across n
  individuals) has mean 0 and variance 1
    Adjust phenotype Y for covariates                   ▷ For continuous traits only

    if V = MPP-GWAS-Admix then
        Load auxiliary population weights (w_1, w_2, ..., w_K)
         from admixture files
    else
        Set equal weights w_1 = w_2 = ... = w_K = 1/K
         for each auxiliary population
    end if
    Compute β^Aux = w_1 β^{Aux_1} + w_2 β^{Aux_2} + ... + w_K β^{Aux_K}

    return (𝒟_processed, β^Aux)

end procedure
```

**Algorithm 3** MultiPopPred Main Algorithm

**procedure** RunMultiPopPredBaseAndMetaLayer($V$) ▷ Module 3: Optimization

$\mathcal{D}_{raw} \leftarrow$ LoadGenomicDataAndBetaAux($V$)
$\mathcal{D}, \beta^{Aux} \leftarrow$ PreProcessData($\mathcal{D}_{raw}, V$)

Initialize seed $\beta \sim \mathcal{N}(0, 10^{-10})$
Initialize smoothing parameter $\mu = 0.1$
Initialize $\lambda_{L_1}$ penalty hyperparameter
Initialize $\lambda_{L_2}$ penalty hyperparameter ▷ For TarSS versions only

**parallel for** each chromosome *chr* $\in \{1, \ldots, 22\}$ **do**

**if** Continuous Trait **then**
**if** $V \in$ {MPP-PRS-TarSS, MPP-GWAS-TarSS} **then**

$$\mathcal{F}(\beta) = (1 - \lambda_{L_2})\beta^T X_{Ext}^T X_{Ext}\beta - 2\beta^T r + \lambda_{L_2}\beta^T\beta + \lambda_{L_1}\sum_{j=1}^{m} f^{\mu}(\beta_j - \beta_j^{Aux})$$

**else**

$$\mathcal{F}(\beta) = (Y - X\beta)^T (Y - X\beta) + \lambda_{L_1}\sum_{j=1}^{m} f^{\mu}(\beta_j - \beta_j^{Aux})$$

**end if**

**else if** Binary Trait **then**
Calculate $\forall_{i=1}^{n}\ z_i = \beta_0 + \beta_1 X_{i1} + \beta_2 X_{i2} + \ldots + \beta_m X_{im}$
Calculate $\forall_{i=1}^{n}\ p_i = \dfrac{1}{1 + e^{-z_i}}$

$$\mathcal{F}(\beta) = -\frac{1}{n}\sum_{i=1}^{n}\left[y_i \log(p_i) + (1 - y_i)\log(1 - p_i)\right] + \lambda_{L_1}\sum_{j=1}^{m} f^{\mu}(\beta_j - \beta_j^{Aux})$$

**end if**

// max_iter: maximum number of iterations
// tol: tolerance
// max_fun_eval: maximum number of function evaluations
Set termination criteria ($max\_iter = 10^4$) $\vee$ ($tol < 10^{-4}$) $\vee$ ($max\_fun\_eval = 10^5$)

$\hat{\beta}^* \leftarrow \arg\min_{\beta} \mathcal{F}(\beta)$ ▷ Using the L-BFGS Optimizer

**end parallel for**

**return** $\{\hat{\beta}_1^*, \ldots, \hat{\beta}_{22}^*\}$ ▷ Refer Supplementary Section 2.1 on prediction strategy for $\hat{Y}$

**end procedure**

#### 2.5 Methodological details pertaining to analyses using true LD versus external LD

In Supplementary Results 1.2.2, we detailed observations from a robustness analysis conducted on 8 real-world continuous traits from UK Biobank to understand the impact of using true LD versus external LD for estimating PRS by running each of the three methods MPP, PROSPER, and SBayesRC-Multi under two different input-matched settings: (i) true LD setting, and separately (ii) external LD setting. It must be noted that despite ensuring an input-matched scenario for all three methods in both true LD and external LD settings, finer differences in terms of each method's implementations and design choices remain (see Supplementary Table 4). Specifically, MultiPopPred processes SNPs in per-chromosome blocks (i.e., one block per chromosome) whereas PROSPER and SBayesRC-Multi employ per-LD block processing. MultiPopPred employs an L-BFGS optimizer, PROSPER uses coordinate descent while SBayesRC-Multi uses an MCMC approximation framework. MultiPopPred does not apply any additional filters to the input set of SNPs, other than considering only the set of common SNPs across all populations. PROSPER, on the other hand, performs filtering (i) to retain SNPs in common with the LD panel it uses, and (ii) to discard SNPs it deems non-informative through internal mechanisms. Similarly, SBayesRC-Multi performs filtering to retain SNPs in common with the LD panel and the functional annotation file it uses. We chose not to harmonize these finer differences across methods since they are fundamental to their respective designs. Tinkering with their methodological choices would not be faithful to their respective intended designs.

Here, it must also be clarified that there are two different versions of Lassosum being employed by the different methods as their base layers — (i) Lassosum2, which is PROSPER's implementation of Lassosum that uses a coordinate descent optimizer, and (ii) Lassosum-TrueLD/Lassosum-ExtLD, which is our implementation of Lassosum that uses the L-BFGS optimizer with true LD/external LD, respectively. Lassosum2 is designed to have some internal SNP filtering mechanisms whereas Lassosum-TrueLD/Lassosum-ExtLD do not perform any such filtering. We chose to use our implementations of Lassosum-TrueLD/Lassosum-ExtLD as the base layer for our MultiPopPred versions, as opposed to Lassosum2, due to the following two reasons. First, Lassosum2 is designed to filter SNPs in a way that suits PROSPER best, which operates on a set of common SNPs between each pair of populations. MultiPopPred, on the other hand, by default operates on a set of SNPs common across all auxiliary and target populations and not any single pair of auxiliary-target populations. Using Lassosum2 as the base layer for MultiPopPred in such a scenario would be unfair, leading to a massive reduction in the total number of common SNPs to be used as features for PRS prediction, often leaving multiple chromosomes with no SNPs in common across populations. Second, intervening in Lassosum2's script and ensuring that both Lassosum2 and hence PROSPER use the exact same set of SNPs as MultiPopPred for their respective PRS computations would be unfaithful to their methodology.

We further clarify our approach to compute base layer performance for each method in these analyses so as to compare them with the corresponding meta layer performances. Assuming $X_{Test}^{SAS}$ to be the target test data, and $\hat{\beta}^{EUR}, \hat{\beta}^{EAS}, \hat{\beta}^{AFR}$ to be the auxiliary populations' effect sizes estimated by the base layer of a given method, we can make three sets of phenotype predictions as:

$\hat{Y}_{Test}^{EUR-SAS} = X_{Test}^{SAS} \cdot \hat{\beta}^{EUR}$,
$\hat{Y}_{Test}^{EAS-SAS} = X_{Test}^{SAS} \cdot \hat{\beta}^{EAS}$, and
$\hat{Y}_{Test}^{AFR-SAS} = X_{Test}^{SAS} \cdot \hat{\beta}^{AFR}$;

and compute the corresponding $R^2$ between each set of predictions and the actual phenotypes. The base layer performance for the method is then computed as the average of these $R^2$ values (i.e., $R^2$ reported by the method's respective auxiliary baselines when applied as is to the target test dataset). Formally, each method's baseline performance was computed as follows.

$$R^2_{\texttt{BaseLayer MPP}} = \frac{(R^2_{\texttt{Lassosum EUR}} + R^2_{\texttt{Lassosum EAS}} + R^2_{\texttt{Lassosum AFR}})}{3}$$

$$R^2_{\texttt{BaseLayer PROSPER}} = \frac{(R^2_{\texttt{Lassosum2 EUR}} + R^2_{\texttt{Lassosum2 EAS}} + R^2_{\texttt{Lassosum2 AFR}})}{3}$$

$$R^2_{\texttt{BaseLayer SBayesRC-Multi}} = \frac{(R^2_{\texttt{SBayesRC EUR}} + R^2_{\texttt{SBayesRC EAS}} + R^2_{\texttt{SBayesRC AFR}})}{3}$$

#### 3 Supplementary Tables

Supplementary Table 1: Hyperparameter Tuning on Simulated Data for the Five MultiPopPred Versions. The optimal hyperparameter values of the methods were used in both simulation and semi-simulation analyses. For each method, the optimal $L_1$ hyperparameter value is chosen from among the set of values $\{5 \times 10^{-8}, 5 \times 10^{-5}, 5 \times 10^{-3}, 5 \times 10^{-2}, 0.1, 0.5, 1, 2, 3, 4, 5, 6, 7, 7.5, 8, 9, 10, 50, 100, 1000\}$. The optimal $L_1$ and $L_2$ hyperparameter values for MPP-GWAS-TarSS is chosen from among the combination of the set of $L_1$ values defined above and the set of $L_2$ values $\{5 \times 10^{-8}, 5 \times 10^{-5}, 5 \times 10^{-3}, 5 \times 10^{-2}, 0.1, 0.5, 0.9, 0.99\}$. Correlation ratio is used as the performance metric for selecting the optimal value. For all but one method, $L_2$ hyperparameter is not applicable (NA).

| Method | Optimal Hyperparameter Value ($L_1$ and $L_2$) |
|---|---|
| MultiPopPred-PRS+ | $L_1$: 7.5, $L_2$: NA |
| MultiPopPred-PRS | $L_1$: 7.5, $L_2$: NA |
| MultiPopPred-GWAS | $L_1$: 7.5, $L_2$: NA |
| MultiPopPred-GWAS-TarSS | $L_1$: 100, $L_2$: 0.05 |
| MultiPopPred-GWAS-Admix | $L_1$: 7.5, $L_2$: NA |

Supplementary Table 2: Evaluating MultiPopPred on PROSPER's Benchmarks (Harvard Dataverse). Benchmark 1 corresponds to a strong negative selection setting with fixed common SNP heritability and 1% causal SNPs, while Benchmarks 2 and 3 have 0.1% and 0.05% causal SNPs respectively.

| | Benchmark 1 $R^2$ | | | Benchmark 2 $R^2$ | | | Benchmark 3 $R^2$ | | |
|---|---|---|---|---|---|---|---|---|---|
| Methods | Train | Val | Test | Train | Val | Test | Train | Val | Test |
| Baseline GWAS | 0.818 | 0.027 | 0.024 | 0.831 | 0.028 | 0.028 | 0.821 | 0.022 | 0.019 |
| PROSPER | 0.568 | 0.165 | 0.044 | 0.501 | 0.209 | 0.069 | 0.470 | 0.195 | 0.084 |
| Lassosum-ExtLD | 0.792 | 0.010 | 0.004 | 0.695 | 0.004 | 0.005 | 0.764 | 0.005 | 0.002 |
| PRS-CSx | 0.787 | 0.027 | 0.028 | 0.771 | 0.077 | 0.064 | 0.748 | 0.084 | 0.091 |
| SBayesRC | 0.630 | 0.012 | 0.012 | 0.520 | 0.021 | 0.016 | 0.180 | 0.014 | 0.014 |
| SBayesRC-Multi | 0.301 | 0.053 | **0.051** | 0.278 | 0.094 | **0.082** | 0.161 | 0.086 | **0.104** |
| Lassosum-True LD | 0.792 | 0.022 | 0.021 | 0.861 | 0.026 | 0.029 | 0.881 | 0.019 | 0.016 |
| MPP-PRS+ | 0.499 | 0.033 | 0.039 | 0.707 | 0.043 | 0.047 | 0.619 | 0.031 | 0.030 |

Supplementary Table 3: Heritability of Real World Traits. $h^2$ values were computed via the LDSC [14] method by the respective cited resources. Note: The $h^2$ values for HDL, LDL, TC and TG were computed using the standard biochemistry based traits in UKBiobank whereas NMR metabolomics based traits were used in the analyses.

| | LDSC Heritability ($h^2$) Value | |
|---|---|---|
| Trait | EUR $h^2$ | SAS $h^2$ |
| Height | 0.542 [15] | 0.261 [15] |
| BMI | 0.227 [15] | 0.0956 [15] |
| SBP | 0.134 [15] | 0.054 [15] |
| DBP | 0.126 [15] | 0.053 [15] |
| HDL | 0.33 [16] | NA |
| LDL | 0.083 [16] | NA |
| TC | 0.112 [16] | NA |
| TG | 0.218 [16] | NA |
| RT | 0.086 [15] | 0.194 [15] |
| EDQ | 0.236 [15] | 0.074 [15] |
| ALG | 0.114 [15] | 0.074 [15] |
| Any CVD | NA | NA |
| AST | 0.107 [15] | 0.056 [15] |
| DLP | 0.127 [12] | NA |
| MP | 0.107 [15] | 0.052 [15] |
| T2D | 0.164 [15] | 0.124 [15] |

Supplementary Table 4: Methodological Divergence between MultiPopPred, PROSPER, and SBayesRC-Multi

| **Features of the method as well as the respective base layer** | **MultiPopPred** | **PROSPER** | **SBayesRC-Multi** |
|---|---|---|---|
| Baseline Framework | Lassosum (Our implementation) | Lassosum2 | SBayesRC |
| Optimizer | L-BFGS | Coordinate Descent | MCMC Approximation |
| LD Processing Unit | Per-chromosome blocks | Per-LD blocks | Per-LD blocks |
| SNP Filtering Strategy | No additional filters (Uses SNPs common across all populations) | Filters for LD panel matching and internal non-informative SNP removal | Filters for LD panel matching and functional annotation file consistency |
| Default Input Mode | True LD (Individual-level genotype) (Recall that this default version is MPP-PRS+) | External LD (GWAS Summary Statistics) | External LD (GWAS Summary Statistics) |
| LD Sensitivity: Extent of performance drop when moving from True LD to External LD | Significant | Modest | Modest |

### 4 Supplementary Figures

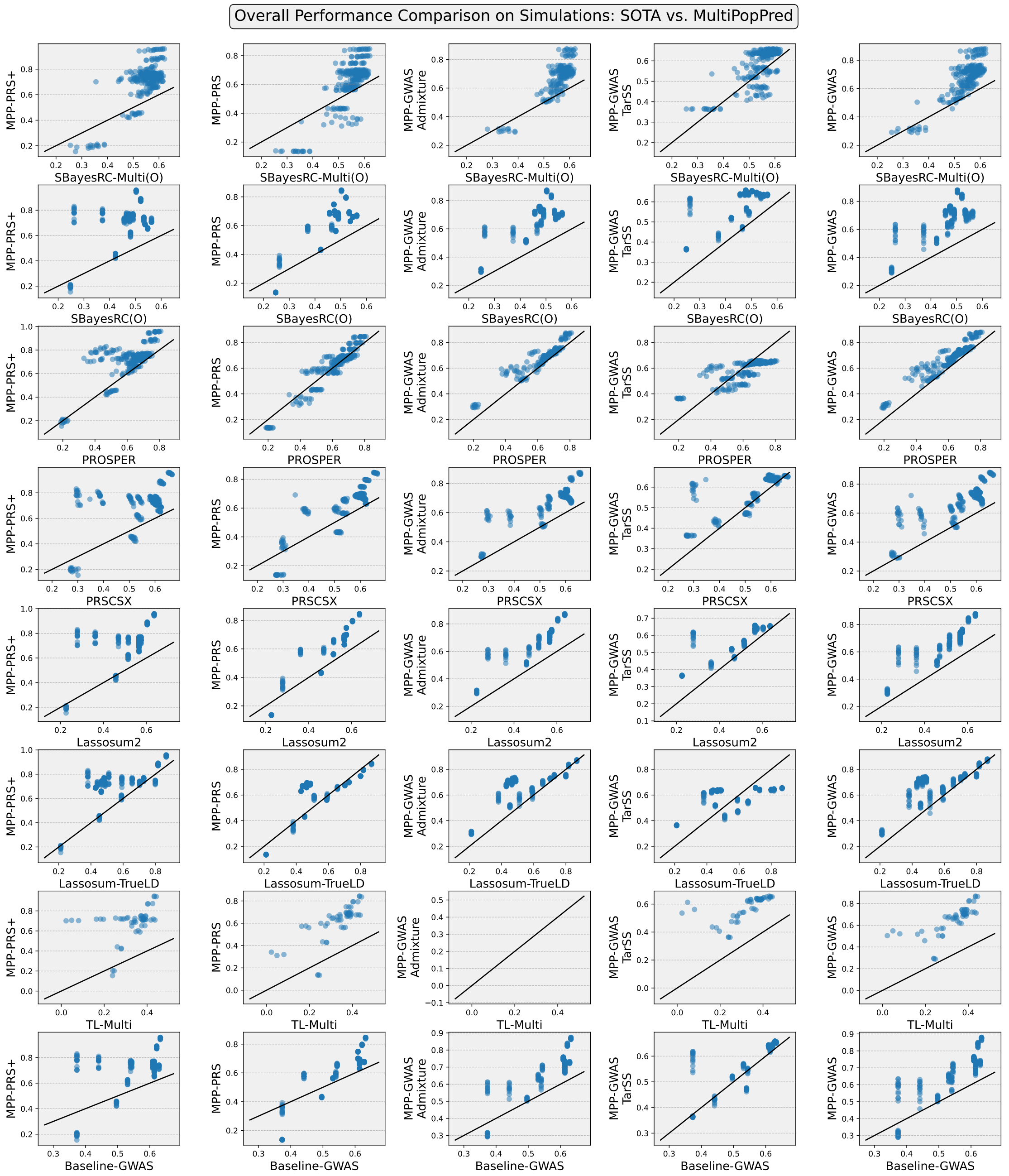

Supplementary Figure 1: Overall Performance Comparison on all Simulations. MPP-GWAS-Admixture versus TL-Multi does not have any points because these two methods do not share any common configurations. MPP-GWAS-Admixture requires at least 2 auxiliary populations to compute admixture proportions, while TL-Multi works with only a single auxiliary population.

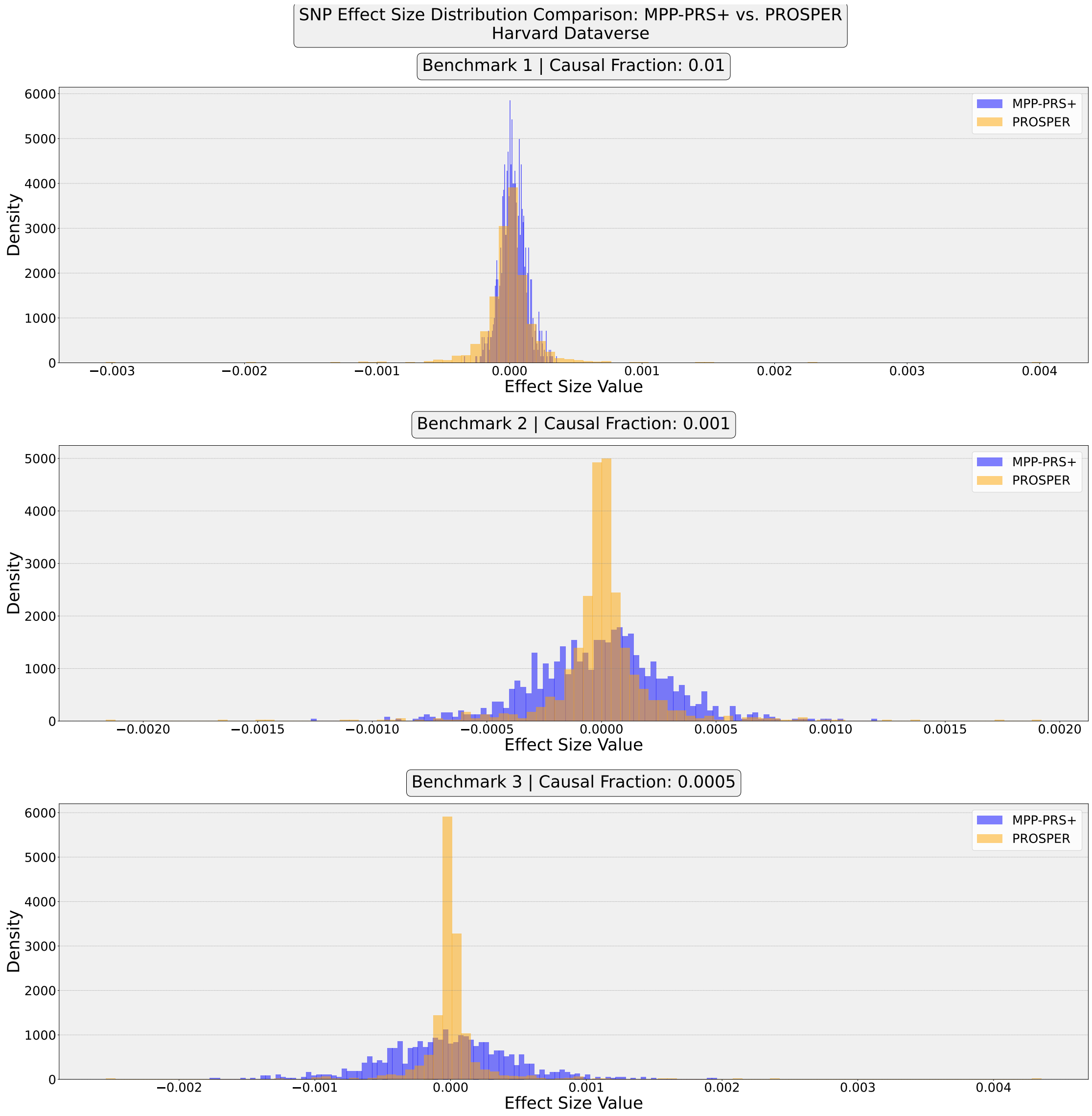

Supplementary Figure 2: Distribution of the predicted SNP effect sizes obtained by MPP-PRS+ and PROSPER, for three benchmarks from PROSPER's simulations in Harvard Dataverse.

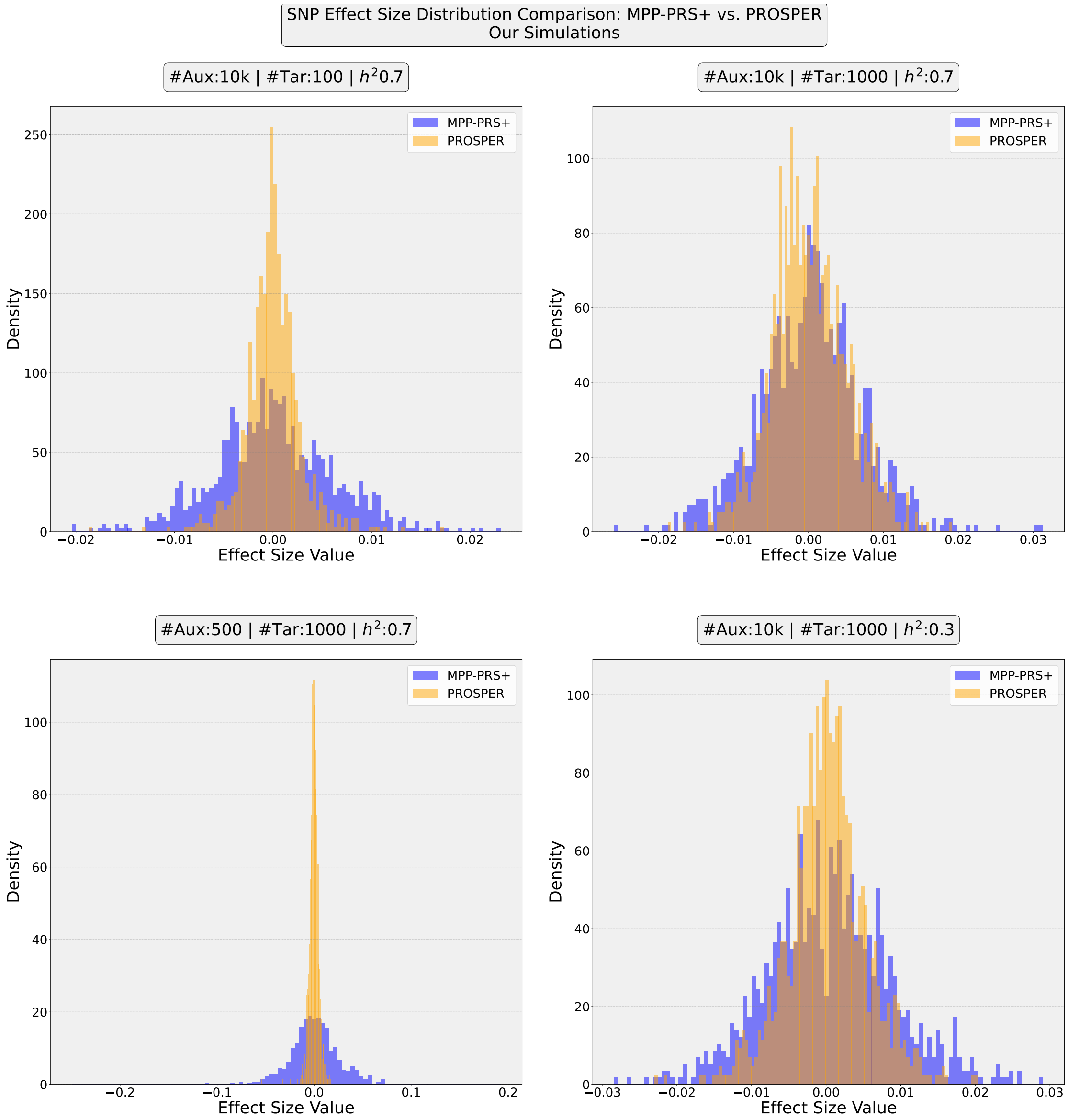

Supplementary Figure 3: Distribution of the predicted SNP effect sizes obtained by MPP-PRS+ and PROSPER, for four benchmarks from our simulations.

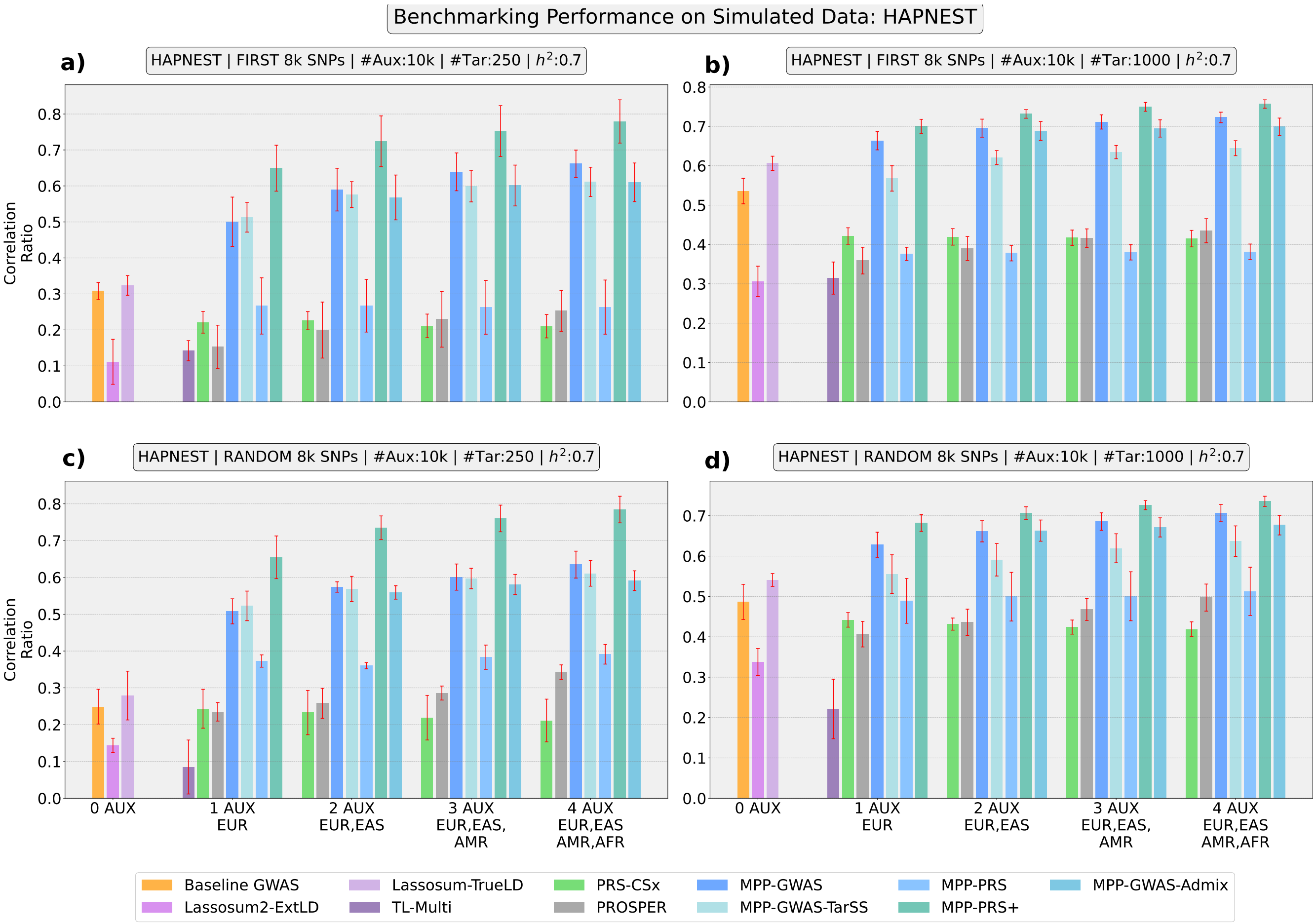

Supplementary Figure 4: Benchmarking Performance (Phenotype Prediction) on HAPNEST Simulated Data: We depict the comparative performance of MultiPopPred with SOTA methods under four different HAPNEST simulation settings and four auxiliary-target population tuples. In (A) and (B), a contiguous set of 8000 SNPs from chromosome 22 was selected for the analysis to observe the performance in the presence of LD structure in the SNP set. In (C) and (D), a random set of 8000 SNPs from chromosome 22 was selected for the analysis to observe the performances in the absence of LD structure in the SNP set. (A) The number of samples per auxiliary population was set to 10,000, while the number of target population samples was set to 250. $h^2$ was set to 70%. (B) The number of auxiliary samples and $h^2$ remained the same as in (A), but the number of target samples was increased to 1000. (C) The number of samples per auxiliary population was set to 10,000, while the number of target population samples was set to 250. $h^2$ was set to 70%. (D) The number of auxiliary samples and $h^2$ remained the same as in (C), but the number of target samples was increased to 1000. It must be noted that MultiPopPred-Admixture is not applicable under the single auxiliary population setting since it does not make sense to calculate admixture proportions for a single reference population. On the other hand, TL-Multi is not applicable for $> 1$ auxiliary population settings. Abbreviations: Aux: Auxiliary Population, Tar: Target Population, $h^2$: Heritability.

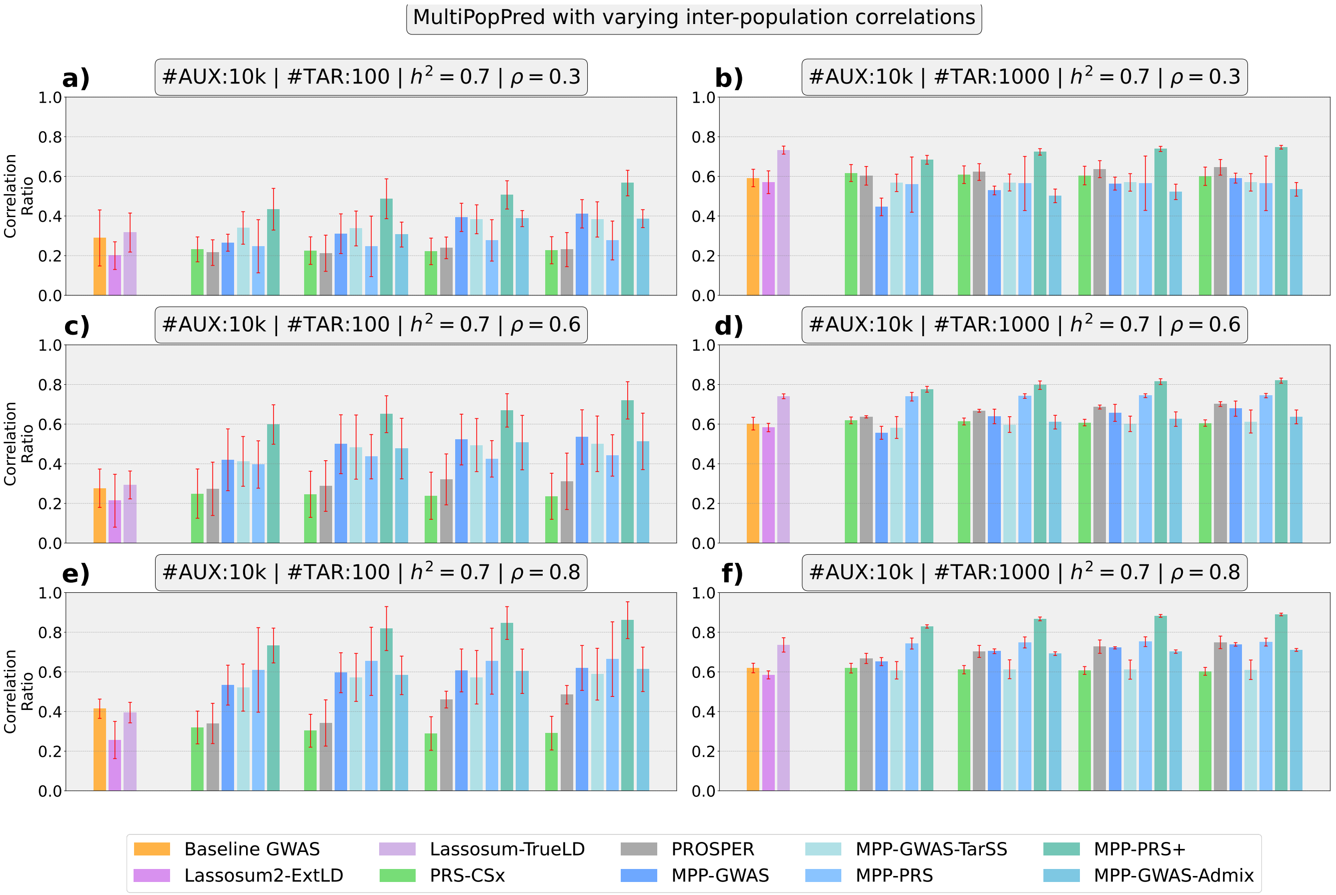

Supplementary Figure 5: Evaluating performance (Phenotype Prediction) of MultiPopPred under varying inter-population correlation values used for simulation. Abbreviations: Aux: Auxiliary Population, Tar: Target Population, $h^2$: Heritability.

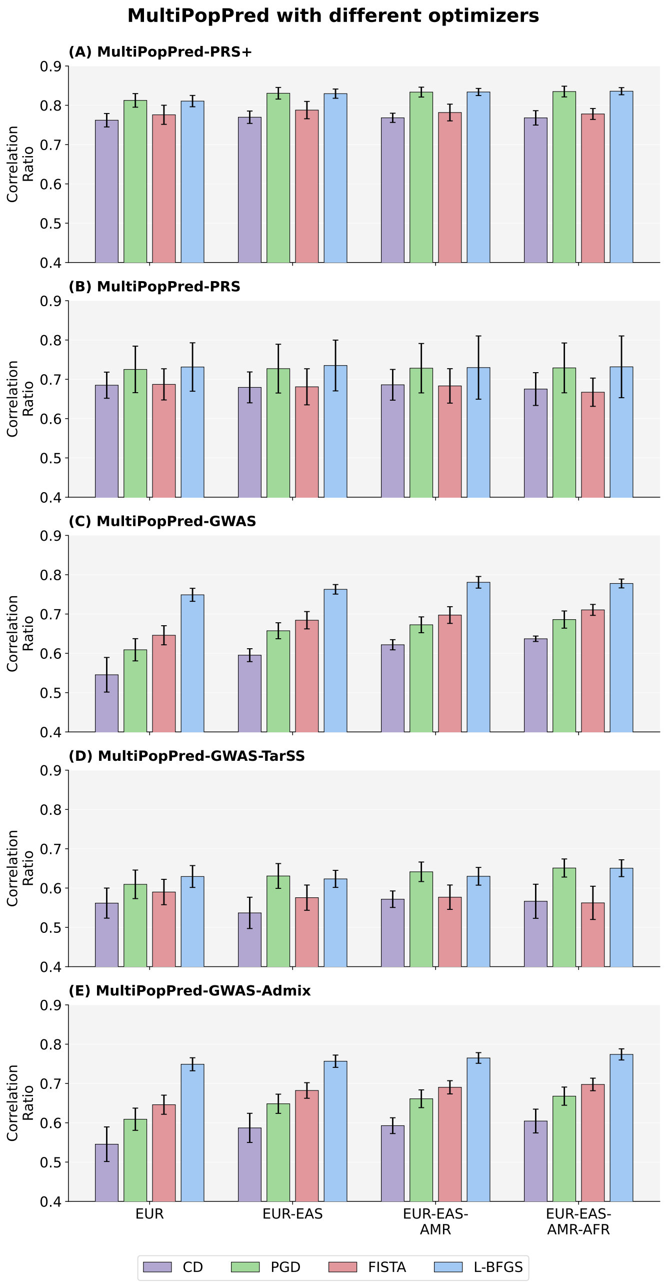

Supplementary Figure 6: Analysis to compare the predictive performance of MultiPopPred versions on simulations with different optimization routines, namely, Coordinate Descent (CD), Proximal Gradient Descent (PGD), Fast Iterative Shrinkage-Thresholding Algorithm (FISTA), and L-BFGS, using four auxiliary populations: EUR, EAS, AMR, and AFR. The number of samples per auxiliary population was set to 10,000, while the number of samples for the target (SAS) population was set to 1000. $h^2$ was set to 70%. The convergence or termination criteria for all four methods was fixed at 10,000 maximum iterations or 0.0001 tolerance or 100,000 maximum function evaluations (in case of L-BFGS only), whichever is reached first. The L-BFGS optimizer was observed to perform better than CD and FISTA, while being on-par with PGD.

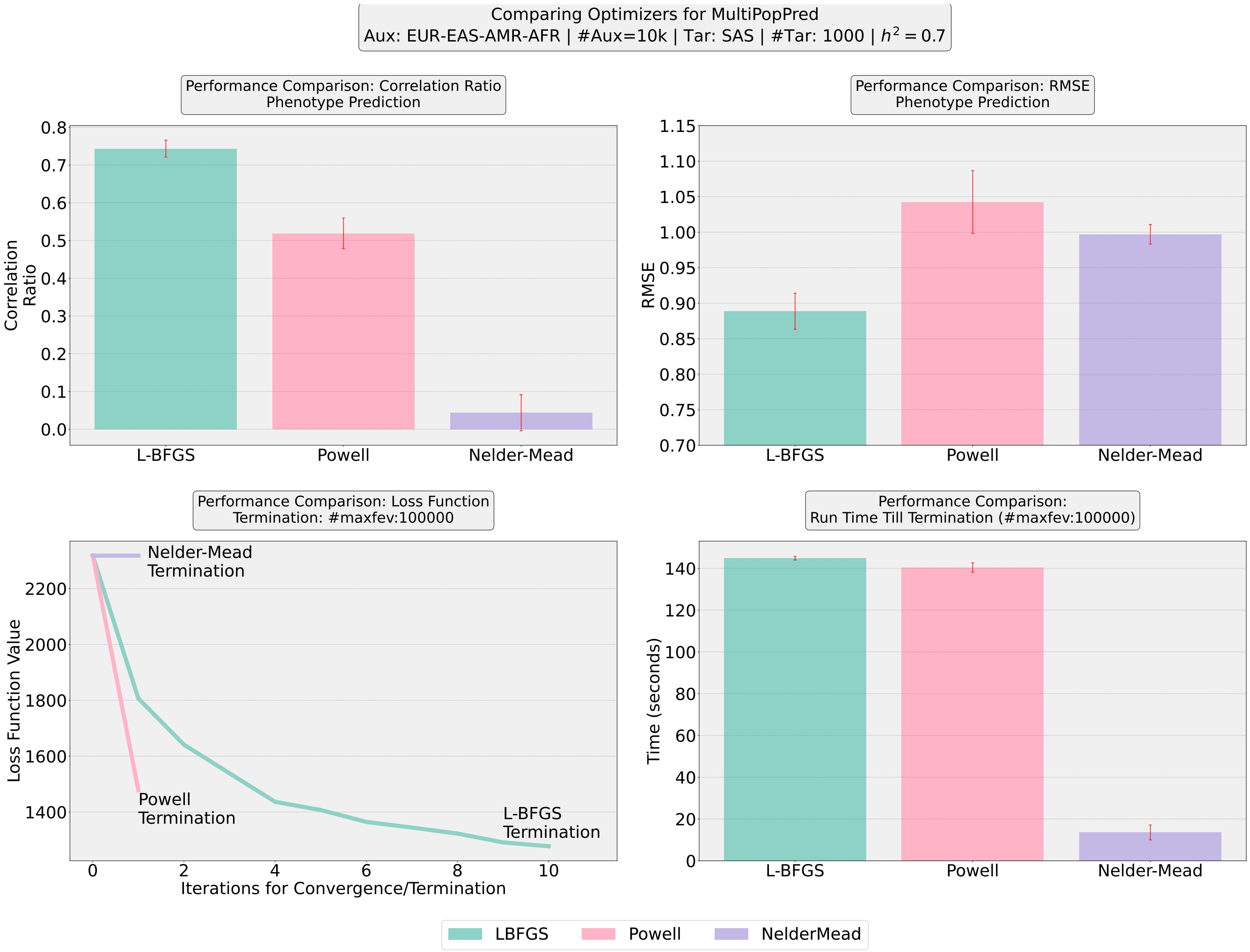

Supplementary Figure 7: Analysis to compare the predictive performance of MultiPopPred versions on simulations with different optimization routines using four auxiliary populations: EUR, EAS, AMR, AFR. The number of samples per auxiliary population was set to 10,000, while the number of samples for the target (SAS) population was set to 1000. $h^2$ was set to 70%. The convergence or termination criteria for all the 3 methods was the same as mentioned in Algorithm 3. The MultiPopPred variants were observed to perform better with an L-BFGS optimization routine as compared to Powell and Nelder-Mead.

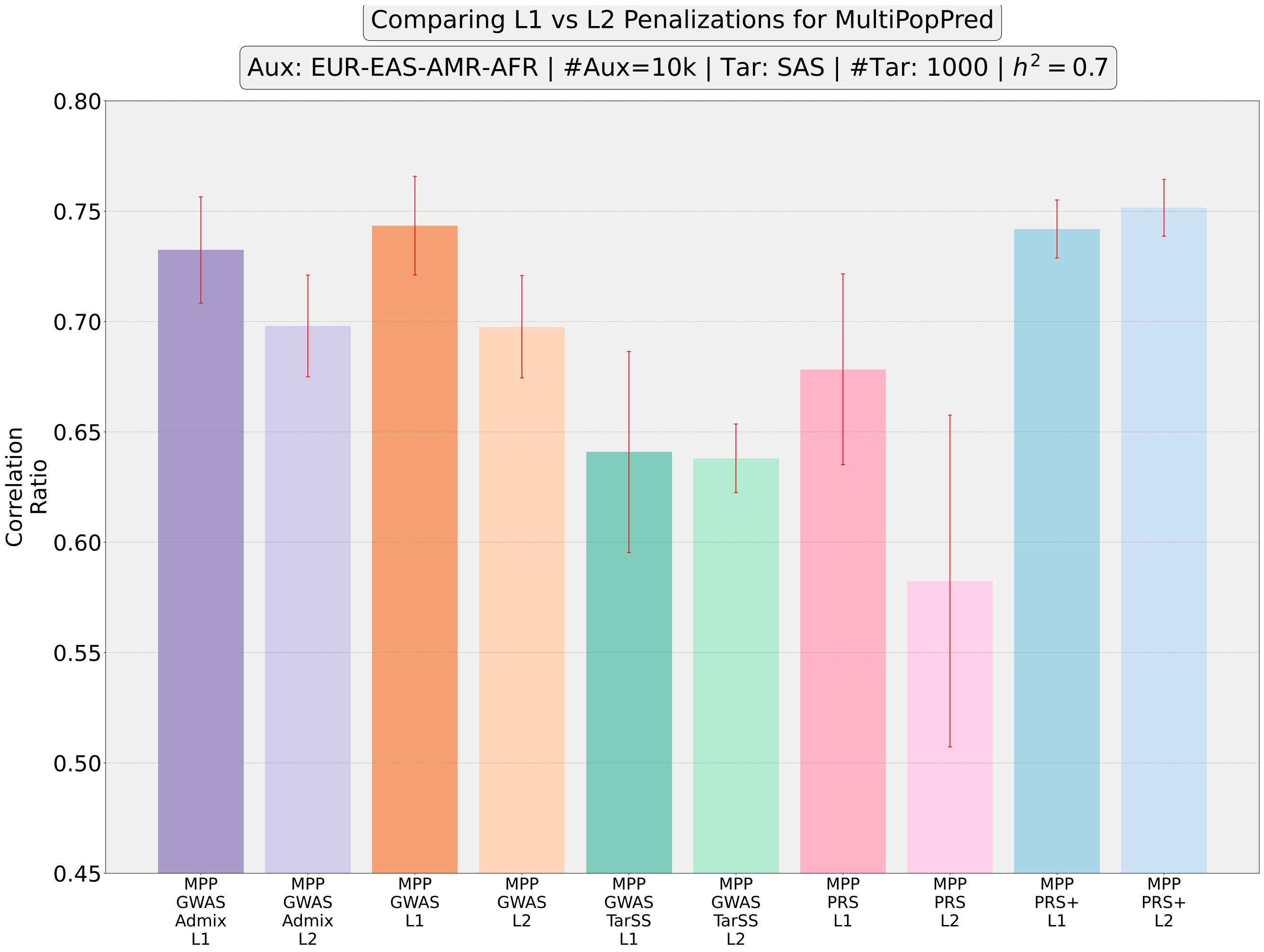

Supplementary Figure 8: Analysis to compare the predictive performance of MultiPopPred variants on simulations with $L_1$ versus $L_2$ penalization using four auxiliary populations: EUR, EAS, AMR, and AFR. The number of samples per auxiliary population was set to 10,000, while the number of samples for the target (SAS) population was set to 1000. $h^2$ was set to 70%. The MultiPopPred versions were observed to perform better with an $L_1$ penalization strategy as compared to an $L_2$ penalization strategy.

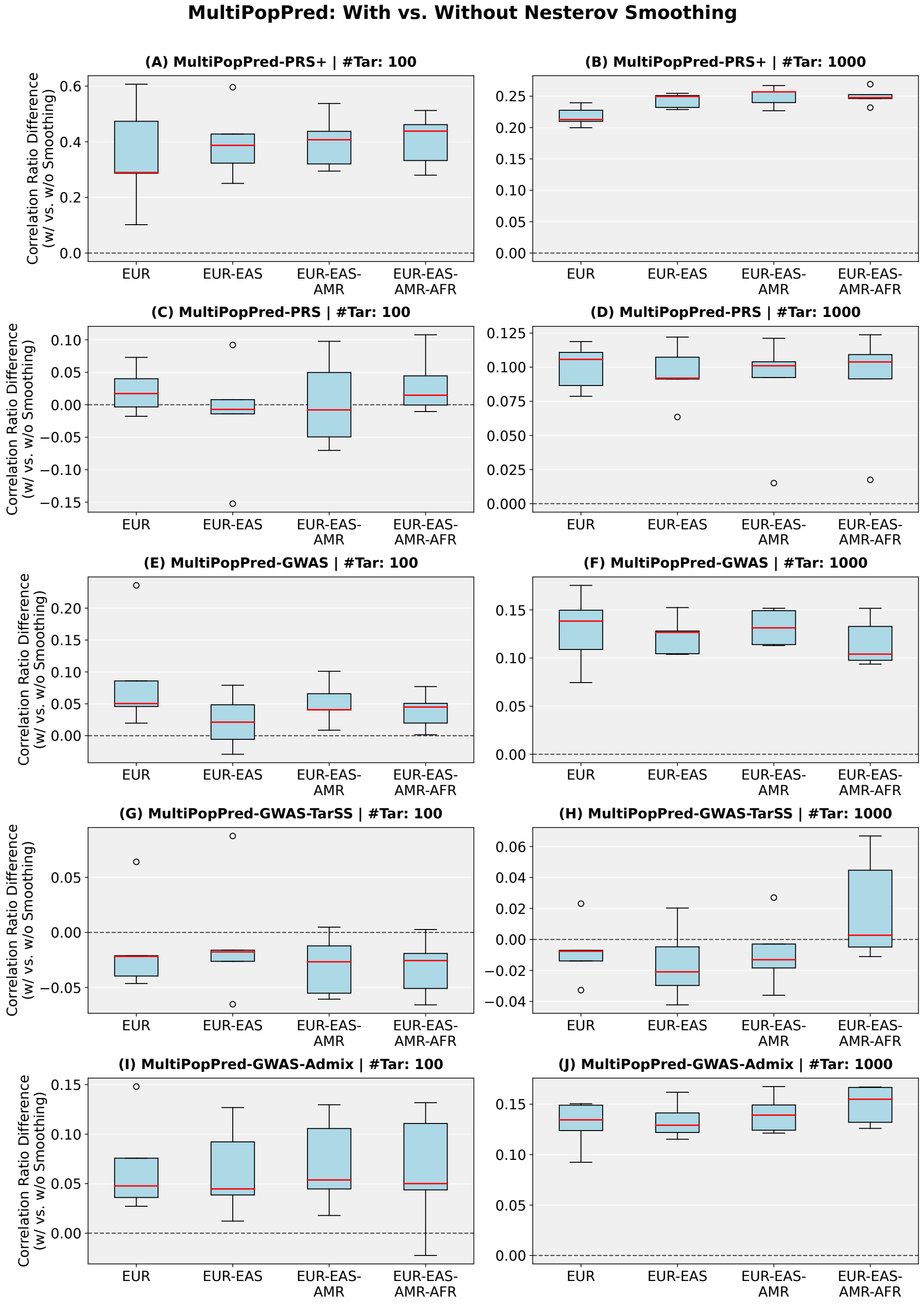

Supplementary Figure 9: Analysis to compare the predictive performance of MultiPopPred versions on simulations with and without Nesterov smoothing. The simulation included four auxiliary populations: EUR, EAS, AMR, and AFR, with the number of samples per auxiliary population being 10,000, while the number of samples for the target (SAS) population was set to 100 and 1000. $h^2$ was set to 70%. The MultiPopPred versions were observed to benefit from the inclusion of Nesterov smoothing of the cost function to be optimized.

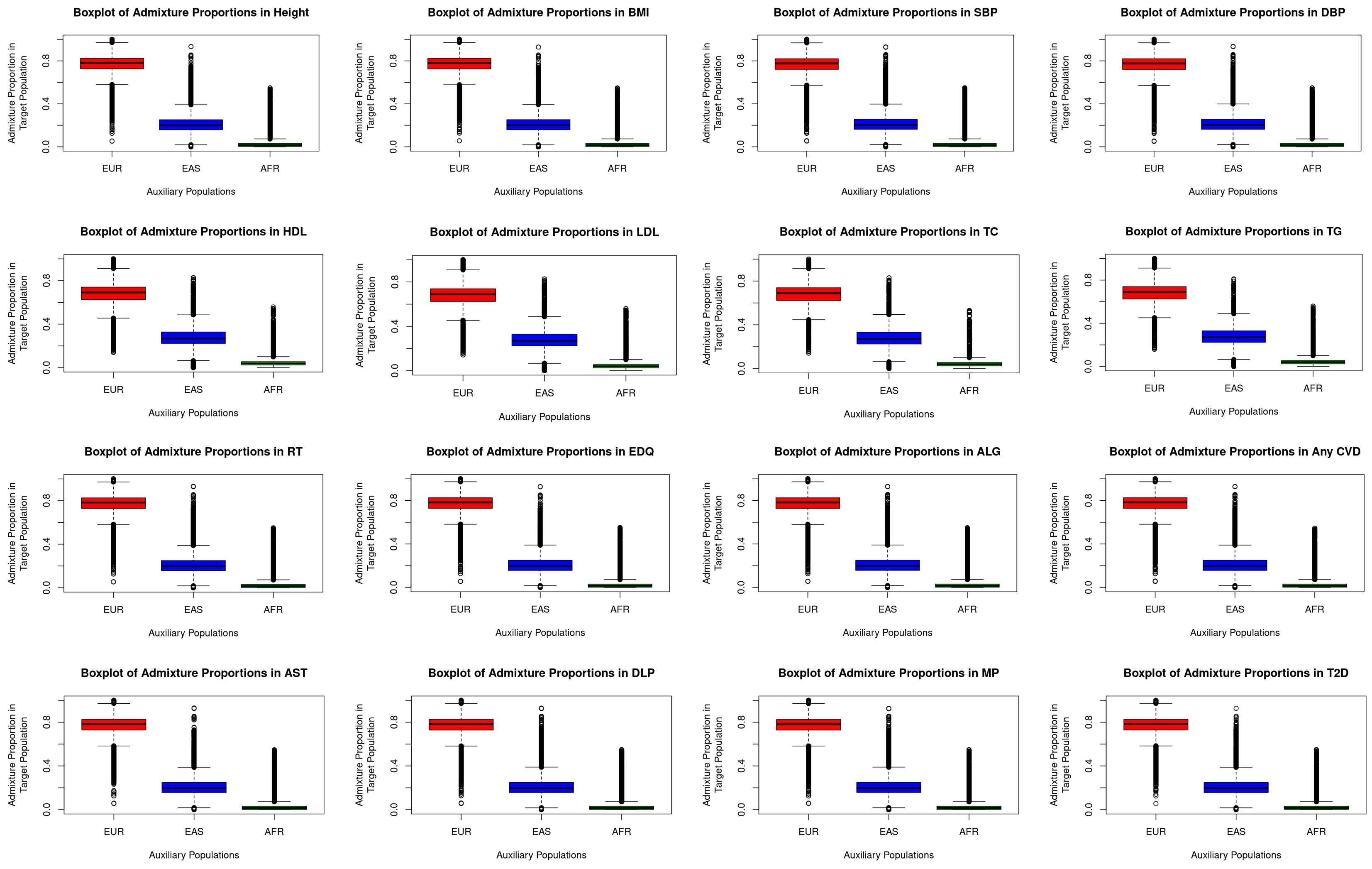

Supplementary Figure 10: Admixture proportions observed in SAS samples across 16 real-world traits from UK Biobank.

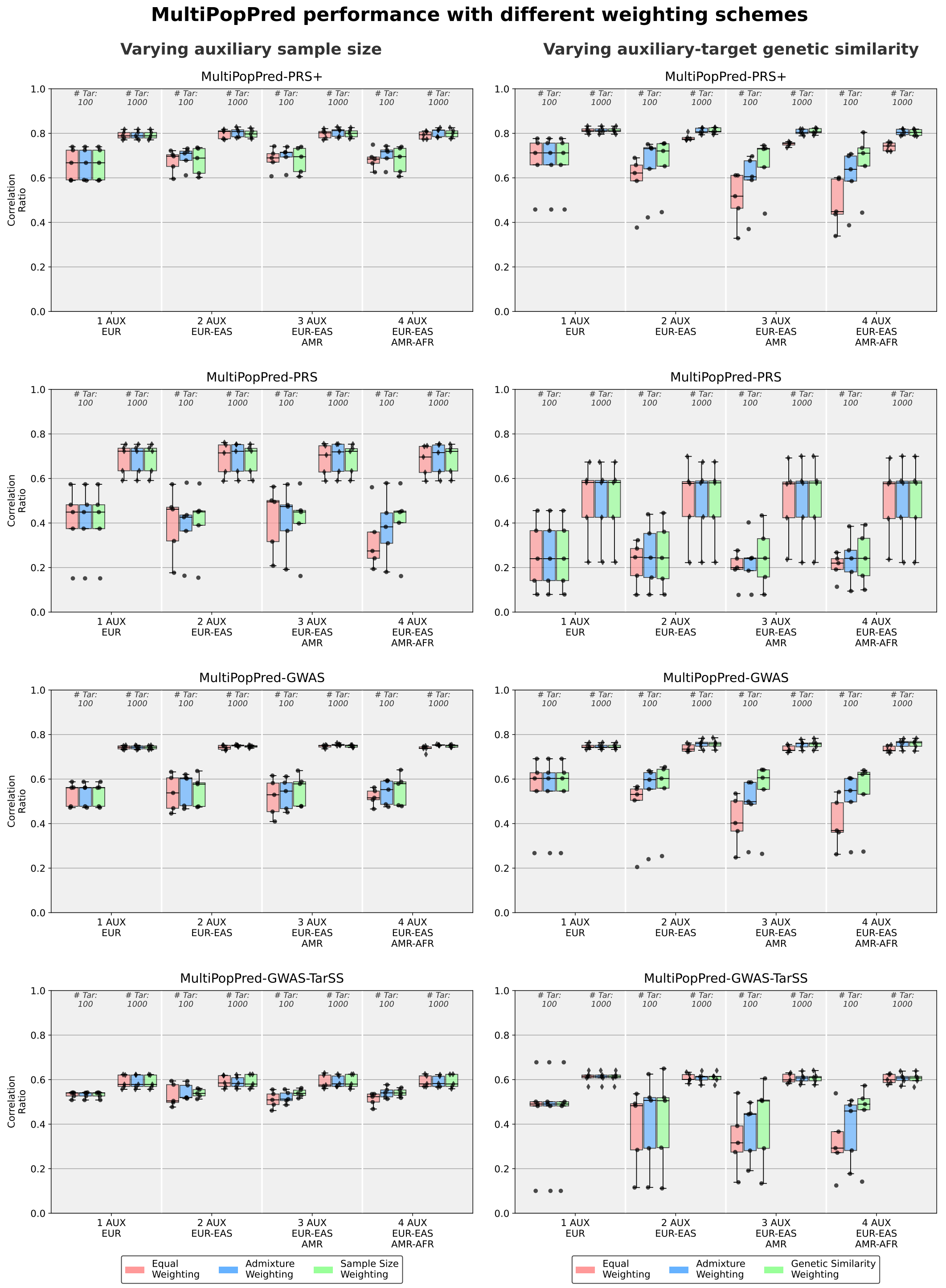

Supplementary Figure 11: Evaluating performance (Phenotype Prediction) of MultiPopPred under different weighing schemes. For ease of interpretation, readers are encouraged to focus on the settings with 4 auxiliary populations where the different weighting schemes can have the most pronounced effect, if any.

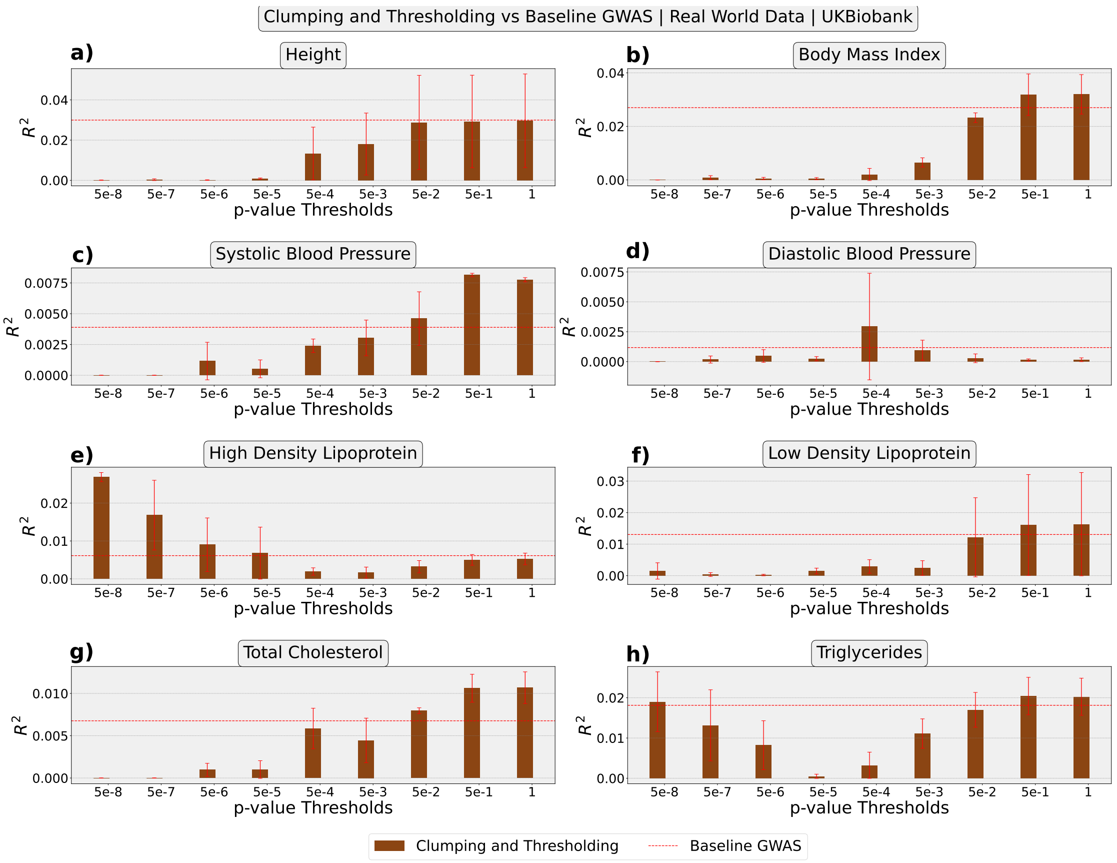

Supplementary Figure 12: Performance Comparison on Real-World Data (Phenotype Prediction): We depict the comparative performance of Baseline GWAS against a traditionally used Clumping and Thresholding PRS computation method, when applied to 8 different quantitative traits from UK Biobank.

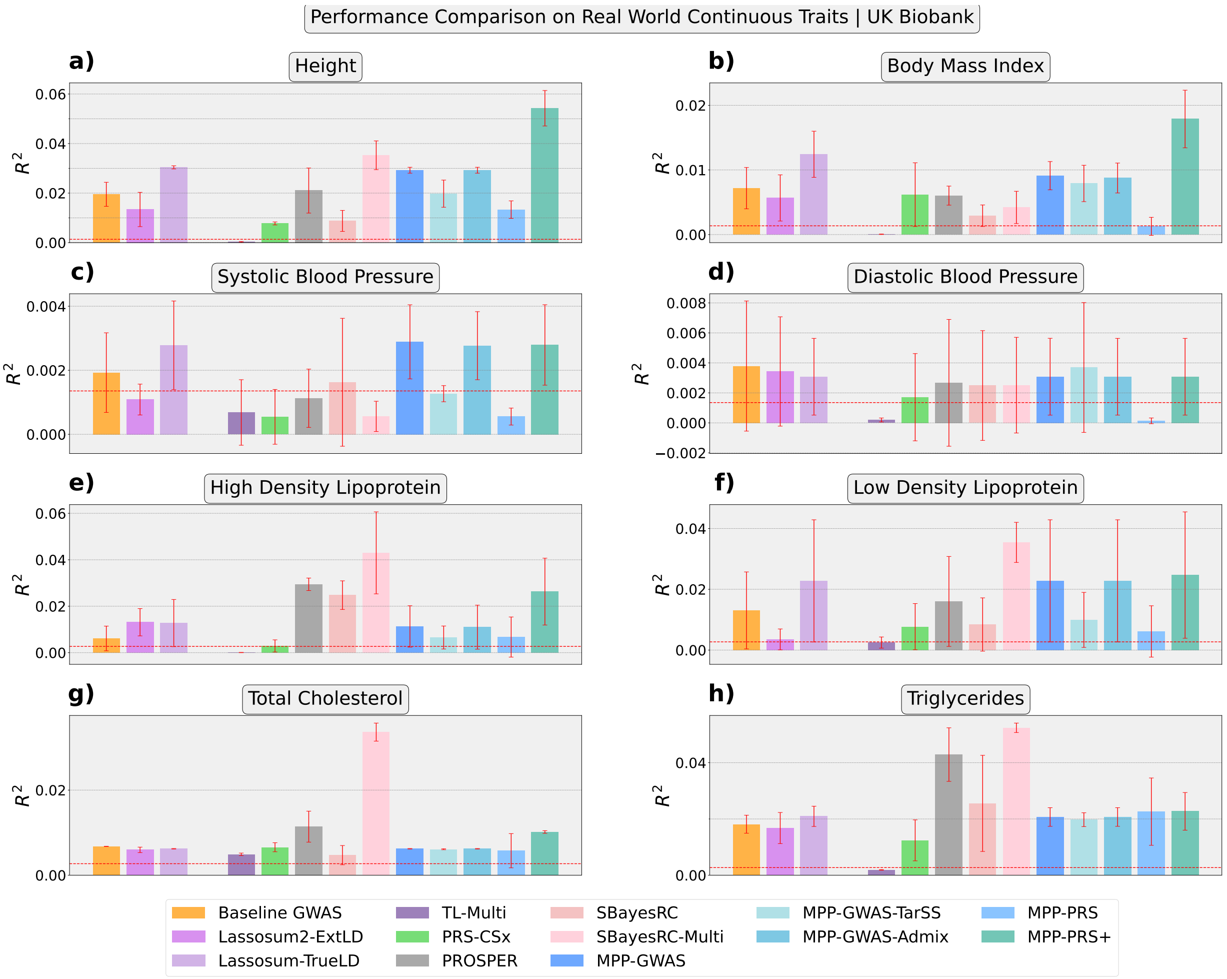

Supplementary Figure 13: Performance Comparison on Real-World Data (Phenotype Prediction): We depict the comparative performance of all MultiPopPred versions with SOTA methods when applied to 8 different quantitative traits from UK Biobank. The horizontal red line in each panel indicates the critical $R^2$ required to achieve a statistically significant (Pearson's) correlation at $\alpha$-level 0.05 between the PRS and ground-truth phenotype.

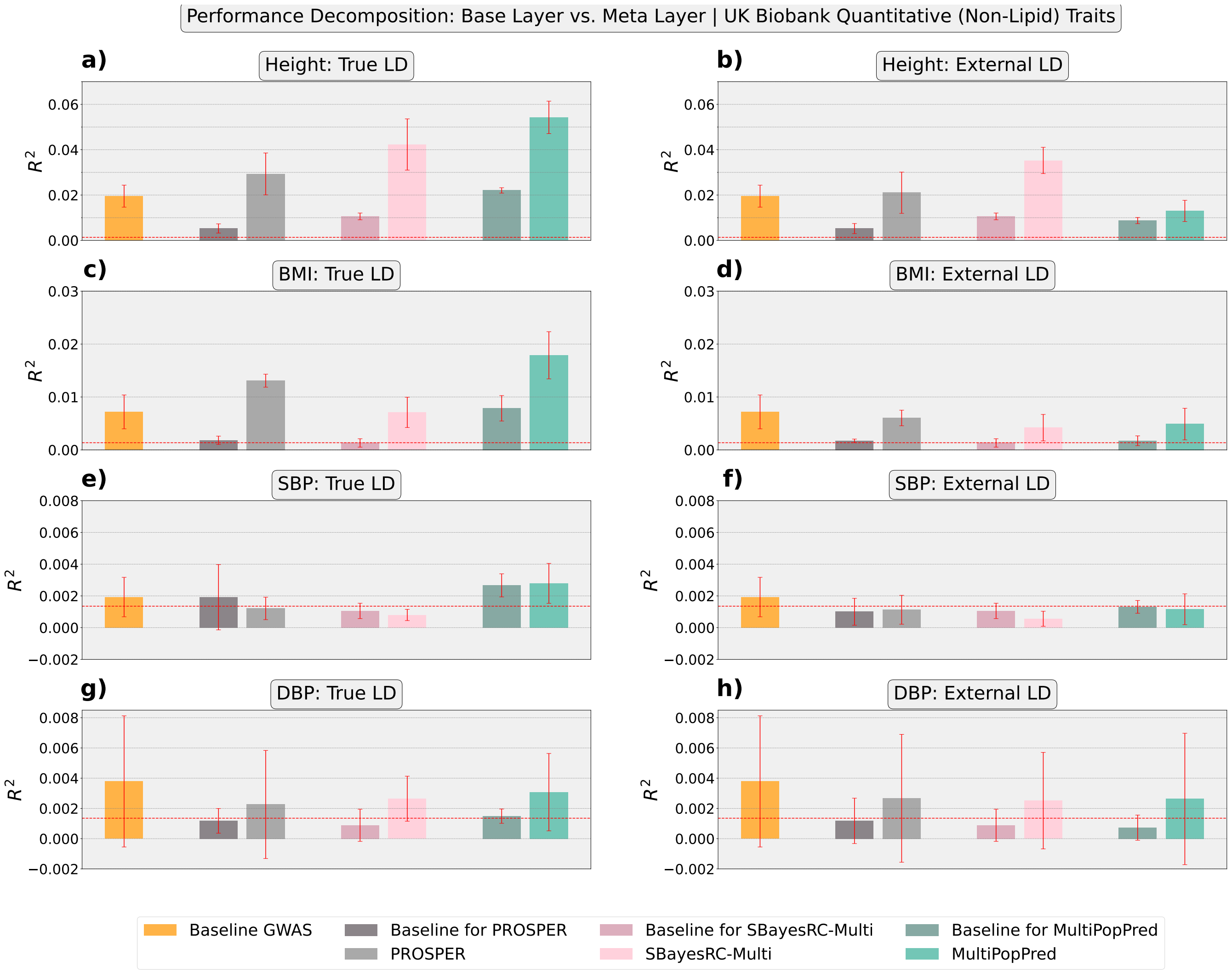

Supplementary Figure 14: Performance comparison of base layer versus meta layer of MultiPopPred, SBayesRC-Multi and PROSPER under true LD and external LD input-matched settings in the case of quantitative non-lipid traits from UK Biobank.

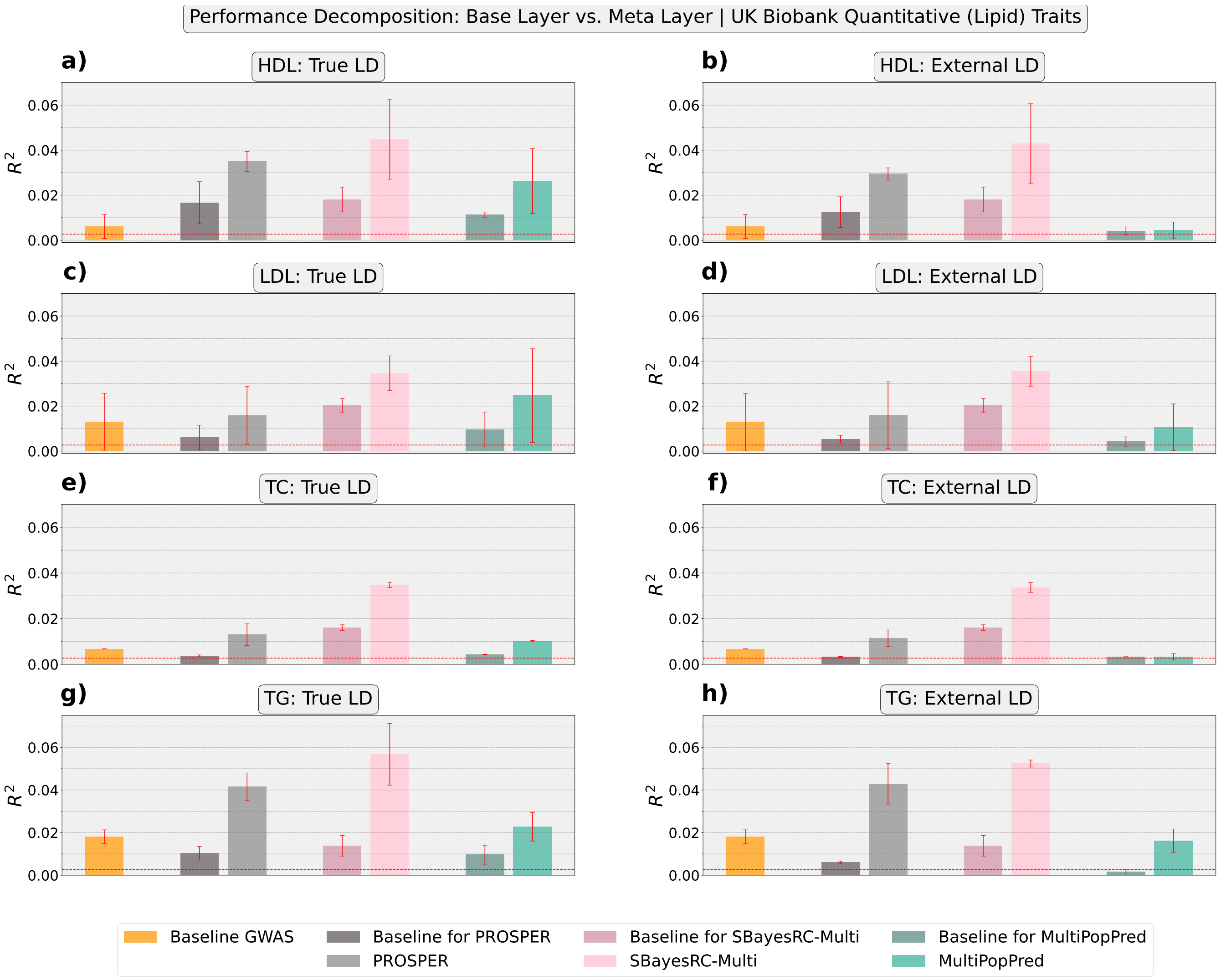

Supplementary Figure 15: Performance comparison of base layer versus meta layer of MultiPopPred, SBayesRC-Multi and PROSPER under true LD and external LD input-matched settings in the case of quantitative lipid traits from UK Biobank.

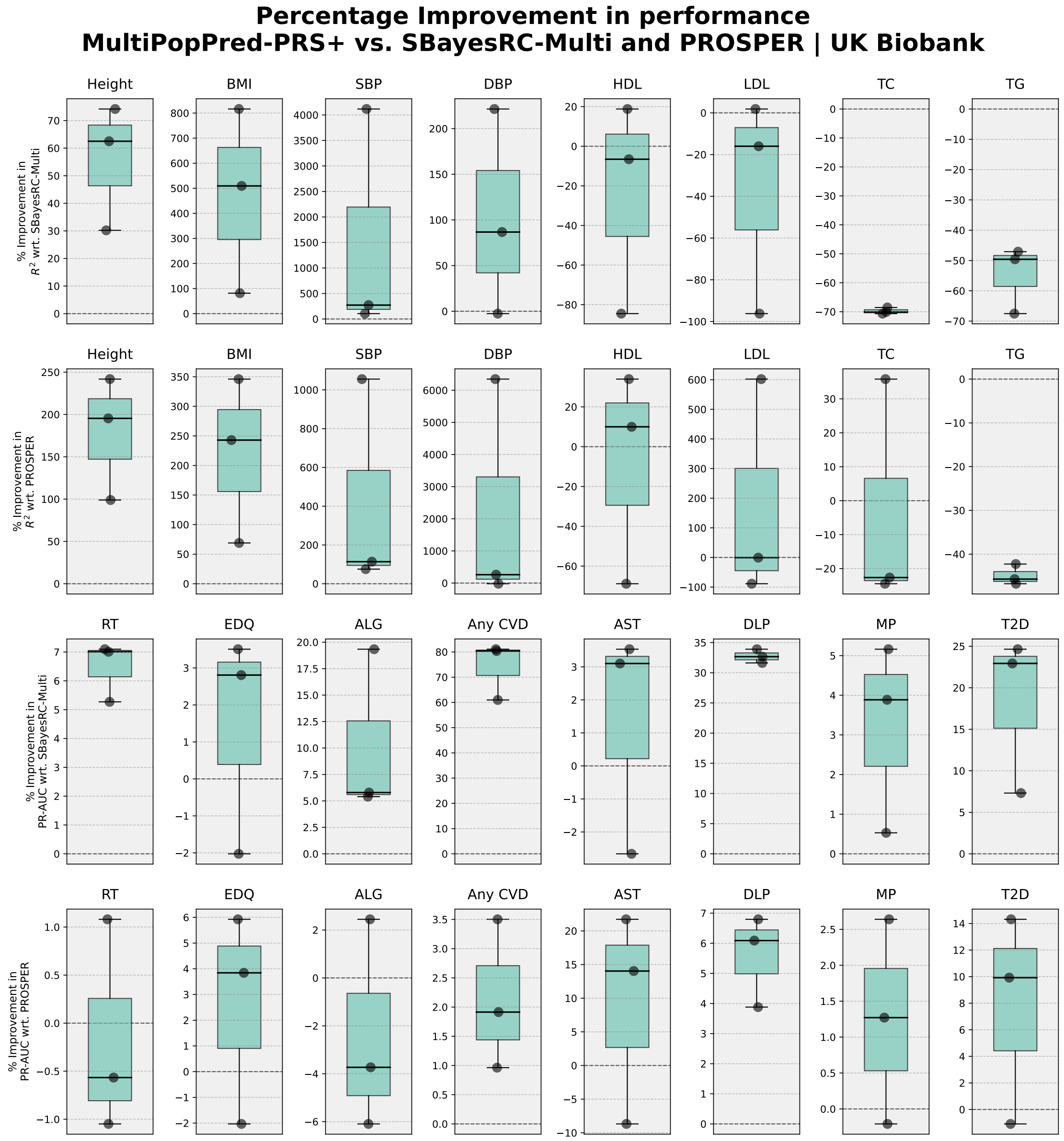

Supplementary Figure 16: Percentage improvement in performance obtained by MPP-PRS+ over SBayesRC-Multi and PROSPER with respect to 8 quantitative and 8 binary traits from UK Biobank.

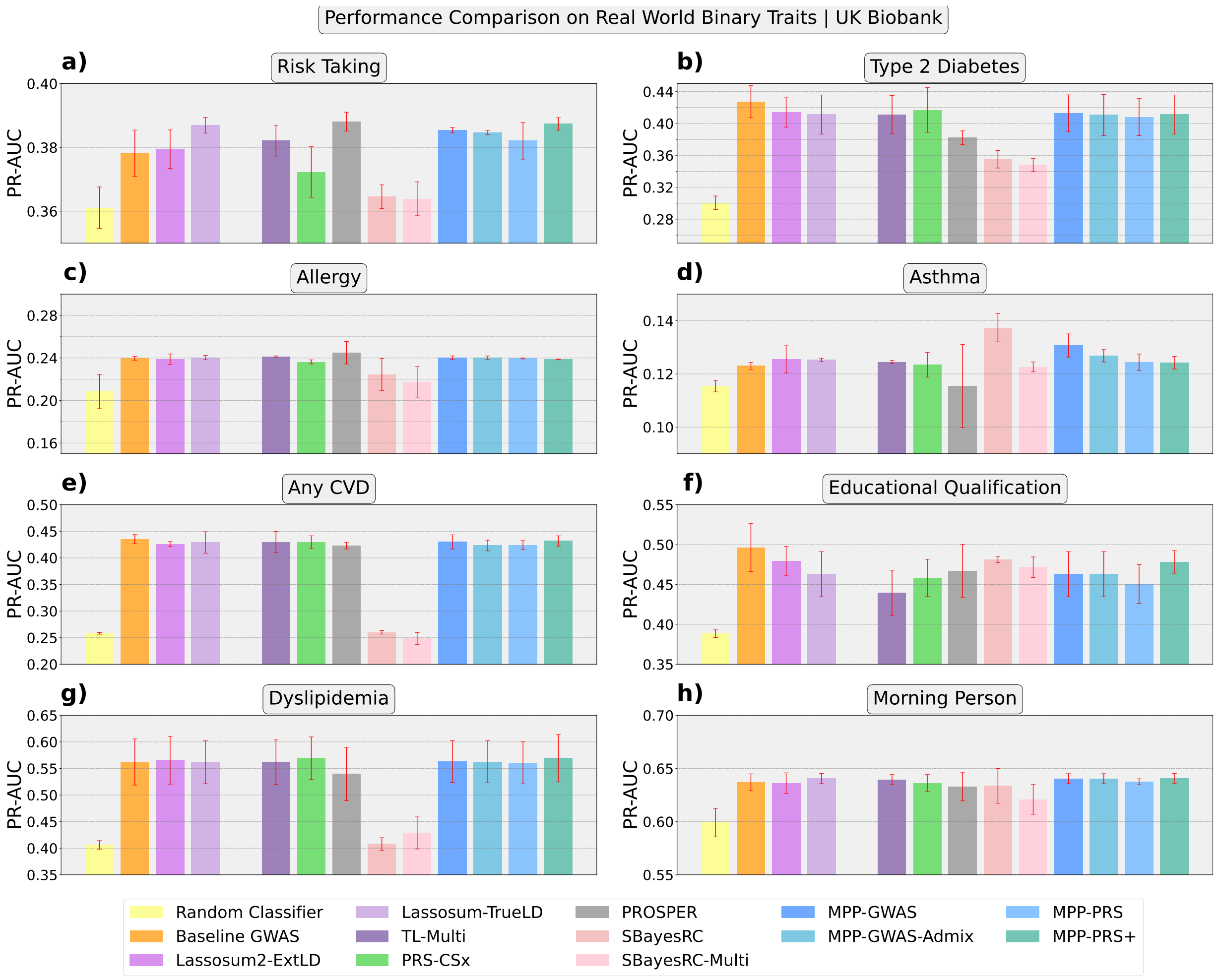

Supplementary Figure 17: Performance Comparison on Real-World Data (Phenotype Prediction): We depict the comparative performance of all MultiPopPred versions with SOTA methods when applied to 8 different binary traits from UK Biobank.

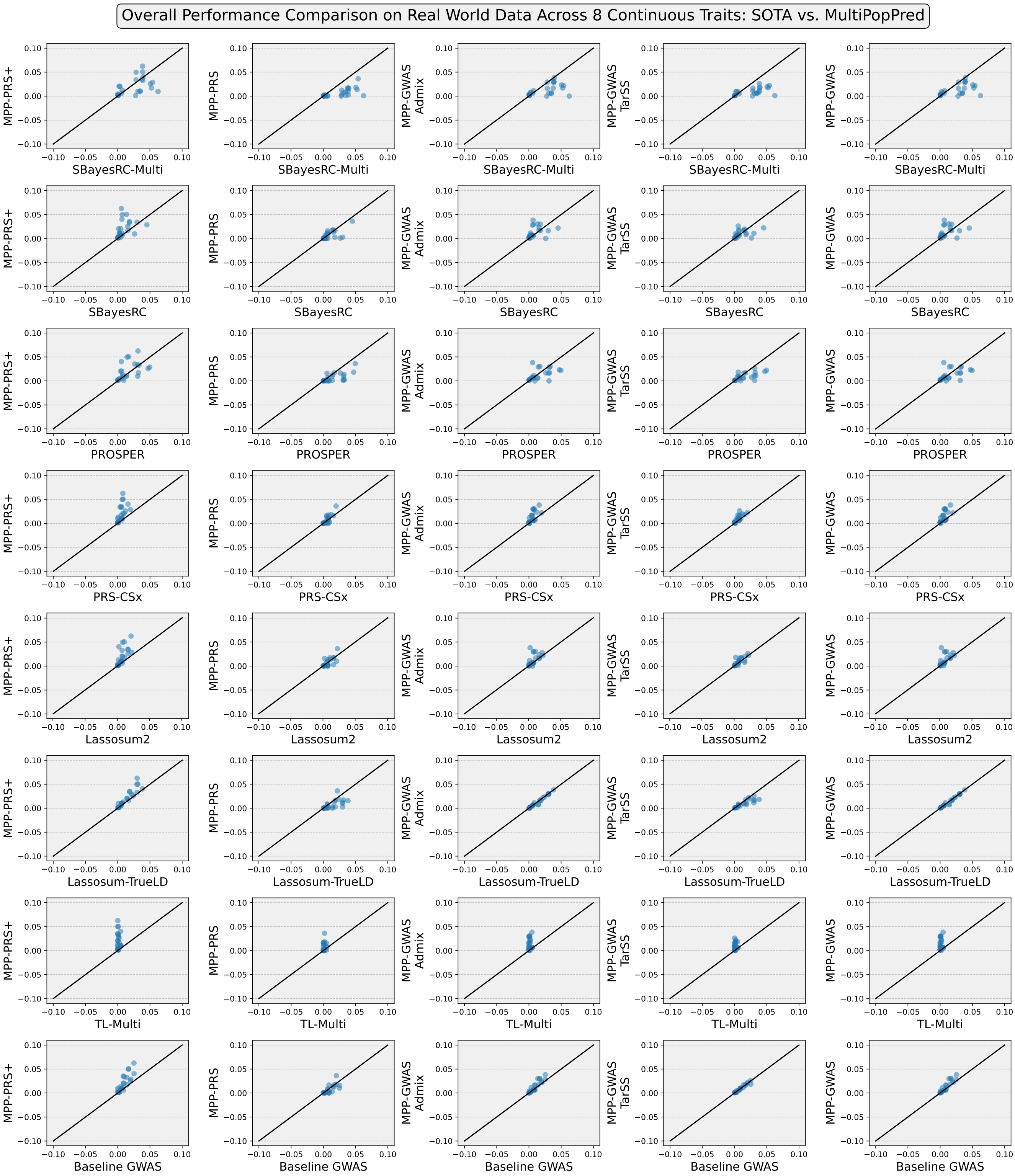

Supplementary Figure 18: Overall performance comparison on real world data across 8 continuous traits from UK Biobank.

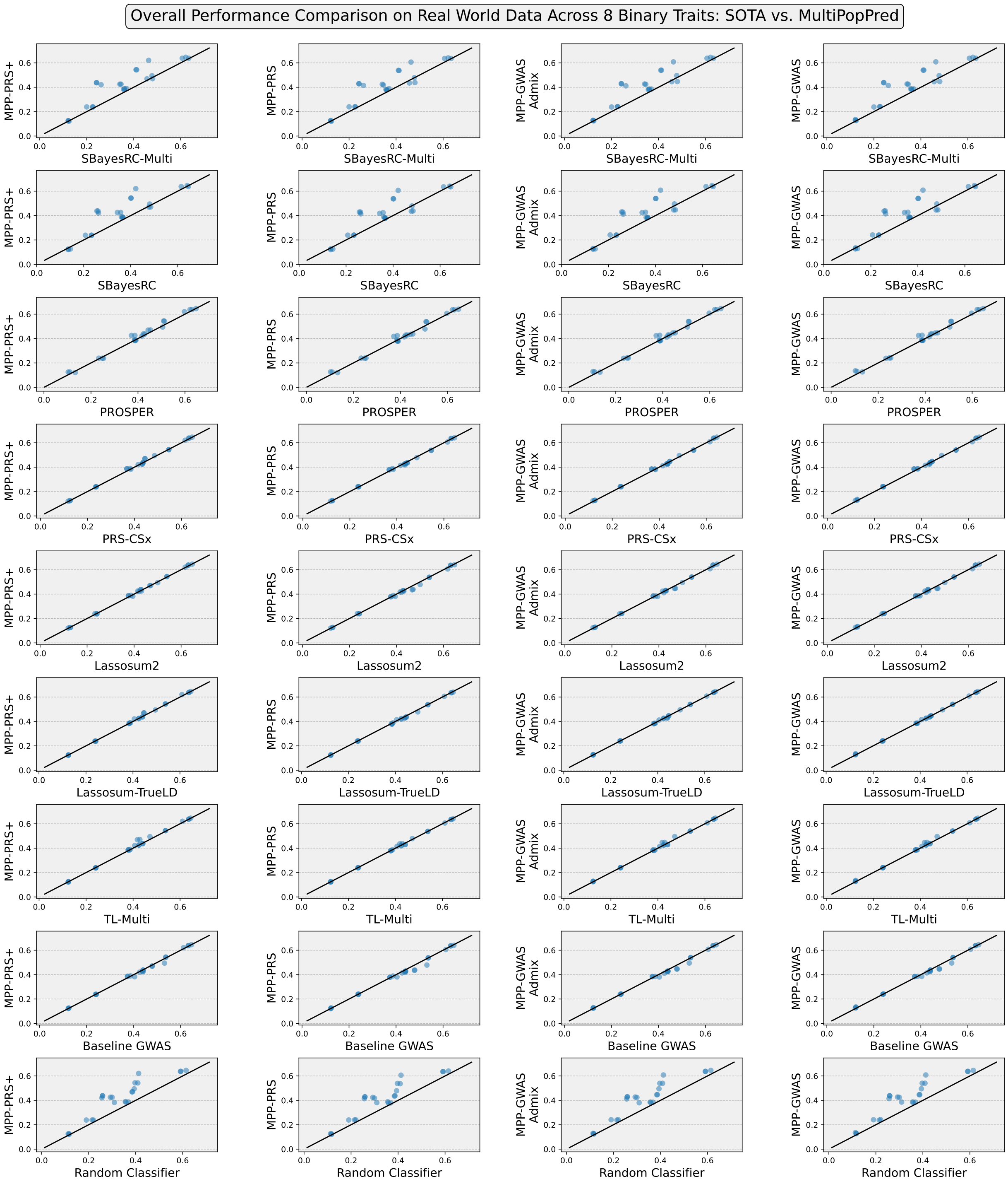

Supplementary Figure 19: Overall performance comparison on real world data across 8 binary traits from UK Biobank.

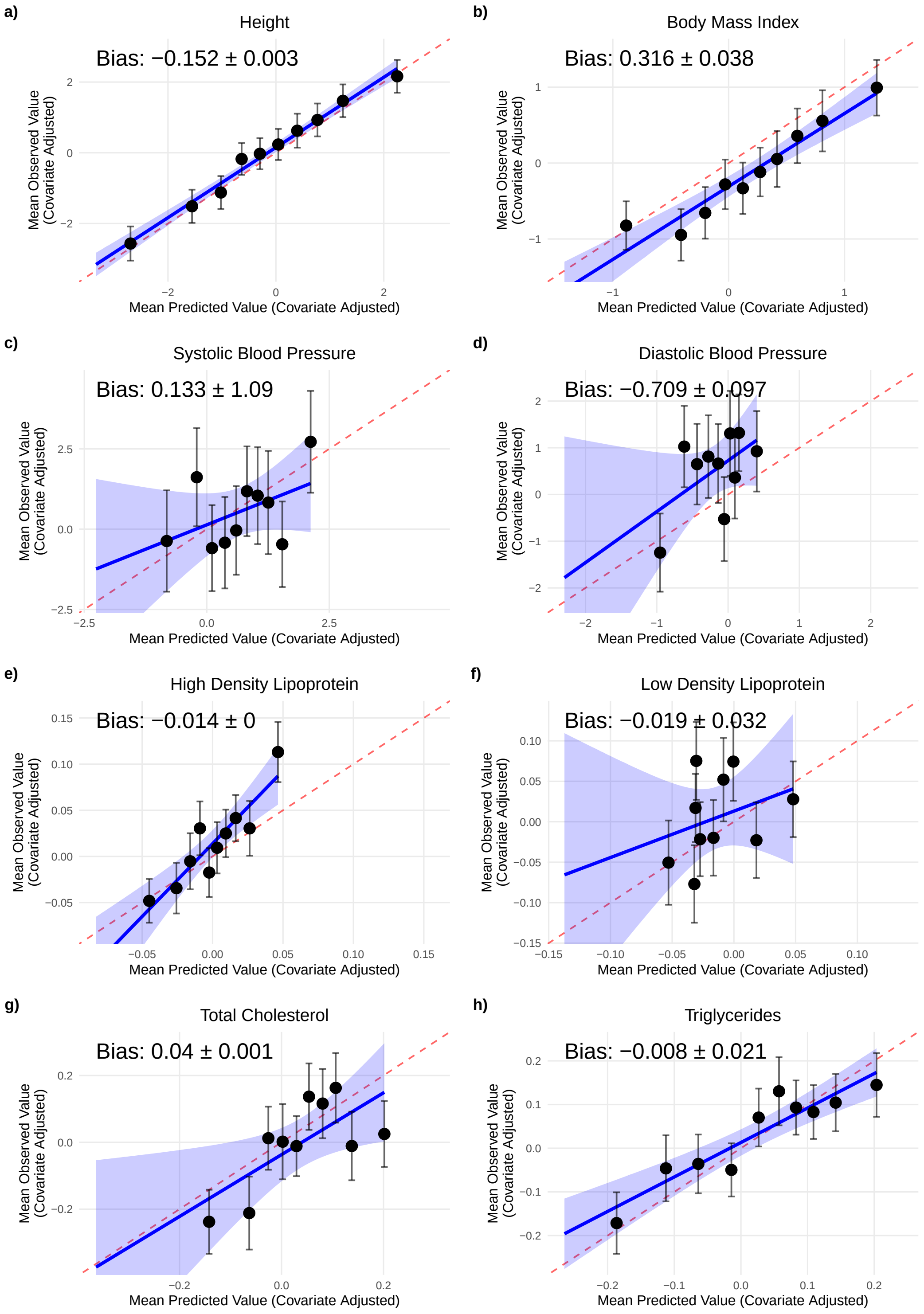

Supplementary Figure 20: Decile calibration plots based on MPP-PRS+ predictions for 8 quantitative traits from UK Biobank. Bias refers to the difference between the mean predicted phenotype and the mean ground-truth phenotype.

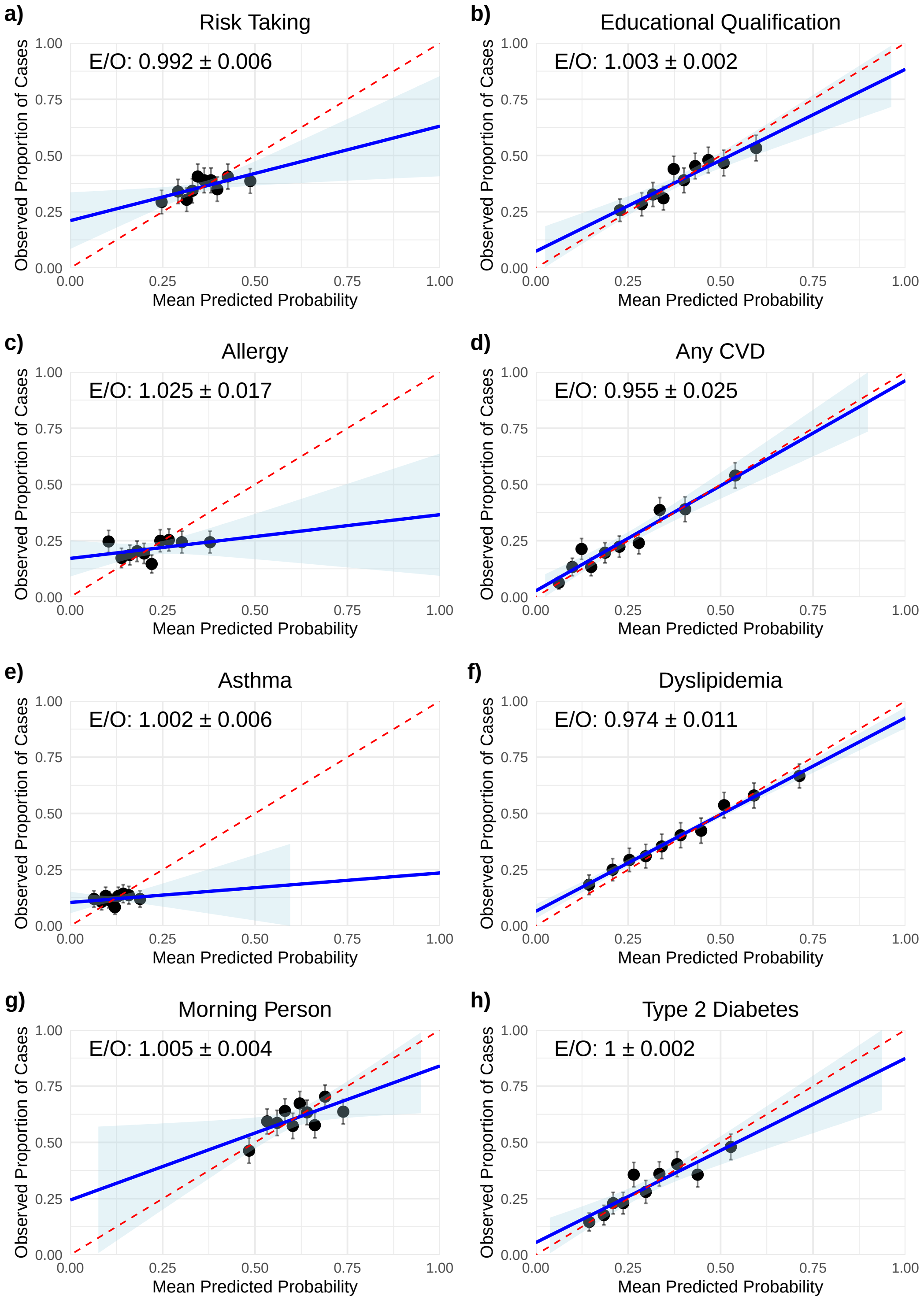

Supplementary Figure 21: Decile calibration plots based on MPP-PRS+ predictions for 8 binary traits from UK Biobank. E/O refers to the ratio of expected cases (sum of all predicted probabilities in the predicted phenotype) versus observed cases (total number of cases in the ground-truth phenotype).

### 5 Supplementary Data Files

The following Supplementary Data Files are present in different tabs within a single google sheet available at this link: `http://bit.ly/46SC61y` .

1. Supplementary Data 1: Metadata on all benchmarks used in this study: Link
2. Supplementary Data 2: Tabular details on all methods used in comparative analyses - MultiPopPred + State-of-the-art-methods: Link
3. Supplementary Data 3: MultiPopPred's optimal hyperparameters for our simulations, PROSPER's simulations, our semi-simulations, and our real-world UK Biobank analyses: Link
4. Supplementary Data 4: Our Simulations (Quantitative Traits) - All results - Varying heritability: Link
5. Supplementary Data 5: Our Simulations (Quantitative Traits) - All results - Varying number of auxiliary samples: Link
6. Supplementary Data 6: Our Simulations (Quantitative Traits) - All results - Varying number of target samples: Link
7. Supplementary Data 7: Our Simulations (Binary Traits) - All results (For each configuration, the simulation was repeated 5 times to obtain 5 replicate datasets, and the test performance metrics were reported for each of these datasets): Link
8. Supplementary Data 8: Percentage improvement in performance - All simulations and semi-simulations: Link
9. Supplementary Data 9: Our Semi-Simulations (Quantitative Traits) - All results: Link
10. Supplementary Data 10: Metadata on UK Biobank traits and samples used in our real-world analyses: Link
11. Supplementary Data 11: Metadata on UK Biobank SNPs used in our real-world analyses: Link
12. Supplementary Data 12: Real-world UK Biobank analyses - Quantitative Traits with 1000 Target population test sample size - All results: Link
13. Supplementary Data 13: Real-world UK Biobank analyses - Quantitative Traits with 2000 Target population test sample size - All results: Link
14. Supplementary Data 14: Real-world UK Biobank analyses - Binary Traits with 1000 Target population test sample size - All results: Link
15. Supplementary Data 15: Percentage improvement and Delta improvement in performance ($R^2$) obtained by MultiPopPred against all SOTA methods - All Quantitative Traits from UK Biobank: Link
16. Supplementary Data 16: Percentage improvement and Delta improvement in performance (PR-AUC) (and Percentage Improvement in terms of Logit Variance) obtained by MultiPopPred against all SOTA methods - All Binary Traits from UK Biobank: Link

17. Supplementary Data 17: Real-world UK Biobank analysis (Quantitative Traits) - All results - C&T versus Baseline GWAS: Link

18. Supplementary Data 18: Robustness Analysis: Real-world UK Biobank Quantitative Traits with true LD input to all methods - All results: Link

19. Supplementary Data 19: Robustness Analysis: Real-world UK Biobank Quantitative Traits with external LD input to all methods - All results: Link

20. Supplementary Data 20: Performance Decomposition analyses on UK Biobank Quantitative Traits (true LD and external LD) - Observing performance gains obtained from base layer versus meta layer - MultiPopPred versus PROSPER versus SBayesRC-Multi: Link

21. Supplementary Data 21: Total number of SNPs provided as inputs versus Total number of SNPs predicted by each method (after their respective internal filtering/post-processing) - MultiPopPred versus PROSPER versus SBayesRC-Multi versus PRS-CSx: Link

22. Supplementary Data 22: Real-world UK Biobank analysis (Height) - Baseline performance at varying sample sizes for European samples: Link

23. Supplementary Data 23: Comparison of computational resources required for training - MultiPopPred versus PROSPER versus SBayesRC-Multi versus PRS-CSx: Link

24. Supplementary Data 24: Comparison of MPP-PRS+ performance at different thresholds of maximum number of function evaluations as a termination criteria: Link

25. Supplementary Data (Plots) 25: Manhattan Plots and QQ Plots for GWAS conducted on all real-world quantitative and binary traits from UK Biobank: Link

[3] Pauli Virtanen, Ralf Gommers, Travis E. Oliphant, Matt Haberland, Tyler Reddy, David Cournapeau, Evgeni Burovski, Pearu Peterson, Warren Weckesser, Jonathan Bright, Stéfan J. van der Walt, Matthew Brett, Joshua Wilson, K. Jarrod Millman, Nikolay Mayorov, Andrew R. J. Nelson, Eric Jones, Robert Kern, Eric Larson, C J Carey, İlhan Polat, Yu Feng, Eric W. Moore, Jake VanderPlas, Denis Laxalde, Josef Perktold, Robert Cimrman, Ian Henriksen, E. A. Quintero, Charles R. Harris, Anne M. Archibald, Antônio H. Ribeiro, Fabian Pedregosa, Paul van Mulbregt, and SciPy 1.0 Contributors. SciPy 1.0: Fundamental Algorithms for Scientific Computing in Python. *Nature Methods*, 17:261–272, 2020.
